# Mg^2+^-dependent mechanism of environmental versatility in a multidrug efflux pump

**DOI:** 10.1101/2024.06.10.597921

**Authors:** Benjamin Russell Lewis, Muhammad R. Uddin, Katie M. Kuo, Laila M. N. Shah, Nicola J. Harris, Paula J. Booth, Dietmar Hammerschmid, James C. Gumbart, Helen I. Zgurskaya, Eamonn Reading

## Abstract

Tripartite resistance nodulation and cell division multidrug efflux pumps span the periplasm and are a major driver of multidrug resistance among Gram-negative bacteria. The periplasm provides a distinct environment between the inner and outer membranes of Gram-negative bacteria. Cations, such as Mg^2+^, become concentrated within the periplasm and, in contrast to the cytoplasm, its pH is sensitive to conditions outside the cell. Here, we reveal an interplay between Mg^2+^ and pH in modulating the dynamics of the periplasmic adaptor protein, AcrA, and its function within the prototypical AcrAB-TolC multidrug efflux pump from *Escherichia coli*. In the absence of Mg^2+^, AcrA becomes increasingly plastic within acidic conditions, but when Mg^2+^ is bound this is ameliorated, resulting in domain specific organisation in neutral to weakly acidic regimes. We establish a unique histidine residue directs these structural dynamics and is essential for sustaining pump efflux activity across acidic, neutral, and alkaline conditions. Overall, we propose Mg^2+^ conserves the structural mobility of AcrA to ensure optimal AcrAB-TolC function within rapid changing environments commonly faced by the periplasm during bacterial infection and colonization. This work highlights that Mg^2+^ is an important mechanistic component in this pump class and possibly across other periplasmic lipoproteins.

## Introduction

Antimicrobial resistance (AMR) is a major threat to global health, and is expected to reduce life expectancy globally by 1.8 years over the next decade without proper action.^1^ The Resistance Nodulation and cell Division (RND) efflux pump protein superfamily significantly contributes to multidrug resistance in Gram-negative bacteria, due to their ability to export an extensive range of chemically diverse molecules and reduce intracellular concentrations of antibitoics.^2–4^ RND efflux pumps are tripartite assemblies that span the double membrane and periplasm of Gram-negative bacteria, and AcrAB-TolC, from *E. coli*, is the classical RND efflux pump.^5–7^ It consists of an inner membrane transporter AcrB - that is energized by the proton motive force - connected to an outer membrane factor (OMF) TolC by the periplasmic adaptor protein (PAP) AcrA, also known as the membrane fusion protein (MFP), forming a 3:6:3 (AcrB:AcrA:TolC) complex stoichiometry (**Figure 1A**).^8,9^ For efficient substrate efflux to occur, AcrA must have the structural mobility capable of transmitting conformational information between AcrB and TolC during substrate recognition and transport, whilst also possessing the stability needed for maintaining a continuous sealed channel.^10^ With homologs across other Gram-negative ESKAPE bacteria, enhanced mechanistic insight into this system can inform on wider RND behaviour as well as emerging efforts to inhibit the protein class to restore antibiotic efficacy.^11–13^

**Figure 1.**
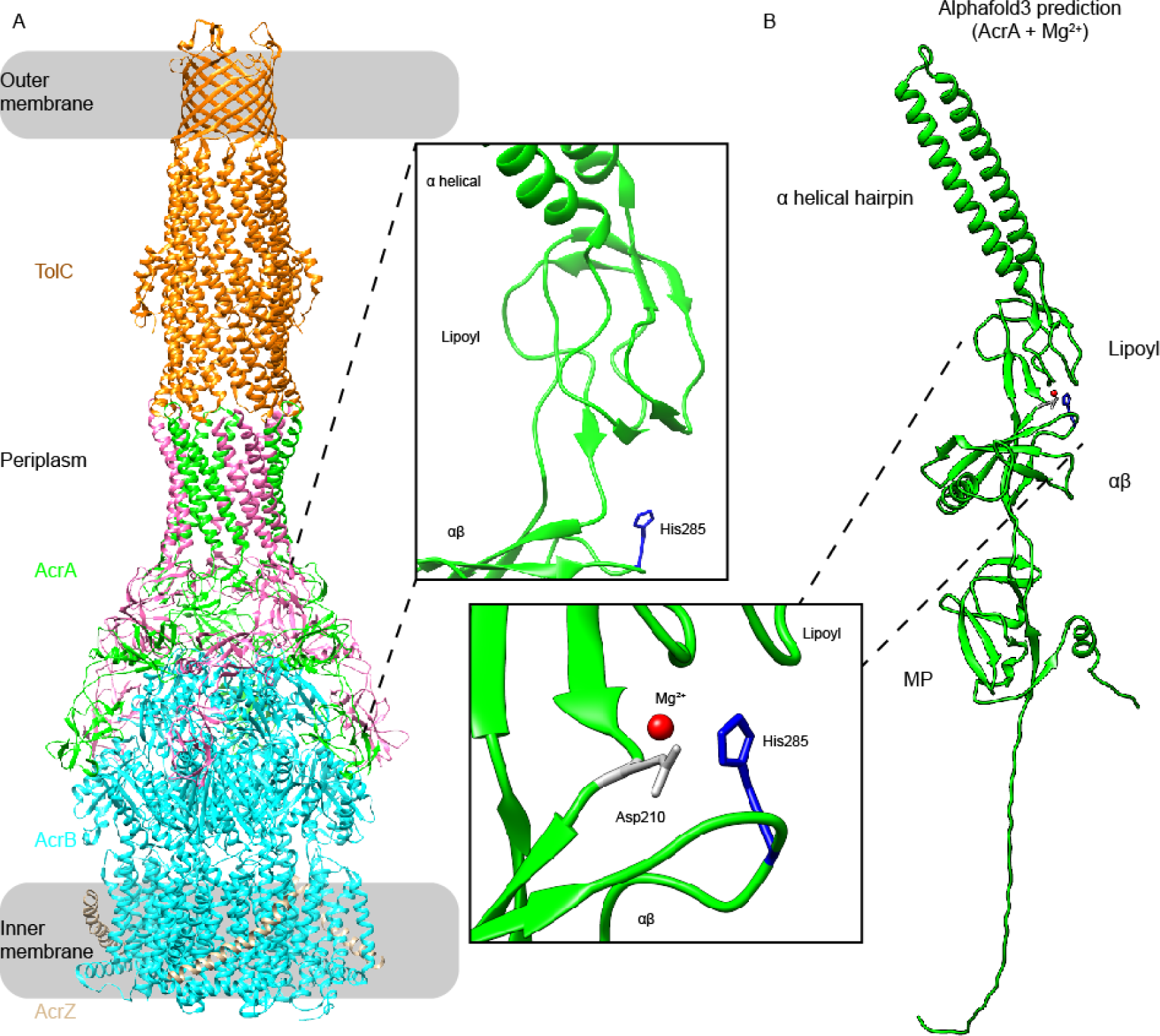
His285 and likely Mg^2+^ position within AcrA from the Escherichia coli AcrAB(Z)-TolC multidrug efflux. **A.** Structure of AcrAB(Z)-TolC in the E. coli Gram-negative cell envelope. Each dimer protomer in the AcrA trimer of dimers is coloured either green or red. An insert shows the location of His285 (blue), previously suggested to be a conformational switch in AcrA.^16,21^ **B.** AlphaFold3 prediction of the AcrA and Mg^2+^ complex. The best scoring model saw Mg^2+^ bind to Asp210, located between the lipoyl and αβ barrel domains and in close proximity to His285. The reported predicted template modelling (pTM) score and the interface predicted template modelling (ipTM) score were 0.59 and 0.73, respectively, and had a ranking score of 0.77. MP = membrane proximal domain.

The periplasmic environment AcrA resides in is distinct compared to the internal cytosolic and external environments of Gram-negative bacteria; it has a pH closer to the outside of the cell, concentrated cation populations, such as Mg^2+^ (driven by the Donnan potential), and a peptidoglycan layer.^14^ Critically, understanding collective contributions of protons and Mg^2+^ becomes important when considering their putative fluctuations within the periplasm during the enteric journey of *E. coli* through a human gut (e.g. stomach pH 1.5-4.0 and intestines pH 4.0-7.5) and within the microenvironments of macrophage phagosomes.^15–19^ Previous investigations have shown that AcrA conformation is pH- and Mg^2+^-sensitive^16,20,21^ and that related heavy metal efflux RND (HME-RND) pump systems - which export metals as part of their function, i.e. Zn^2+^ by ZneCAB, and Cu^+^ and Ag^+^ by CusCBA - can interact with the divalent metal cations they export to induce conformational changes.^22,23^ However, it remains uncertain whether Mg^2+^ has a directive role within RND multidrug efflux pump structure and function.

Here, through a combination of hydrogen/deuterium exchange mass spectrometry (HDX-MS), molecular dynamics (MD) simulations, and biophysical investigations we uncover Mg^2+^ as a structural cofactor within the periplasmic adaptor protein AcrA. We show that Mg^2+^ transforms the dynamics of AcrA when exposed to increased acidity but has little influence at neutral pH, and that without Mg^2+^ acidification leads to backbone destabilisation throughout the protein. In contrast, when Mg^2+^ is present then acidification causes domain-localised backbone stability. Mg^2+^, therefore, directs the route pH-induced conformational alterations can take. This discovery harmonizes previous, seemingly conflicting, computational models, which found that protonation of a unique His285 within AcrA caused both extensive destabilisation and localised stabilisation of backbone hydrogen-bonding networks.^16^ Using bacterial efflux assays, we go further to substantiate the importance of both Mg^2+^ and His285 for AcrA regulation, revealing that His285 is essential in ensuring AcrAB-TolC functions across a broad pH range (pH 5-8). Taken together, this work supports that AcrA recruits available protons and Mg^2+^ ions to ensure continued AcrAB-TolC function and bacterial survival, a process which may be general across related periplasmic protein adaptor proteins also involved in multidrug efflux resistance.

## Results & Discussion

Motivated by the heightened concentration of protons and Mg^2+^ AcrA experiences within the *E. coli* periplasm, we explored the effect, and interplay, of Mg^2+^ and pH on its structure-function. To simulate acidic and neutral periplasmic pH conditions *in vitro* we investigated AcrA at pH 6.0 and 7.4, with and without the addition of 1 mM MgCl_2_, with bacterial functional studies performed across a broader pH range of 5.0 to 8.0. Wild type, lipidated AcrA, was used for all functional assays and a previously established functional and soluble AcrA construct (AcrA^s^), with its lipidation site removed by mutation, was used for structural and biophysical investigations as it provided the required sample homogeneity.^12,20^ This delipidated AcrA construct was also utilized for MD simulations, with (de)protonation of the unique His285 residue used as a proxy for pH 7.4 and 6.0 conditions. His285 is located at the αβ barrel domain surface facing the lipoyl domain and is the only residue within AcrA that has a titratable protonation state change between mildly acidic and neutral regimes (with a pKa ∼6) (**Figure 1A**).

### Magnesium is a possible AcrA cofactor

Mg^2+^ binding to proteins has been well considered.^24^ As a hard metal, Mg^2+^ is prone to bind oxygens from the side chain carboxylate groups in Glu and Asp residues (these residues in AcrA are highlighted in **Figure S1**), with several classes of recognised Mg^2+^ cofactor sites that share an abundance of Glu and Asp residues found across diverse classes of enzymes.^24^ To investigate whether AcrA could bind metal cations we first used *Me*Bi*Pred* – which uses machine learning algorithms to determine metal binding based on a protein primary sequence^25^ – to explore binding to a variety of mono/di/trivalent metal cations, including Ca, Co, Cu, Fe, K, Mg, Mn, Na, Ni, and Zn. Only Mg^2+^ was identified as a binding metal (**Table 1**). Expansion of the *Me*Bi*Pred* analysis predicts Mg^2+^ binding to related, lipidated RND periplasmic adaptor proteins, indicating Mg^2+^ binding may be generic across this type. Indeed, when performing an analysis on the subcellular proteomes within the periplasmic space of *Escherichia coli* K-12^26,27^ (see **Methods**), inner membrane lipoproteins (IMLPs) had the highest Mg^2+^ binding proportion predicted at 43.5%, compared to outer membrane lipoproteins (OMLPs, 21.6%), periplasmic proteins peripherally associated with IM (IM-peri, 29.2%) and soluble proteins located in the periplasm (periplasm, 23.6%) (**Figure S2**).

**Table 1.**
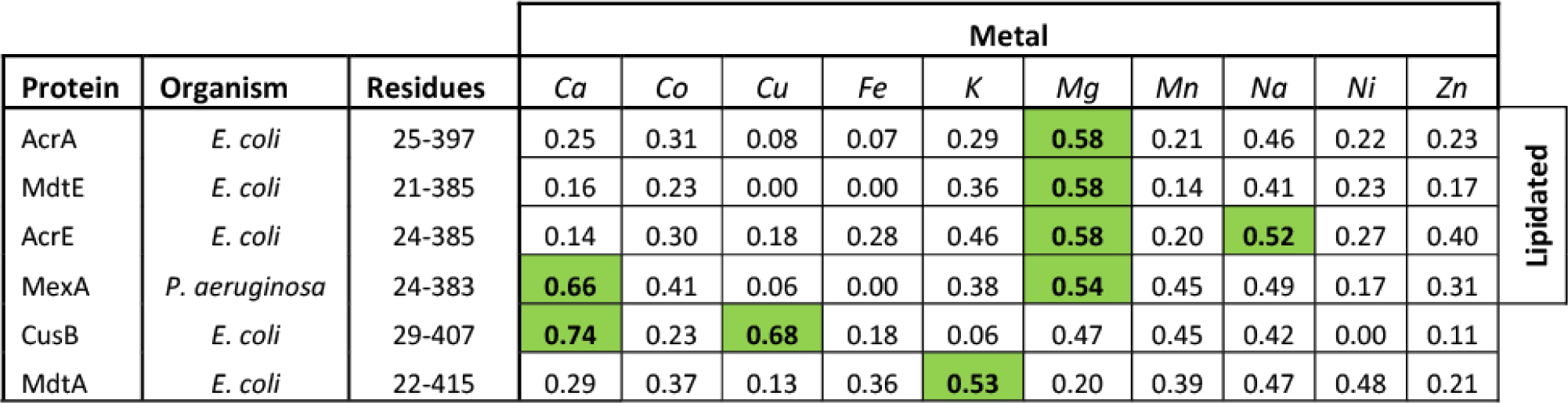
Predictions of metal binding in AcrA, and related PAPs. The *mebipred* (v2.0) program was used to identify metal-binding potential of protein sequences. Protein sequences that exceeded the cutoff of 0.5 (coloured green) are considered an indication of metal-binding to either Fe, Ca, Na, K, Mg, Mn, Cu, K, Co, Ni metals as defined in Aptekmann *et al.*^25^

| Protein | Organism | Residues | Metal |  |  |  |  |  |  |  |  |  |  |
| --- | --- | --- | --- | --- | --- | --- | --- | --- | --- | --- | --- | --- | --- |
|  |  |  | Ca | Co | Cu | Fe | K | Mg | Mn | Na | Ni | Zn |  |
| AcrA | *E. coli* | 25-397 | 0.25 | 0.31 | 0.08 | 0.07 | 0.29 | 0.58 | 0.21 | 0.46 | 0.22 | 0.23 | Lipidated |
| MdtE | *E. coli* | 21-385 | 0.16 | 0.23 | 0.00 | 0.00 | 0.36 | 0.58 | 0.14 | 0.41 | 0.23 | 0.17 |  |
| AcrE | *E. coli* | 24-385 | 0.14 | 0.30 | 0.18 | 0.28 | 0.46 | 0.58 | 0.20 | 0.52 | 0.27 | 0.40 |  |
| MexA | *P. aeruginosa* | 24-383 | 0.66 | 0.41 | 0.06 | 0.00 | 0.38 | 0.54 | 0.45 | 0.49 | 0.17 | 0.31 |  |
| CusB | *E. coli* | 29-407 | 0.74 | 0.23 | 0.68 | 0.18 | 0.06 | 0.47 | 0.45 | 0.42 | 0.00 | 0.11 |  |
| MdtA | *E. coli* | 22-415 | 0.29 | 0.37 | 0.13 | 0.36 | 0.53 | 0.20 | 0.39 | 0.47 | 0.48 | 0.21 |  |

Building upon this further, AlphaFold3 was applied to AcrA and Mg^2+^ (**Figure 1B**). AlphaFold3 is capable of joint structure predictions of protein complexes with DNA/RNA, ligands and ions.^28^ It provides both a predicted template modelling (pTM) score and an interface predicted template modelling score (ipTM) as measures of accuracy for its ligand-protein structural interaction predictions. For AcrA and Mg^2+^, the best scoring model (pTM = 0.59, ipTM = 0.73, and ranking score = 0.77) showed Mg^2+^ bound to Asp210, which is between the lipoyl and αβ barrel domains of AcrA and in close proximity to His285 (**Figure 1B**). Comparable scores were seen for ZneB and Zn^2+^, a confirmed zinc binder, and other homologous PAPs were predicted to bind Mg^2+^ in similar areas as seen for AcrA (**Figure S3**).

Next, we used isothermal titration calorimetry (ITC) to verify AcrA binding to Mg^2+^ and found that Mg^2+^ binds to AcrA similarly, within the low mM range, at both pH 6.0 and pH 7.4 (**Figure S4C-D**). Native MS analysis supports the occurrence of weak, and possibly multivalent, Mg^2+^ binding to AcrA^S^ without modifying its oligomeric state (**Figure S4E**). When incubated with 100 μM MgCl_2_ increased peak widths were observed, characteristic of multiple Mg^2+^ ions remaining bound to the protein in the gas phase. Although, resolution of the quadrupole time-of-flight (Q-ToF) mass spectrometer used prevented exact stoichiometries being assigned. Circular dichroism (CD) spectroscopy was then used to evaluate secondary structure and thermal stability consequences of Mg^2+^ binding to AcrA.^29^ Secondary structure content was indistinguishable in the presence or absence of MgCl_2_ within the pH 7.4 condition, whereas at pH 6.0 MgCl_2_ caused an increase in β-sheet content (**Figure S4F** and **Table S1**).^30^ Interestingly, the related HME-RND periplasmic adaptor protein, ZneB, had an increase in β-sheet content when bound to its Zn^2+^ cofactor.^22^ We also performed CD thermal melts on AcrA^S^ ± Mg^2+^, which revealed Mg^2+^ binding did not significantly affect the thermal stability of AcrA^S^ at either pH 6.0 or 7.4 (**Figure S4G**).

To better understand how AcrA is interacting with Mg^2+^, we performed MD simulations on AcrA ± MgCl_2_ or NaCl and monitored the occupancy of the ions across the different domains of AcrA^S^ (**Figure S5**). Binding was observed for both Na^+^ and Mg^2+^, and although Mg^2+^ binding was found more localised to the same lipoyl domain site found by AlphaFold3, the simulations suggested that there are likely several other regions of AcrA that can interact with Mg^2+^. Furthermore, cation binding site positions were not significantly affected by protonation of His285. Our previous MD simulations revealed free AcrA exhibited a range of orientations but had two main conformational basins^31^: one was a *cis*-like formation where the MP and α-helical domains point in the same direction, and another was a *trans*-like conformation where they point in opposite directions. Cryo-EM structures of assembled AcrAB-TolC show AcrA to be in the *trans* conformation^32,33^, and locking AcrA in the *cis* conformation was proposed to compromise the assembly of the complex. Therefore, we analysed both *cis*- and *trans*- populations of our simulations but found no statistical significance between either population, suggesting that neither Mg^2+^ or Na^+^ binding can bias, or enforce, either state (**Table S2**).

Altogether, this data supports Mg^2+^ is capable of binding AcrA in both neutral and acidic regimes but was only seen to affect structure within the latter condition. Its weak binding affinity may suggest a reversible mode of interaction across multiple areas within the structure. Intriguingly, considering that the *E. coli* periplasmic concentration of Mg^2+^ can be ∼7 times higher than in the surrounding medium, when bacteria are within environments containing 0.03-10 mM MgCl_2_, the periplasmic Mg^2+^ concentration will be in the range of 0.21-70 mM.^34^ Mg^2+^ may therefore be recruited as a cofactor to AcrA to different degrees depending on the composition of the bacterial surroundings.

### Mg^2+^ binding counteracts acid-triggered dynamics

To understand the suggested acid-selective influence of Mg^2+^ on AcrA structure further we used HDX-MS to define its backbone structural dynamics at pH 6.0 and pH 7.4, with and without MgCl_2_. For HDX-MS a protein sample is made accessible to D_2_O solvent, with HDX initiated when backbone amide hydrogens are broken through structure fluctuations and unfolding. HDX is fast within unfolded and solvent-exposed regions and slow within protected and stably folded regions. Differential HDX-MS (ΔHDX) analysis was performed to localize D-label differences between two states at peptide-level structural resolution (**Figure 2**).^35^ When comparing results collected at pH 6.0 to those at pH 7.4 all conditions were kept the same except the pH of the HDX buffers, all samples were quenched to the same final pH of 2.4, and labelling times were corrected according to Equation 3 (**see Methods**).^36^

**Figure 2.**
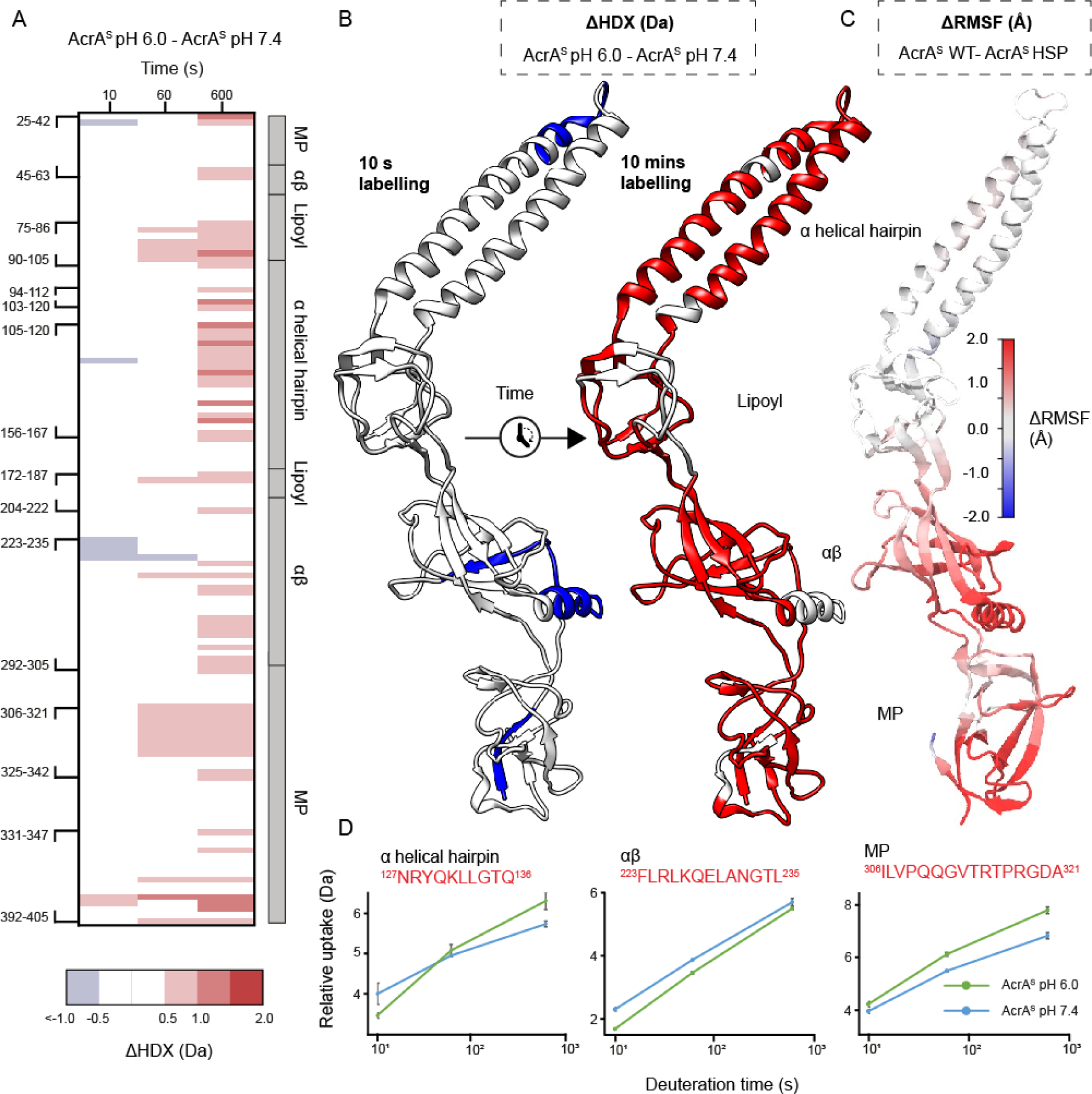
Structural dynamics of AcrA in the absence of MgCl_2_ at pH 6.0 versus pH 7.4. **A.** Chiclet plot displaying the differential HDX (ΔHDX) plots for AcrA^S^ pH 6.0 - AcrA^S^ pH 7.4. Blue signifies areas with decreased HDX between states and red signifies areas with increases HDX. Significance was defined to be ≥ 0.5 Da change with a P-value ≤ 0.01 in a Welch’s t-test (n=4). White areas represent regions with insignificant ΔHDX. **B.** ΔHDX extent (increased and decreased HDX being red and blue, respectively) for the earliest and latest time points are coloured onto the AcrA structure (PDB:5O66) using HDeXplosion and Chimera.^44,45^ **C.** AcrA coloured according to the difference in root-mean-square fluctuations (RMSF) between simulations of the AcrA^S^ wild-type state and HSP (doubly protonated His285) state (AcrA^S^ HSP -AcrA^S^ WT), averaged over four replicas for each. Red indicates that the RMSF has increased whereas blue indicates it has decreased. RMSF was calculated over the last 70 ns of each 100-ns simulation. **D**. D-label uptake plots for three select peptides in different domains of AcrA. Uptake plots are the average deuterium uptake and error bars indicate the standard deviation (n=4).

In the absence of Mg^2+^, acidification increased the backbone HDX throughout AcrA^S^, most prominently observed progressively within the latest time incubations. The earliest time point shows less difference as well as localised stabilisation within the αβ barrel domain, which is lost at the later time points. This supports that the overall protein fold is not significantly influenced, as global structure loss would result in considerable exposure of backbone amides to HDX at the earliest time point.^37^ Instead, it suggests an expanded temporal structural plasticity within more acidic conditions is occurring (**Figure 2**).

To further understand the effect of increased acidity on AcrA backbone dynamics in the absence of magnesium, differential root-mean-square fluctuations (ΔRMSF) between AcrA with His285 doubly protonated (resembling pH 6.0) and deprotonated (resembling pH 7.4) were measured from MD simulations (**Figure 2C**). Protonation of His285 led to an increase in RMSF across all four domains of AcrA; increased dynamics was most prominent in the αβ barrel and MP domains. The agreement of ΔRMSF, both extent and location, with the ΔHDX measurements supports that protonation of His285 alone dictates the structural mobility observed within more acidic conditions. Taken together, this suggests that, when considered in contexts where Mg^2+^ concentrations are limited, a more acidic periplasmic environment increases the global plasticity of AcrA.

Mg^2+^ has a key role in maintaining cell wall integrity of Gram-negative bacteria especially.^38^ Therefore, it is important to consider periplasmic protein structure-function relationships in the context of Mg^2+^, especially when considering its varying abundance within different media. Moreover, in the human small intestine, increased luminal acidity enhances Mg^2+^ absorption.^18,39^ We, therefore, investigated whether Mg^2+^ could influence the structural dynamics of AcrA within the two pH regimes (pH 7.4 and pH 6.0) previously tested.

ΔHDX experiments were first performed on AcrA^S^ at pH 6.0 comparing HDX profiles ± MgCl_2_ (**Figure 3**). Mg^2+^ had a significant stabilising effect on AcrA^S^, exhibited by decreased deuterium incorporation within areas across all four domains throughout the HDX time course. Most of the α-helical hairpin shows protection, whereas the other three domains have more limited portions of protection. Areas with insignificant ΔHDX suggest backbone dynamics within these regions are not affected by Mg^2+^. Interestingly, there is protection on several flexible linkers between AcrA’s domains; regions between the αβ barrel-MP domains (e.g. ^45^VKTEPLQITTELPGRTSAY^63^) and lipoyl-α-helical domains (e.g. ^90^GVSLYQIDPATYQATY^105^) display reduced HDX. These flexible linkers are known to be key for AcrA to freely position its four domains and change conformations.^32,40^ It is likely, therefore, that any backbone restriction of these areas influences the functionally required tertiary structure conformer populations, which AcrA is capable of forming during efflux. MD analysis of protonated AcrA ± Mg^2+^ also revealed that AcrA backbone dynamics are significantly stabilised in the presence of Mg^2+^, with reduced ΔRMSF across all four domains, which agrees with the ΔHDX (**Figure 3C**). Together this supports that within acidic conditions Mg^2+^ binding causes AcrA^S^ to become more rigid across its backbone (**Figure 3**). This structural rigidification is perhaps a contributing factor to the small, but statistically significant, 16% reduction in length of AcrA^S^ previously observed in the presence of MgCl_2_ by dynamic light scattering (DLS) and velocity centrifugation.^20^

**Figure 3.**
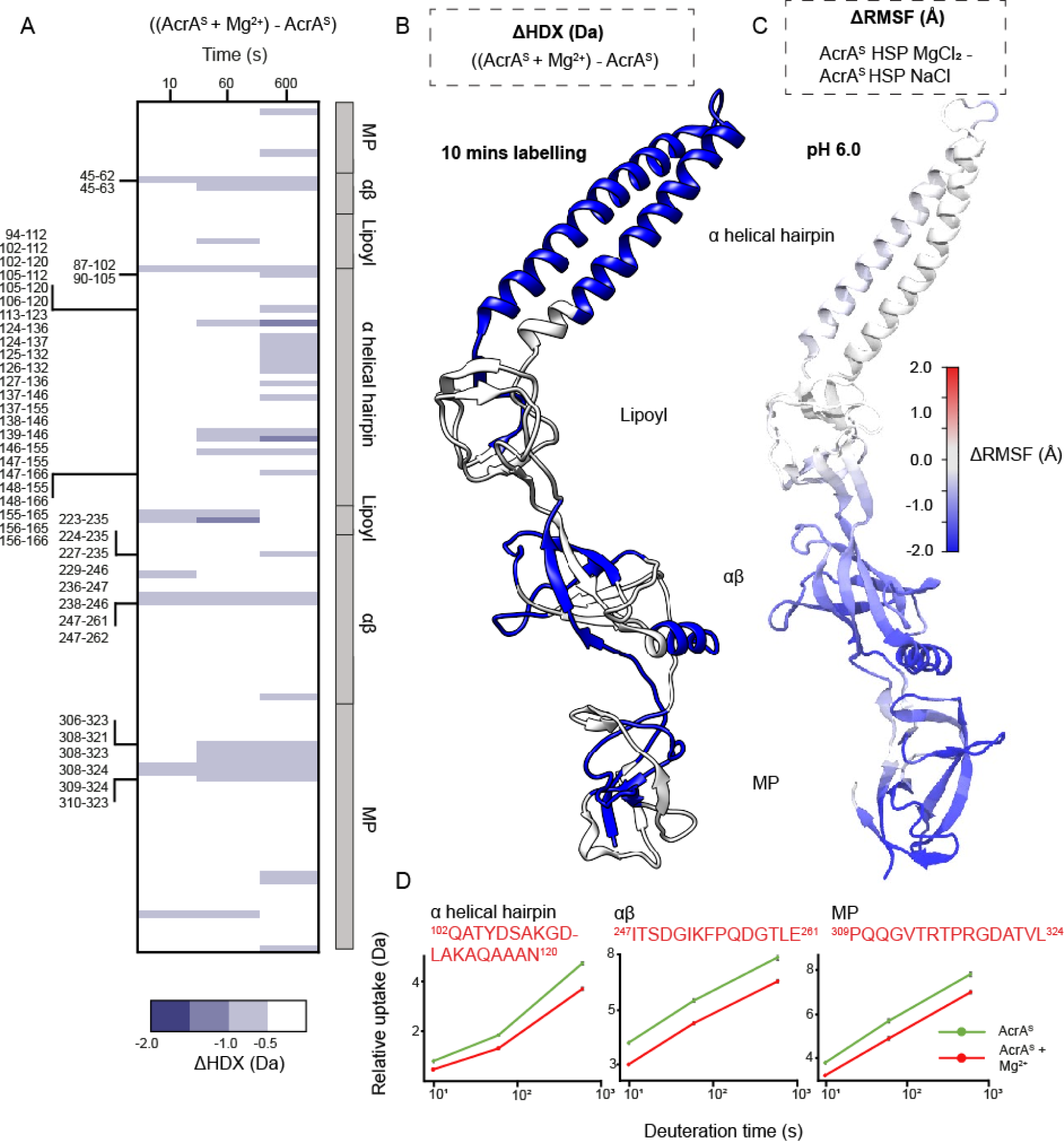
Influence of magnesium on the structural dynamics of AcrA at pH 6.0. **A.** Chiclet plot displaying the differential HDX (ΔHDX) plots for ((AcrA^S^ + Mg^2+^) - AcrA^S^), at pH 6.0. Blue signifies areas with decreased HDX between states and white signifies areas with no significant change in HDX. Significance was defined to be ≥ 0.5 Da change with a P-value ≤ 0.01 in a Welch’s t-test (n=4). **B.** ΔHDX for ((AcrA^S^ + Mg^2+^) - AcrA^S^) for the latest time point is coloured onto the AcrA structure (PDB:5O66) as performed in Figure 2B. **C.** AcrA coloured according to the difference in RMSF between simulations of the AcrA^S^ HSP (doubly protonated His285) state in the presence of MgCl_2_ and NaCl (AcrA^S^ HSP MgCl_2_ - AcrA^S^ HSP NaCl), averaged over four replicas for each. Plots presented as in Figure 2C**. D.** D-label uptake plots for three select peptides in different domains of AcrA. Uptake plots are the average deuterium uptake and error bars indicate the standard deviation (n=4).

Surprisingly, Mg^2+^ had no significant effect on the HDX profile of AcrA at pH 7.4, suggesting that it does not stabilise the backbone dynamics in the same manner as observed under acidic conditions (**Figure S6**). MD simulations supported that Mg^2+^ did not stabilise AcrA^s^ when His285 was deprotonated in the same manner as it did when His285 was protonated (**Figure 3**), with increased flexibility being observed in areas of the αβ barrel and MP domains instead (**Figure S7**). The structural effect of Mg^2+^, therefore, appears to be pH-dependent, only significantly acting on the backbone dynamics of AcrA within acidic conditions, when His285 is protonated, to generate a structural plasticity more like that found in neutral conditions. Interestingly, within our MD simulations a higher number of Mg^2+^ ions, when compared to Na^+^, reside near (within 7 Å) of both protonated and deprotonated His285; this was the case even when His285 was mutated to an Ala residue (**Table S2**). The localisation of Mg^2+^ to residues near His285 may be how Mg^2+^ enacts its influence over the local hydrogen bonding network managed by His285 (de)protonation.

### His285 protonation leads to specific domain alterations within a Mg^2+^ context

To conclude the relationship between Mg^2+^ binding at neutral and mildly acidic conditions, ΔHDX was performed between the two pH states in the context of magnesium (AcrA^S^ + Mg^2+^ at pH 6.0) - (AcrA^S^ + Mg^2+^ at pH 7.4) (**Figure 4**). Greater HDX profile parity was observed between pH 6.0 and 7.4 conditions, further supporting that Mg^2+^ compensates for increased dynamics exhibited by AcrA^S^ when His285 is protonated, with far fewer peptides possessing significantly increased HDX in the latest timepoint (**Figure 4**). However, from this ΔHDX analysis, more domain specific differences become pronounced, revealing pH-dependent stabilisation within the αβ barrel domain (e.g. peptide ^223^FLRLKQELANGTL^235^), which is protected across the entire HDX time course. Interestingly, stabilisation was prominent within the earliest HDX timepoint (10 seconds) (**Figure 4**), which points towards His285 protonation leading to a backbone structural rearrangement, rather than conformational plasticity changes typically observed at later timepoints like those observed for Mg^2+^ stabilisation at pH 6.0 (e.g. 10 minutes, **Figure 3**). The CD data supports this (**Figure S4F** and **Table S1**), showing that secondary structure is significantly altered by a transition from pH 7.4 to 6.0 only when in MgCl_2_; without Mg^2+^ the global structural content remains similar. Within identical Mg^2+^ conditions MD simulations show reduced dynamics within similar sections of the αβ and MP domains when His285 is protonated, as well as the increased dynamics of the α-helical hairpin tip.

**Figure 4.**
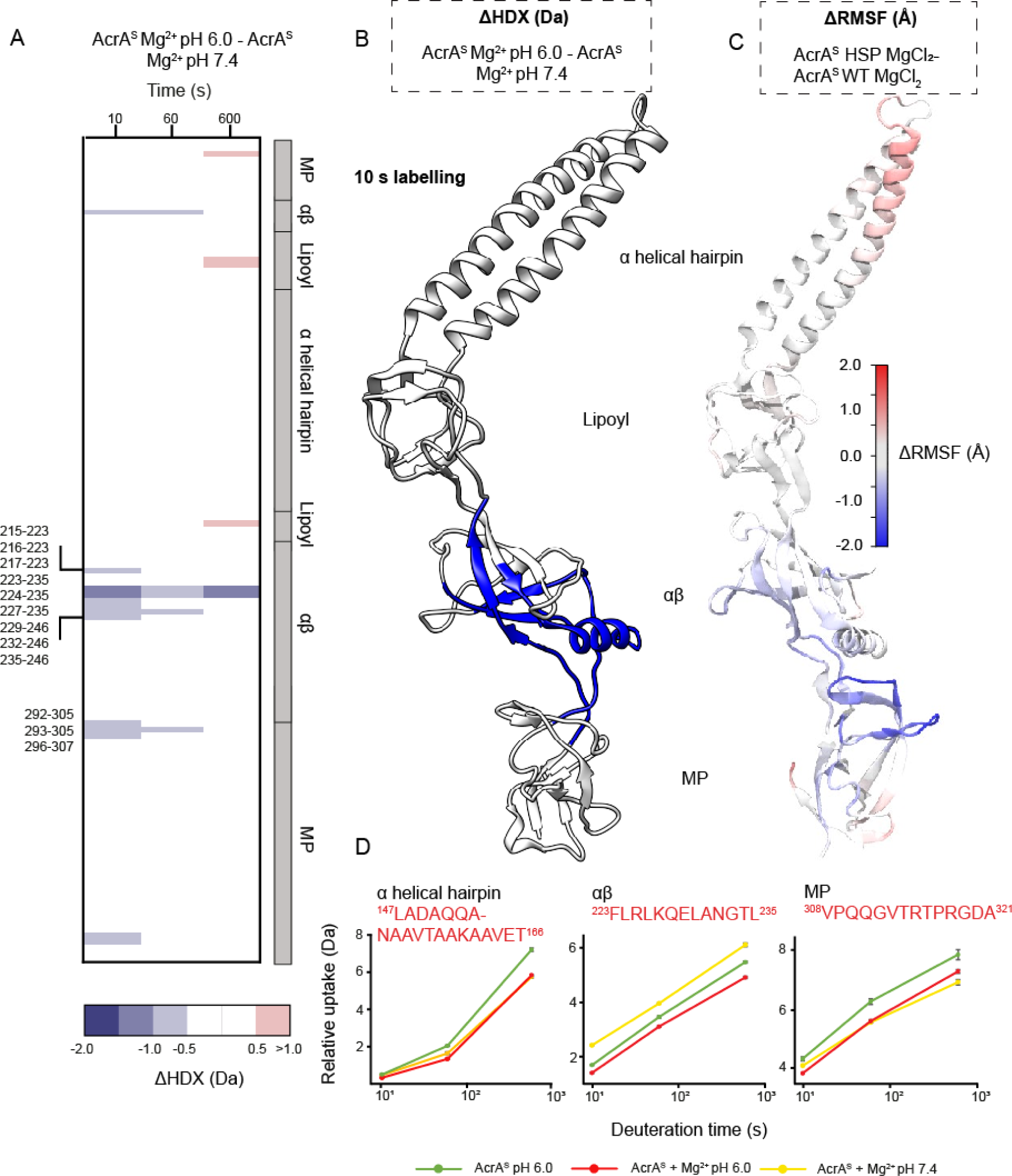
Effect of increased acidity on AcrA structural dynamics within a MgCl_2_ context. **A.** Chiclet plot displaying the differential HDX (ΔHDX) plots for ((AcrA^S^ + Mg^2+^ pH 6.0) - AcrA^S^ + Mg^2+^ pH 7.4). Blue signifies areas with decreased HDX between states, red signifies areas with increased HDX between states and white signifies areas with no significant change in HDX. Significance was defined to be ≥ 0.5 Da change with a P-value ≤ 0.01 in a Welch’s t-test (n=4). **B.** ΔHDX for ((AcrA^S^ + Mg^2+^) - AcrA^S^) for the earliest time point is coloured onto the AcrA structure (PDB:5O66) as performed in Figure 2B. **C.** AcrA coloured according to the difference in RMSF between simulations of the AcrA^S^ wild-type state and HSP (doubly protonated His285) state, in a MgCl_2_ environment (AcrA^S^ HSP MgCl_2_ - AcrA^S^ WT MgCl_2_), averaged over four replicas for each, presented as found in Figure 2C. **D.** Uptake plots for three peptides in different domains of AcrA. Uptake plots are the average deuterium uptake and error bars indicate the standard deviation (n=4).

Importantly, these results partly decouple the effects of Mg^2+^ from pH effects on AcrA structural dynamics, revealing that increasing acidity leads to more specific alterations within both the αβ barrel and MP domains. These pH-induced structural changes may produce the conformational organisation required for AcrA to function during efflux within more acidic contexts.

### His285 and Mg^2+^ are essential for pH-tolerant efflux pump activity

To analyse the effect of pH on AcrAB-TolC efflux activity *in vivo*, we constructed an AcrA H285A mutant (AcrA^H285A^) and compared the activities of this mutant and the wild-type AcrA in efflux bacterial growth-dependent and -independent assays. In the antibiotic susceptibility assay, the efflux deficient *E. coli* Δ9 cells complemented with AcrA^H285A^ variant were indistinguishable from cells complemented with the pump carrying the wild-type AcrA. Thus, at pH 7.4 the function of the pumps is not compromised by H285A substitution (**Table S3**). The immunoblotting analysis of the complemented cells showed that the expression levels of AcrA^H285A^ and the parent protein were similar (**Figure S8**).

Measurements of minimal inhibitory concentrations (MICs) at different pH are complicated by strong physiological responses as well as by physio-chemical properties of compounds and transmembrane diffusion. Hence, to analyse the effect of pH on efflux activity of AcrAB-TolC, we carried out the fluorescence probe efflux assay. In this assay, hyperporinated *E. coli* Δ9(Pore) cells carrying an empty vector or producing plasmid-borne AcrAB-TolC variants are incubated with increasing concentrations of probes, the fluorescence of which is enhanced when they bind to intracellular targets. As a result, the efflux activity is seen as lower fluorescence levels of the probes in efflux-proficient cells. As a probe we used *N*-phenyl-naphthylamide (NPN), a non-ionizable membrane probe (**Figure 5A**), which fluoresces when bound to the inner membrane of Gram-negative bacteria. At four different pH values, 8.0, 7.0, 6.0 and 5.0, NPN rapidly accumulated in efflux-deficient cells as seen from similar kinetics of changes of fluorescence intensity (**Figure S9**) and the calculated steady-state accumulation levels (**Figure 5B**). Surprisingly, the wild-type AcrAB-TolC maintained NPN efflux at all four pH values (**Figure S9** and **Figure 5C**). However, the efflux was the most efficient at pH 6.0-7.0 and declined by ∼50% at pH 5.0 and pH 8.0. Since the pKa of the imidazole ring of His is typically at 6.2, this result suggests that the optimal activity is when His285 exists in both the protonated and deprotonated states. In contrast, the activity of the pump assembled with AcrA^H285A^ was strongly pH-dependent (**Figure S9** and **Figure 5D**). Judging from the steady-state levels of NPN accumulation at the highest probe concentration of 32 μM, AcrA^H285A^ was twice as efficient as the wild type at pH 7.0 and somewhat less at pH 8.0. However, AcrA^H285A^ containing pump was ∼50% less efficient than the wild type at pH 6.0 and lost all its efflux activity at pH 5.0. Thus, the positively charged His285 in AcrA is required for the activity of AcrAB-TolC at acidic pH, whereas the alanine substitution in this position reducing the pH-dependence in this interface is favourite at pH 7.0-8.0.

**Figure 5.**
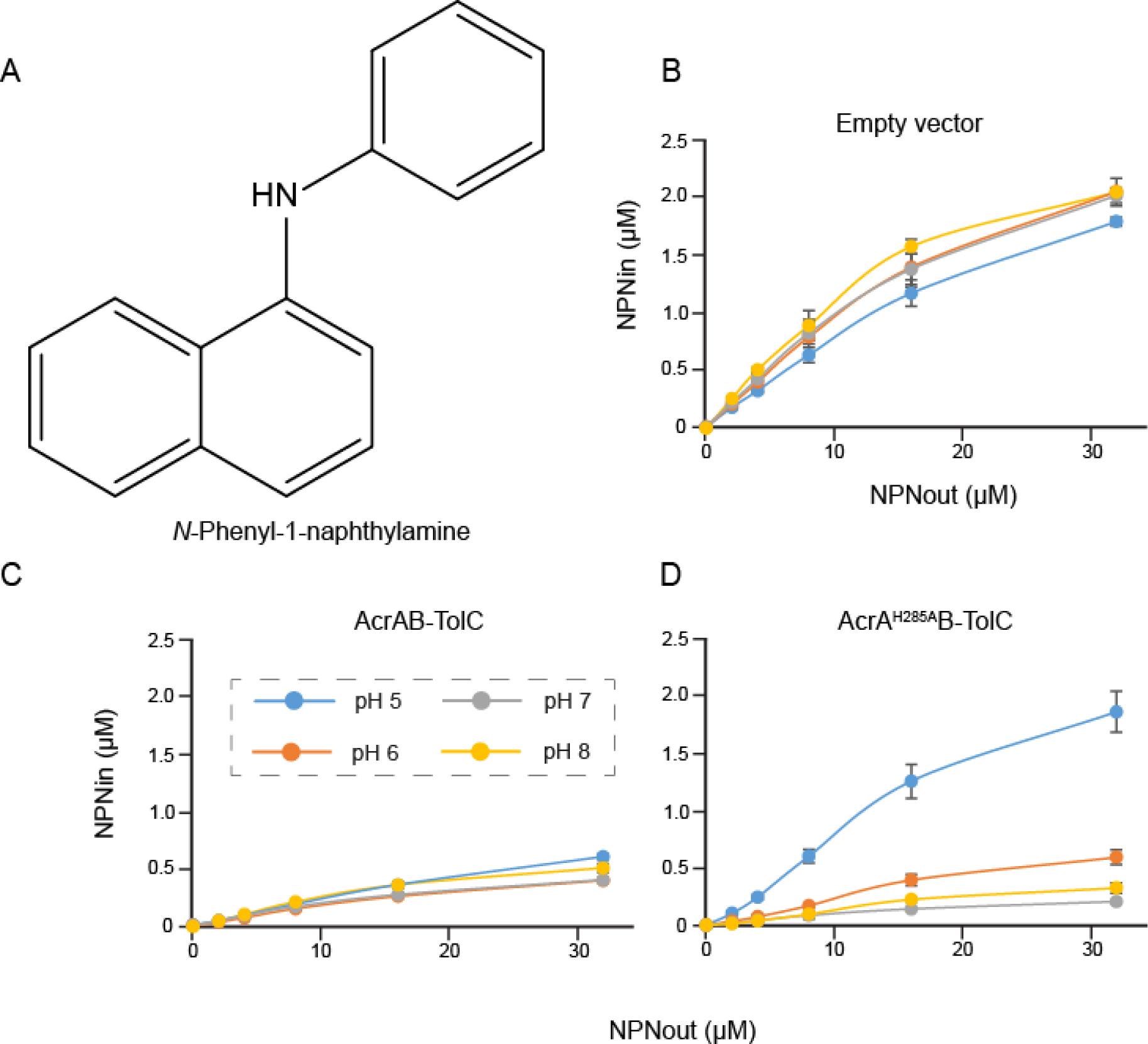
Activity of AcrAB-TolC is dependent on the protonation state of His285 of AcrA. **A.** Chemical structure of the non-ionisable efflux substrate NPN. **B.** Steady-state accumulation levels of the fluorescent probe NPN in E. coli Δ9(Pore) cells carrying an empty vector as a function of the external NPN concentration. Cells were induced with 0.1% arabinose to express an outer membrane Pore, washed, and resuspended in buffer with indicated pH values and supplemented with 5 mM MgCl_2_ and 0.1% glucose as an energy source. NPN was added in indicated concentrations (NPNout) and fluorescence was measured in real-time for 10 min. Kinetic curves (**Figure S8**) were fitted into a two-exponential kinetic model to calculate the steady-state levels of intracellular NPN (NPNin). **C.** The same as B, but cells produced the wild-type AcrAB-TolC **D**. The same as B but cells produced the AcrA^H285A^ containing efflux pump.

We also carried out the experiments in the presence and absence of MgCl_2_. However, in agreement with previous studies we found that Mg^2+^ is needed for the stabilization of the outer and inner membrane lipid bilayers and the cells were leaky and unstable in the buffer pH 6.0 without MgCl_2_ (**Figure S10**). As a result, no difference was seen between efflux-deficient and -proficient cells in the accumulation of NPN at pH 6.0. In the buffer pH 8.0 without MgCl_2_, the pump assembled with the wild-type AcrA was not able to overcome NPN influx, whereas the AcrA^H285A^ pump remained highly efficient (**Figure S10**). This result further supports that Mg^2+^ is required for wild-type AcrA function and the conclusion that alanine is preferred in the position 285 at pH 7.0 and above.

### NSC 60339 inhibitor conformationally restricts AcrA in the presence of Mg^2+^

Previously, we proposed a mechanism for the AcrA efflux pump inhibitor (EPI) NSC 60339.^12^ Using a combination of HDX-MS, MD simulations and cellular efflux assays, we suggested that NSC 60339 acts as a molecular wedge between the lipoyl and αβ barrel domains of AcrA, reducing its structural dynamics across all four domains. Our previous work utilised HDX-MS investigations at pH 6.0, reflecting the often more acidic conditions of the periplasmic environment, where AcrA resides.^12,16^ However, to investigate whether the inhibitor functions similarly with Mg^2+^ present, as it would be in the periplasm, HDX-MS experiments were repeated at pH 6.0, 1 mM MgCl_2_, and ± 500 μM NSC 60339 (containing 5% DMSO to maintain drug solubility) (**Figure S11**). First attempts at collecting this dataset saw that the Mg^2+^/DMSO induced extensive peptide carryover. Efforts to reduce carryover following previously described guidelines^41,42^ were made and we obtained high-quality data from a pulsed HDX experiment using a 251s labelling reaction at pH 6.0 (83 peptides, 92.9% protein coverage). This analysis showed that similar key regions across multiple domains of AcrA^S^ exhibited reduced structural dynamics by NSC 60339 in the presence of Mg^2+^, for example ^75^IILKRNFKEGSD^86^ in site IV and ^308^VPQQGVTRTPRGDATVL^32^ in the MP domain away from site IV showed the characteristic strong HDX protection (**Figure S11**). Furthermore, regions that previously saw no change in HDX were also consistent. Therefore, it appears that NSC 60339 still inhibits AcrA via the same mechanism when magnesium is present, with inhibitor binding between lipoyl and αβ barrel domains enacting conformational restriction of AcrA throughout specific areas across its domains (**Figure S10C**).

## Conclusions

*E. coli* and other bacteria experience frequent changes in pH of their external media, whether it is a human host or waste waters. Since the periplasm is accessible for most ions, the outer membrane and periplasmic proteins are expected to be able to withstand the rapid changes in pH of the medium. Indeed, we found that AcrAB-TolC is functional in the pH range 5.0-8.0 with the optimum at pH 6.0-7.0. The conformational state of AcrA is critical for the activity of the pump and our findings show that binding of Mg^2+^ within AcrA and (de)protonation of His285 likely work together to modulate the conformational states and dynamics of AcrA within the AcrAB-TolC efflux pump (**Figure 6**).

**Figure 6.**
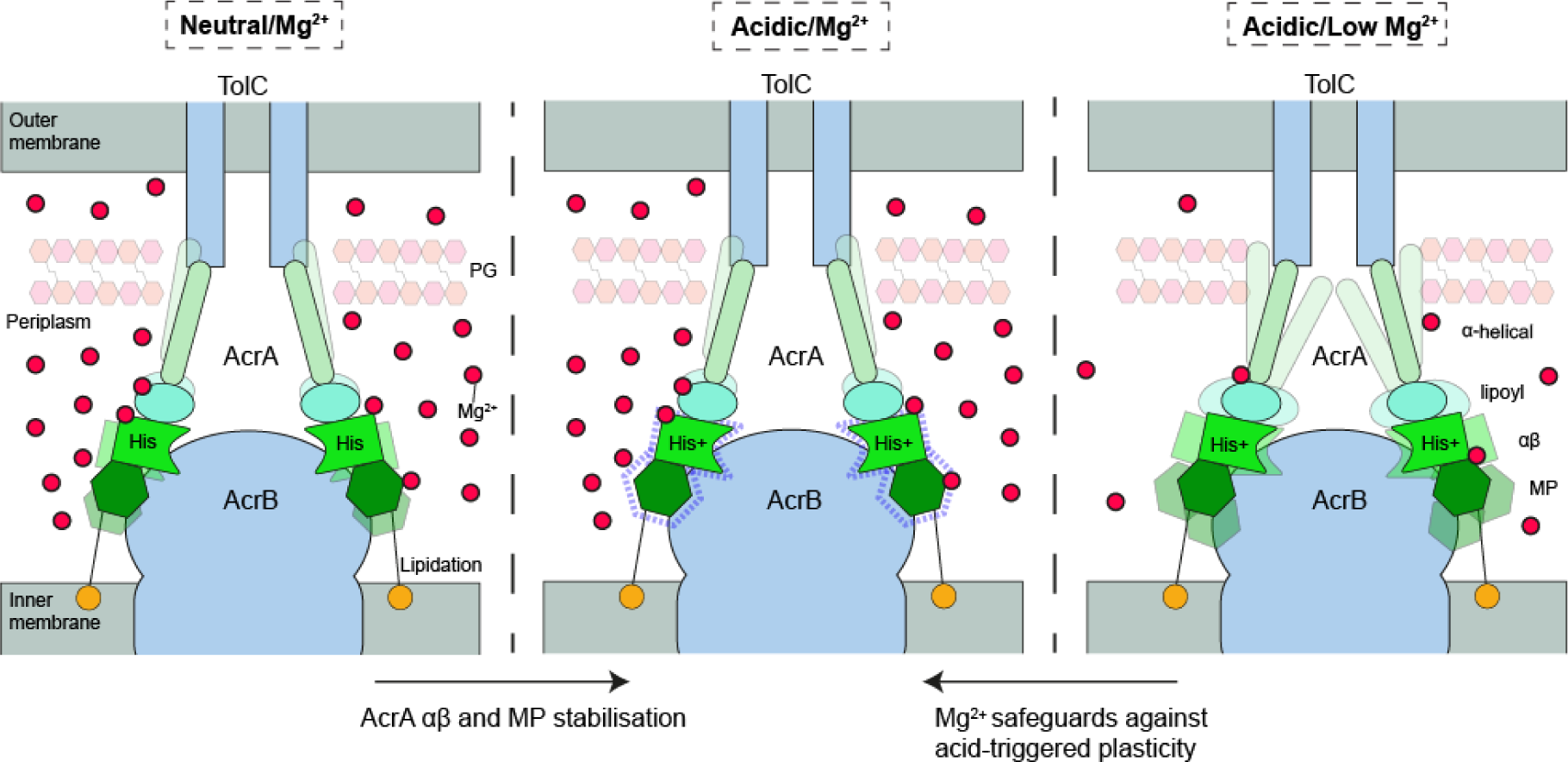
Proposed operational role of Mg^2+^ in modulating AcrA structural mobility to secure pump activity within acidic regimes. AcrA is a dynamic protein which ensures a sealed connecting pore between the AcrB transporter and the porin TolC. Importantly, AcrA transmits conformational motions during substrate recognition and translocation by AcrB for the gating of TolC to enable drug export from the periplasm to the outside of the Gram-negative bacterial wall. The periplasmic space has a pH like that of the external medium. When that external environment becomes more acidic, for example during enteric infection and colonization, then an acidified periplasmic space will occur and His285 becomes protonated. His285 being critical to modulating the hydrogen bonding network within AcrA. In a low Mg^2+^ environment AcrA undergoes a global increase in conformational plasticity which could prevent proper multidrug efflux pump function or pore concealment of its substrates. Whereas, within a Mg^2+^-rich environment, at acidic pH, regions of the αβ barrel domain are stabilised and exhibit structural reorganisation, likely to ensure proper function within increased acidity.

Interestingly, His285 of AcrA is in the NSC 60339 binding site and is surrounded by aspartate and arginine residues, the hydrogen bonding to which is expected to be sensitive to the protonation state of His285. The imidazole ring of NSC 60339 has a positive charge on it and can interfere with bonding of the positively charged His285 or its rearrangement at acidic pH. Previous MD and docking showed that NSC 60339 H-bonds with the backbone of His285, not its imidazole ring.^43^ Such binding to the backbone could prevent His285 from forming electrostatic interactions needed for the activity of AcrAB-TolC complex. The importance of these bonds is striking at pH 5.0, as the AcrA^H285A^ variant, but not the wild type, is inactivated. HDX and MD simulations together showed that at pH 6.0, when His285 is protonated, the backbone of AcrA is more dynamic, suggesting that the positive charge on His285 leads to rearrangements of contacts.

All these data point to the critical role of the αβ barrel domain of AcrA, which is located between the AcrA funnel formed by the lipoyl and the α-helical hairpin on one side and the MP domain on the other. The funnel and the MP domain play distinct roles in AcrAB-TolC function, with the funnel acting on TolC to open it and the MP domain acting on AcrB and stimulating its functional rotations. The αβ barrel domain interfaces with the lipoyl and the MP domains are stabilized by H-bonding, which is particularly sensitive to changes in pH. The lipoyl domain does not interact with AcrB, whereas the αβ and MP domains form two different interfaces with the AcrB trimer. In the *trans* state of AcrA seen in the cryo-EM structure of the resting AcrAB-TolC complex, the αβ barrel domain has only one contact with the MP domain.^32^ The two domains have multiple contacts when the free AcrA is transitioning into the *cis* state.^31^ The energy barrier between the two AcrA states is significant. In the *in situ* complex determined by cryo-ET, the interfaces of the αβ barrel domain of AcrA with the funnel and the MP domain have different states depending on the site of the interaction with AcrB.^10^ The MP domain is slightly rotated (8°) and also moves up and down perpendicular to the membrane plane, the motion requiring the rearrangements of the contacts of the αβ barrel domain with the MP domain and the funnel. Our findings suggest that inactivation of AcrAB-TolC due to rigidification of AcrA’s structure by NSC 60339 and the inactivation of AcrA^H285A^B-TolC at pH 5.0 due to the loss of critical contacts might have the same underlying molecular mechanism: the loss of critical contacts of the αβ barrel domain needed for transmission of conformational changes between AcrB and TolC.

This work suggests a wider role for Mg^2+^ in the function of periplasmic adaptor proteins, and thus multidrug efflux systems, to work across a variety of conditions the cell may exhibit. In this work we have shown that Mg^2+^ binding to AcrA broadly rectifies increased backbone dynamics exhibited under acidic conditions, whilst specifically stabilising the αβ barrel domain portion (**Figure 6**). As discussed above, the importance of this domain for pump function is well characterised; the localised stabilisation of the αβ barrel domain by Mg^2+^ specifically in mildly acidic conditions may offer a route to specialised conformations for robust efflux within these regimes. Mg^2+^ acting as a structural cofactor in this manner may be commonplace among other periplasmic adaptor proteins from Gram-negative bacteria and periplasmic lipoproteins more generally.

## Acknowledgements

The studies at King’s College London and the University of Southampton were supported by a UKRI Future Leaders Fellowship (MR/S015426/1, MR/X009580/1) to E.R., a Wellcome Trust Investigator Award (214259/Z/18/Z) and a BBSRC Pioneer Award (BB/Y512849/1) to P.J.B, and a King’s College London PhD studentship to B.R.L. The studies at University of Oklahoma and Georgia Institute of Technology were supported by the National Institutes of Health grant R01-AI052293 to H.I.Z. and J.C.G. Computational resources were provided through ACCESS (grant TG-MCB130173), which is supported by National Science Foundation grants 2138259, 2138286, 2138307, 2137603, and 2138296. This work also used the Hive cluster, which is supported by the NSF (MRI-1828187) and is managed by the Partnership for an Advanced Computing Environment at Georgia Tech.

## Contributions

B.R.L., M.R.U, K.M.K, J.C.G., H.I.Z., and E.R. designed the project; B.R.L., L.M.N.S., D.H., and E.R. performed mass spectrometry experiments and analysis; B.R.L. and N.J.H. performed biophysical experiments and analysis; B.R.L., M.R.U., E.R., and H.I.Z. cloned, purified, and characterised all protein constructs; M.R.U. and H.I.Z. performed bacterial efflux and susceptibility assays and analysis; K.M.K. and J.C.G. carried out molecular dynamics experiments and post-molecular dynamics analyses; P.J.B., D.H., J.C.G., H.I.Z., and E.R. supervised and/or financially supported the project; B.R.L., M.R.U., K.M.K, J.C.G., H.I.Z., and E.R. wrote the manuscript with input from the other authors.

## Competing interests

The authors declare no competing interests.

## Methods

### Cloning, expression, and purification

**AcrA^S^.** AcrA^S^ lacking the signal peptide residues 1-24, and the site of palmitoylation Cys25, was cloned in a pET28a plasmid with a 6xHis and LE linker.^20,46^ AcrA^S^ is purified from the cytoplasmic fraction as previously reported.^12,20^

### Circular dichroism

The circular dichroism (CD) experiments performed in this work were completed on a Chirascan V2 instrument. For standard CD scans, AcrA^S^ was buffer exchanged into protein buffer (50 mM NaHPO_4_, 150 mM NaCl, +/- 1 mM MgCl_2_ pH 6.0). Proteins were analysed at a concentration of 0.32 mg/mL. A coverslip was used with a pathlength of 0.005 cm. Scans were repeated three times between wavelengths 185-280 nm. BeStSeL online algorithm analysed the secondary structure.^30^

AcrA^S^ was buffer exchanged into the same buffer as before but diluted to 0.0075 mg/mL. Thermal melts were performed at 15 temperatures from 30-95 °C, with 5 °C increments to first identify where the transition from a folded state to an unfolded state occurs. Then a more accurate scan ranging between 40-60 °C with 1 °C increments was completed to find the melting point (*T_m_*). From this scan, values at 222 nm were taken for each temperature and thermodynamic parameters calculated from the following equations.

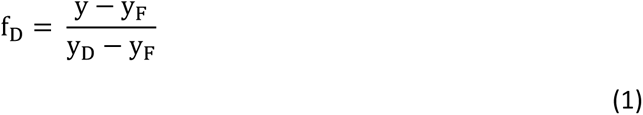

f_D_ is the fraction denatured, y_F_ is the gradient of folded protein and y_D_ is the gradient of the denatured protein. Spectra analysed on SigmaPlot.

### Hydrogen/deuterium exchange mass spectrometry

HDX-MS experiments were performed on a nanoAcquity ultra-performance liquid chromatography (UPLC) Xevo G2-XS QTof mass spectrometer system (Waters). Optimised peptide identification and peptide coverage for AcrA^S^ was performed from undeuterated controls. The optimal sample workflow for HDX-MS of AcrA^S^ was as follows: 5 μl of AcrA^S^ (20 μM) was diluted into 95 μl of either equilibration buffer (50 mM sodium phosphate, 150 mM NaCl, +/- 1mM MgCl_2_, pH 6.0/pH 7.4) or labelling buffer (deuterated equilibration buffer) at 20 °C. After fixed times of deuterium labelling, the samples were mixed with 100 μl of quench buffer (formic acid, 1.6 M GuHCl, 0.1% Fos-choline, pH 1.9) to provide a quenched sample at pH 2.4. 70 μl of quenched sample was then loaded onto a 50 μl sample loop before being injected onto an online Enzymate™ pepsin digestion column (Waters) in 0.1% formic acid in water (200 μl/min flow rate) at 20 °C. The peptic fragments were trapped onto an Acquity BEH c18 1.7 μM VANGUARD pre-column (Waters) for 3 min. The peptic fragments were then eluted using an 8- 35% gradient of 0.1% formic acid in acetonitrile (40 μl/min flow rate) into a chilled Acquity UPPLC BEH C18 1.7 μM 1.0 × 100mm column (Waters). The trap and UPLC were both maintained at 0 °C. The eluted peptides were ionized by electrospray into the Xevo G2-XS QTof mass spectrometer. MS^E^ data were acquired with a 20–30 V trap collision energy ramp for high-energy acquisition of product ions. Argon was used as the trap collision gas at a flow rate of 2 mL/min. Leucine enkephalin was used for lock mass accuracy correction and the mass spectrometer was calibrated with sodium iodide. The online Enzymate™ pepsin digestion column (Waters) was washed three times with pepsin wash (1.5 Gu-HCl, 4% MeOH, 0.8% formic acid, 0.1% Fos-choline).

All deuterium time points and controls were performed in triplicate/quadruplicate. Sequence identification was performed from MS^E^ data of digested undeuterated samples of AcrA^S^ using ProteinLynx Global Server 2.5.1 (PLGS) software (Waters). The output peptides were then filtered using DynamX (v. 3.0) using these parameters: minimum intensity of 1481, minimum and maximum peptide sequence length of 5 and 20 respectively, minimum MS/MS products of 1, minimum products per amino acid of 0.11, and a maximum MH+ error threshold of 5 ppm.^47^ All the spectra were visually examined and only those with a suitable signal to noise ratio were used for analysis. The amount of relative deuterium uptake for each peptide was determined using DynamX (v. 3.0) and are only corrected for back exchange when specified. The relative fractional uptake (RFU) was calculated from the following equation, where Y is the deuterium uptake for peptide a at incubation time (*t*), and D is the percentage of deuterium in the final labelling solution:

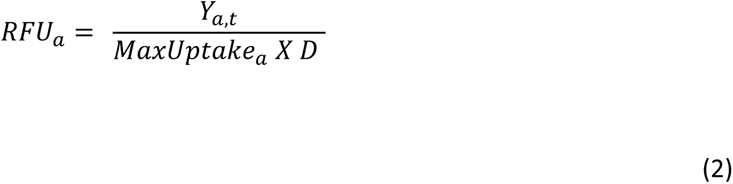

A significance level cut-off for experiments without biological replicates was set at 0.5 Da. This deuterium difference was greater than the confidence intervals calculated based on the pooled standard deviation values.^48^ Only peptides which satisfied the corresponding ΔHDX cut-off and a *P*-value ≤ 0.01 in a Welch’s t-test (n = 4) were considered significant. All ΔHDX structure figures were generated from the data using HDeXplosion and Chimera.^44,45^ The HDX-MS data generated in this study have been provided as Supplementary Data 1, and the HDX-MS summary tables and uptake plots have been provided as Supplementary Data 2 and 3, as per consensus guidelines.^41^

At pH 6.0, the rate of deuterium exchange is slower. Therefore, to compare proteins to pH 7.4, labelling times were adjusted using Equation 3.^36^

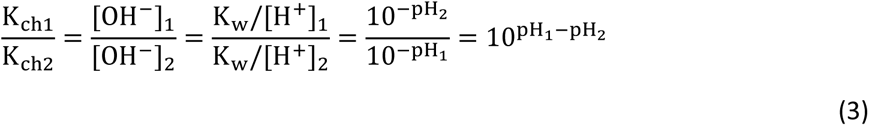

### Native mass spectrometry

AcrA^S^ was buffer exchanged into a volatile solution (100 mM ammonium acetate, pH 6.0) using a centrifugal exchange device (Micro Bio-Spin 6, Bio-Rad) according to manufacturer’s instructions native MS experiments were performed on a Synapt G2-Si mass spectrometer (Waters). The following instrument parameters, optimised to avoid ion activation and protein unfolding: capillary voltage: 1.6 kV, sampling cone: 30 V, trap DC bias; 15 V, trap collision energy: 2 V, transfer collision energy: 2 V. Pressures were set to 5.91 ×10^−2^ mbar in the source region (backing) and to 1.58 × 10^−2^ in both trap and collision cells. The collision gas was Helium.

### Metal-binding potential analysis

The *mebipred* (v 2.0) program^25^ was used to identify metal-binding potential of protein sequences. All protein sequences analysed were abstracted from UniProt^49^ and protein sequences which met the default cutoff of 0.5 within *mebipred* was considered an indication of metal-binding to either Fe, Ca, Na, K, Mg, Mn, Cu, K, Co, Ni metals (at this cutoff the first-tier model identified non-redundant metal-binding proteins with ∼92% precision at 26% recall). Global localisation annotation for *Escherichia coli* K-12 proteins was acquired from Sueki *et al.*,^26^ which utilize the database of the **s**ub-cellular **t**opology and localisation of the ***E****scherichia coli* **p**olypeptides (https://stepdb.eu/)^27,50^; except MdtE was defined as an inner membrane lipoprotein (IMLP).

### Site-directed mutagenesis and functional studies

The H285A amino acid substitution in AcrA was introduced by QuikChange II XL Site-Directed Mutagenesis Kit using p151AcrABHis as the template.^51^ Primer design and Polymerase chain reaction (PCR) reaction for each substitution were performed by following manufacturer’s protocol.

*E. coli* Δ9-Pore strain (Δ*acrB* Δ*acrD* Δ*acrEF::spc* Δ*emrB* Δ*emrY* Δ*entS::cam* Δ*macB* Δ*mdtC* Δ*mdtF att*Tn7::mini-Tn7T Tpr *araC* ParaBAD *fhuAΔC/Δ4L*)^52^ was used in antibiotic susceptibility and efflux assays. For these assays, Δ9-Pore cells were transformed with an empty pUC18 vector, p151AcrABHis or its p151AcrA^H285A^BHis derivative. Susceptibilities to novobiocin, SDS, erythromycin and vancomycin were determined by two-fold broth microdilution as described before.^12^

For efflux assays, overnight cultures of *E. coli* Δ9-Pore cells carrying pUC18, p151AcrABHis or p151AcrA^H285A^BHis plasmids were sub-cultured into a fresh LB medium supplemented with ampicillin (100 µg/ml final concentration) and incubated at 37°C with aeration until OD_600_ reached 0.1-0.3. The expression of the Pore was induced by addition 0.1% L-arabinose and incubation for another 4 hrs to reach OD_600_ ∼1.0.

Cells were washed with HMG buffer (50 mM HEPES-KOH buffer pH 7.0, 1 mM magnesium sulfate and 0.4 mM glucose) and resuspended in HMG buffer pH 5.0, 6.0, 7.0 or 8.0 to an OD_600_ of ∼1.0, at room temperature. Increasing concentrations of NPN were added to measure the substrate efflux efficiency of efflux-deficient Δ9-Pore(pUC18) and Δ9-Pore cells overproducing AcrAB or AcrA^H295A^B pumps. Kinetics of intracellular accumulation of NPN was monitored in at λex = 350 nm and λem = 405 nm in real time. Data was normalized and kinetic parameters were calculated as described previously using a MATLAB program.^53^ Briefly, the time courses of NPN uptake (**Figure S9**) were fit to the burst-single exponential decay function *F* = *A_1_*+*A_2_*·(1-exp(-*kt*)), where *A_1_* and *A_2_* describe the magnitude of the fast and slow steps, respectively, and *k* is the rate of the slow step. The fast and slow steps were attributed to NPN binding to the cell surface and in the inner membrane, respectively.

### Molecular dynamics simulations

To investigate the effects of pH and ion content on the AcrA monomer, twelve distinct systems were built and simulated using MD (**Table S4**). The AcrA monomer was initiated in either the cis or trans conformation.^31^ The systems were built in TIP3P water boxes of uniform dimensions of 171 Å per side and ionized with either 0.15 M NaCl or 0.15 M MgCl_2_.^54^ The systems were ∼470,000 atoms each. His285 was singly protonated to reflect the pH 7.0 experimental conditions. To reflect pH 6.0 conditions, His285 was doubly protonated. An additional set of systems were built for the H285A mutation. Two replicas of each system were run for a total of 24 simulations and a net simulation time of 12 μs.

Each system was equilibrated with an initial minimization followed by (1) relaxation of the water and ion molecules for 1 ns and (2) relaxation of the protein sidechains for 1 ns. An additional minimization step was applied (5000 steps) before beginning the production runs. The equilibration steps were performed with NAMD 2.14 with the productions performed with NAMD3.^55^ Hydrogen mass repartitioning was applied to enable a timestep of 4 fs.^56^ All simulations were done at constant temperature (310K) with Langevin dynamics and a damping coefficient of 1/ps, at constant pressure (1 atm) with the Langevin piston barostat, and with periodic boundary conditions. Short range non-bonded interactions were cut off at 12 Å with the force-based switching function starting at 10 Å. Long range interactions were calculated with particle-mesh Ewald method with a grid density of 1/Å^3^.^57^ The CHARMM36m force-field was used for the protein.^58^

All analysis and visualization were done with VMD.^59^ In addition, analysis shown represents the averages of four replicas – across both *cis* and *trans* conformations and two replicas for each condition. RMSF was calculated per residue over the last 470 ns of the simulation, discarding the first 30 ns as done previously.^12^ The number of ions within 7 Å of each residue in the AcrA monomer was also calculated across the entire simulation.

## Notes

### Competing Interest Statement

The authors have declared no competing interest.

## References

1. Global Leaders Group on AMR (2024). Towards specific commitments and action in the response to antimicrobial resistance https://www.amrleaders.org/resources/m/item/glg-report.

2. Nikaido, H. (2009). Multidrug Resistance in Bacteria. Annu Rev Biochem. 78, 119–146. 10.1590/S1807-59322011001000006.

3. Alav, I., Bavro, V.N., and Blair, J.M.A. (2021). Interchangeability of periplasmic adaptor proteins AcrA and AcrE in forming functional efflux pumps with AcrD in *Salmonella enterica* serovar Typhimurium. J. Antimicrob. Chemother. 76, 2558–2564. 10.1093/jac/dkab237.

4. Tseng, T.T., Gratwick, K.S., Kollman, J., Park, D., Nies, D.H., Goffeau, A., and Saier, M.H. (1999). The RND permease superfamily: An ancient, ubiquitous and diverse family that includes human disease and development proteins. J. Mol. Microbiol. Biotechnol. 1, 107–125.

5. Rice, L.B. (2008). Federal funding for the study of antimicrobial resistance in nosocomial pathogens: No ESKAPE. J. Infect. Dis. 197, 1079–1081. 10.1086/533452.

6. Piddock, L.J.V. (2006). Multidrug-resistance efflux pumps — not just for resistance. Nat. Rev. Microbiol. 4, 629–636. 10.1038/nrmicro1464.

7. Du, D., Wang-Kan, X., Neuberger, A., van Veen, H.W., Pos, K.M., Piddock, L.J.V., and Luisi, B.F. (2018). Multidrug efflux pumps: structure, function and regulation. Nat. Rev. Microbiol. 16, 523–539. 10.1038/s41579-018-0048-6.

8. Fernando, D.M., and Kumar, A. (2013). Resistance-Nodulation-Division multidrug efflux pumps in Gram-negative bacteria: Role in virulence. Antibiotics. 2, 163–181. 10.3390/antibiotics2010163.

9. Zgurskaya, H.I., Weeks, J.W., Ntreh, A.T., Nickels, L.M., and Wolloscheck, D. (2015). Mechanism of coupling drug transport reactions located in two different membranes. Front. Microbiol. 6, 1–13. 10.3389/fmicb.2015.00100.

10. Chen, M., Shi, X., Yu, Z., Fan, G., Serysheva, I.I., Baker, M.L., Luisi, B.F., Ludtke, S.J., and Wang, Z. (2022). In situ structure of the AcrAB-TolC efflux pump at subnanometer resolution. Structure. 30, 107–113. 10.1016/j.str.2021.08.008.

11. Abdali, N., Parks, J.M., Haynes, K.M., Chaney, J.L., Green, A.T., Wolloscheck, D., Walker, J.K., Rybenkov, V. V., Baudry, J., Smith, J.C., et al. (2017). Reviving antibiotics: Efflux pump inhibitors that interact with AcrA, a membrane fusion protein of the AcrAB-TolC multidrug efflux pump. ACS Infect. Dis. 3, 89–98. 10.1021/acsinfecdis.6b00167.

12. Lewis, B.R., Uddin, M.R., Moniruzzaman, M., Kuo, K.M., Higgins, A.J., Shah, L.M.N., Sobott, F., Parks, J.M., Hammerschmid, D., Gumbart, J.C., et al. (2023). Conformational restriction shapes the inhibition of a multidrug efflux adaptor protein. Nat. Commun. 14, 1–14. 10.1038/s41467-023-39615-x.

13. De Oliveira, D.M.P., Forde, B.M., Kidd, T.J., Harris, P.N.A., Schembri, M.A., Beatson, S.A., Paterson, D.L., and Walker, M.J. (2020). Antimicrobial resistance in ESKAPE pathogens. Clin. Microbiol. Rev. 33, 1–19. 10.1128/CMR.00181-19.

14. Miller, S.I., and Salama, N.R. (2018). The gram-negative bacterial periplasm: Size matters. PLoS Biol. 16, 1–7. 10.1371/journal.pbio.2004935.

15. Nagy, T.A., Crooks, A.L., Quintana, J.L.J., and Detweiler, C.S. (2020). Clofazimine Reduces the Survival of *Salmonella enterica* in Macrophages and Mice. ACS Infect. Dis. 6, 1238–1249. 10.1021/acsinfecdis.0c00023.

16. Wang, B., Weng, J., Fan, K., and Wang, W. (2012). Interdomain flexibility and pH-induced conformational changes of AcrA revealed by molecular dynamics simulations. J. Phys. Chem. B. 116, 3411–3420. 10.1021/jp212221v.

17. Wilks, J.C., and Slonczewski, J.L. (2007). pH of the cytoplasm and periplasm of *Escherichia coli*: Rapid measurement by green fluorescent protein fluorimetry. J. Bacteriol. 189, 5601–5607. 10.1128/JB.00615-07.

18. Chamniansawat, S., Suksridechacin, N., and Thongon, N. (2023). Current opinion on the regulation of small intestinal magnesium absorption. World J. Gastroenterol. 29, 332–342. 10.3748/wjg.v29.i2.332.

19. Alegun, O., Pandeya, A., Cui, J., Ojo, I., and Wei, Y. (2021). Donnan Potential across the Outer Membrane of Gram-Negative Bacteria and Its Effect on the Permeability of Antibiotics. Antibiotics. 10, 1–16. 10.3390/antibiotics10060701.

20. Zgurskaya, H.I., and Nikaido, H. (1999). AcrA is a Highly Asymmetric Protein Capable of Spanning the Periplasm. J. Mol. Biol. 285, 409–420. 10.1006/jmbi.1998.2313.

21. Ip, H., Stratton, K., Zgurskaya, H., and Liu, J. (2003). pH-induced Conformational Changes of AcrA, the Membrane Fusion Protein of *Escherichia coli* Multidrug Efflux System. J. Biol. Chem. 278, 50474–50482. 10.1074/jbc.M305152200.

22. De Angelis, F., Lee, J.K., O’Connell, J.D., Miercke, L.J.W., Verschueren, K.H., Srinivasan, V., Bauvois, C., Govaerts, C., Robbins, R.A., Ruysschaert, J.M., et al. (2010). Metal-induced conformational changes in ZneB suggest an active role of membrane fusion proteins in efflux resistance systems. Proc. Natl. Acad. Sci. U.S.A. 107, 11038–11043. 10.1073/pnas.1003908107.

23. Bagai, I., Liu, W., Rensing, C., Blackburn, N.J., and McEvoy, M.M. (2007). Substrate-linked conformational change in the periplasmic component of a Cu(I)/Ag(I) efflux system. J. Biol. Chem. 282, 35695–35702. 10.1074/jbc.M703937200.

24. Tang, S., and Yang, J.J. (2013). Encyclopedia of Metalloproteins. In Encyclopedia of Metalloproteins (Springer Science), pp. 1243–1250.

25. Aptekmann, A.A., Buongiorno, J., Giovannelli, D., Glamoclija, M., Ferreiro, D.U., and Bromberg, Y. (2022). mebipred: identifying metal-binding potential in protein sequence. Bioinformatics. 38, 3532–3540. 10.1093/bioinformatics/btac358.

26. Sueki, A., Stein, F., Savitski, M.M., Selkrig, J., and Typas, A. (2020). Systematic Localization of *Escherichia coli* Membrane Proteins. mSystems 5, 808–819. 10.1128/msystems.00808-19.

27. Orfanoudaki, G., and Economou, A. (2014). Proteome-wide subcellular topologies of *E. coli* polypeptides database (STEPdb). Mol. Cell. Proteomics. 13, 3674–3687. 10.1074/mcp.O114.041137.

28. Abramson, J., Adler, J., Dunger, J., Evans, R., Green, T., Pritzel, A., Ronneberger, O., Willmore, L., Ballard, A.J., Bambrick, J., et al. (2024). Accurate structure prediction of biomolecular interactions with AlphaFold 3. Nature. 10.1038/s41586-024-07487-w.

29. Greenfield, N.J. (2007). Using circular dichroism spectra to estimate protein secondary structure. Nat. Protoc. 1, 2876–2890. 10.1038/nprot.2006.202.

30. Micsonai, A., Wien, F., Bulyáki, É., Kun, J., Moussong, É., Lee, Y.H., Goto, Y., Réfrégiers, M., and Kardos, J. (2018). BeStSel: A web server for accurate protein secondary structure prediction and fold recognition from the circular dichroism spectra. Nucleic Acids Res. 46, 315–322. 10.1093/nar/gky497.

31. Hazel, A.J., Abdali, N., Leus, I. V., Parks, J.M., Smith, J.C., Zgurskaya, H.I., and Gumbart, J.C. (2019). Conformational Dynamics of AcrA Govern Multidrug Efflux Pump Assembly. ACS Infect. Dis. 5, 1926–1935. 10.1021/acsinfecdis.9b00273.

32. Du, D., Wang, Z., James, N.R., Voss, J.E., Klimont, E., Ohene-Agyei, T., Venter, H., Chiu, W., and Luisi, B.F. (2014). Structure of the AcrAB-TolC multidrug efflux pump. Nature. 509, 512–515. 10.1038/nature13205.

33. Kim, J.S., Jeong, H., Song, S., Kim, H.Y., Lee, K., Hyun, J., and Ha, N.C. (2015). Structure of the tripartite multidrug efflux pump AcrAB-TolC suggests an alternative assembly mode. Mol. Cells. 38, 180–186. 10.14348/molcells.2015.2277.

34. Thanassi, D.G., Suh, G.S.B., and Nikaido, H. (1995). Role of outer membrane barrier in efflux-mediated tetracycline resistance of *Escherichia coli*. J. Bacteriol. 177, 998–1007. 10.1128/jb.177.4.998-1007.1995.

35. Chalmers, M.J., Busby, S.A., Pascal, B.D., West, G.M., and Griffin, P.R. (2011). Differential hydrogen/deuterium exchange mass spectrometry analysis of protein-ligand interactions. Expert Rev. Proteomics. 8, 43–59. 10.1586/epr.10.109.

36. Li, J., Rodnin, M. V, Ladokhin, A.S., and Gross, M.L. (2014). Hydrogen−Deuterium Exchange and Mass Spectrometry Reveal the pH-Dependent Conformational Changes of Diphtheria Toxin T Domain. Biochemistry. 53, 6849–6854. 10.1021/bi500893y.

37. Fowler, M.L., McPhail, J.A., Jenkins, M.L., Masson, G.R., Rutaganira, F.U., Shokat, K.M., Williams, R.L., and Burke, J.E. (2016). Using hydrogen deuterium exchange mass spectrometry to engineer optimized constructs for crystallization of protein complexes: Case study of PI4KIIIβ with Rab11. Protein Sci. 25, 826–839. 10.1002/pro.2879.

38. Silhavy, T.J., Kahne, D., and Walker, S. (2010). The Bacterial Cell Envelope. Cold Spring Harb Perspect Biol. 2, 1–16. 10.1101/cshperspect.a000414.

39. Heijnen, A.M.P., Brink, E.J., Lemmens, A.G., and Beynen, A.C. (1993). Ileal pH and apparent absorption of magnesium in rats fed on diets containing either lactose or lactulose. Br. J. Nutr. 70, 747–756. 10.1079/bjn19930170.

40. Mikolosko, J., Bobyk, K., Zgurskaya, H.I., and Ghosh, P. (2006). Conformational flexibility in the multidrug efflux system protein AcrA. Structure. 14, 577–587. 10.1016/j.str.2005.11.015.

41. Masson, G.R., Burke, J.E., Ahn, N.G., Anand, G.S., Borchers, C., Brier, S., Bou-assaf, G.M., Engen, J.R., Englander, S.W., Faber, J., et al. (2019). Recommendations for performing, interpreting and reporting hydrogen deuterium exchange mass spectrometry (HDX-MS) experiments. Nat. Methods. 16, 595–602. 10.1038/s41592-019-0459-y.

42. Ball, D., Nguyen, T., Zhang, N., and Arcy, S.D. (2022). Using hydrogen-deuterium exchange mass spectrometry to characterize Mtr4 interactions with RNA. Methods Enzymol. 673, 475–516. 10.1016/bs.mie.2022.04.002.Using.

43. Darzynkiewicz, Z.M., Green, A.T., Abdali, N., Hazel, A., Fulton, R.L., Kimball, J., Gryczynski, Z., Gumbart, J.C., Parks, J.M., Smith, J.C., et al. (2019). Identification of Binding Sites for Efflux Pump Inhibitors of the AcrAB-TolC Component AcrA. Biophys. J. 116, 648–658. 10.1016/j.bpj.2019.01.010.

44. Zhang, N., Yu, X., Zhang, X., and Arcy, S.D. (2021). Structural bioinformatics HD-eXplosion: visualization of hydrogen – deuterium exchange data as chiclet and volcano plots with statistical filtering. Bioinformatics. 37, 1926–1927. 10.1093/bioinformatics/btaa892.

45. Pettersen, E.F., Goddard, T.D., Huang, C.C., Couch, G.S., Greenblatt, D.M., Meng, E.C., and Ferrin, T.E. (2004). UCSF Chimera — A Visualization System for Exploratory Research and Analysis. J. Comput. Chem. 25, 1605–1612. 10.1002/jcc.20084.

46. Tikhonova, E.B., Yamada, Y., and Zgurskaya, H.I. (2011). Sequential Mechanism of Assembly of Multidrug Efflux Pump AcrAB-TolC. Chem. Biol. 18, 454–463. 10.1016/j.chembiol.2011.02.011.

47. Sørensen, L., and Salbo, R. (2018). Optimized Workflow for Selecting Peptides for HDX-MS Data Analyses. J. Am. Soc. Mass Spectrom. 29, 2278–2281. 10.1007/s13361-018-2056-1.

48. Hageman, T.S., and Weis, D.D. (2019). Reliable Identification of Significant Differences in Differential Hydrogen Exchange-Mass Spectrometry Measurements Using a Hybrid Significance Testing Approach. Anal. Chem. 91, 8008–8016. 10.1021/acs.analchem.9b01325.

49. The UniProt Consortium (2023). UniProt: the Universal Protein Knowledgebase in 2023. Nucleic Acids Res. 51, 523–531. 10.1093/nar/gkac1052.

50. Loos, M.S., Ramakrishnan, R., Vranken, W., Tsirigotaki, A., Tsare, E.P., Zorzini, V., De Geyter, J., Yuan, B., Tsamardinos, I., Klappa, M., et al. (2019). Structural basis of the subcellular topology landscape of *Escherichia coli*. Front. Microbiol. 10, 1–22. 10.3389/fmicb.2019.01670.

51. Tikhonova, E.B., and Zgurskaya, H.I. (2004). AcrA, AcrB, and TolC of *Escherichia coli* form a stable intermembrane multidrug efflux complex. J. Biol. Chem. 279, 32116–32124. 10.1074/jbc.M402230200.

52. Krishnamoorthy, G., Wolloscheck, D., Weeks, J.W., Croft, C., Rybenkov, V. V., and Zgurskaya, H.I. (2016). Breaking the permeability barrier of *Escherichia coli* by controlled Hyperporination of the outer membrane. Antimicrob. Agents Chemother. 60, 7372–7381. 10.1128/AAC.01882-16.

53. Westfall, D.A., Krishnamoorthy, G., Wolloscheck, D., Sarkar, R., Zgurskaya, I., and Rybenkov, V. V (2017). Bifurcation kinetics of drug uptake by Gram-negative bacteria. PLoS One. 12, 1–18. 10.1371/journal.pone.0184671.

54. Jorgensen, W.L., Chandrasekhar, J., Madura, J.D., Impey, R.W., and Klein, M.L. (1983). Comparison of simple potential functions for simulating liquid water. J. Chem. Phys. 79, 926–935. 10.1063/1.445869.

55. Phillips, J.C., Hardy, D.J., Maia, J.D.C., Stone, J.E., Ribeiro, J. V., Bernardi, R.C., Buch, R., Fiorin, G., Hénin, J., Jiang, W., et al. (2020). Scalable molecular dynamics on CPU and GPU architectures with NAMD. J. Chem. Phys. 153, 1–33. 10.1063/5.0014475.

56. Balusek, C., Hwang, H., Lau, C.H., Lundquist, K., Hazel, A., Pavlova, A., Lynch, D.L., Reggio, P.H., Wang, Y., and Gumbart, J.C. (2019). Accelerating Membrane Simulations with Hydrogen Mass Repartitioning. J. Chem. Theory Comput. 15, 4673–4686. 10.1021/acs.jctc.9b00160.

57. Darden, T., York, D., and Pedersen, L. (1993). Particle mesh Ewald: An N·log(N) method for Ewald sums in large systems. J. Chem. Phys. 98, 10089–10092. 10.1063/1.464397.

58. Huang, J., Rauscher, S., Nawrocki, G., Ran, T., Feig, M., de Groot, B.L., Grubmüller, H., and and MacKerell Jr, A.D. (2017). CHARMM36m: An Improved Force Field for Folded and Intrinsically Disordered Proteins. Nat. Methods. 176, 139–148. 10.1038/nmeth.4067.CHARMM36m.

59. Humphrey, W., Dalke, A., and Schulten, K. (1996). VMD: Visual Molecular Dynamics. J. Mol. Graph. 14, 33–38. 10.1016/0263-7855(96)00018-5.

