## Supporting Information for "Mg^2+^-dependent mechanism of environmental versatility in a multidrug efflux pump"

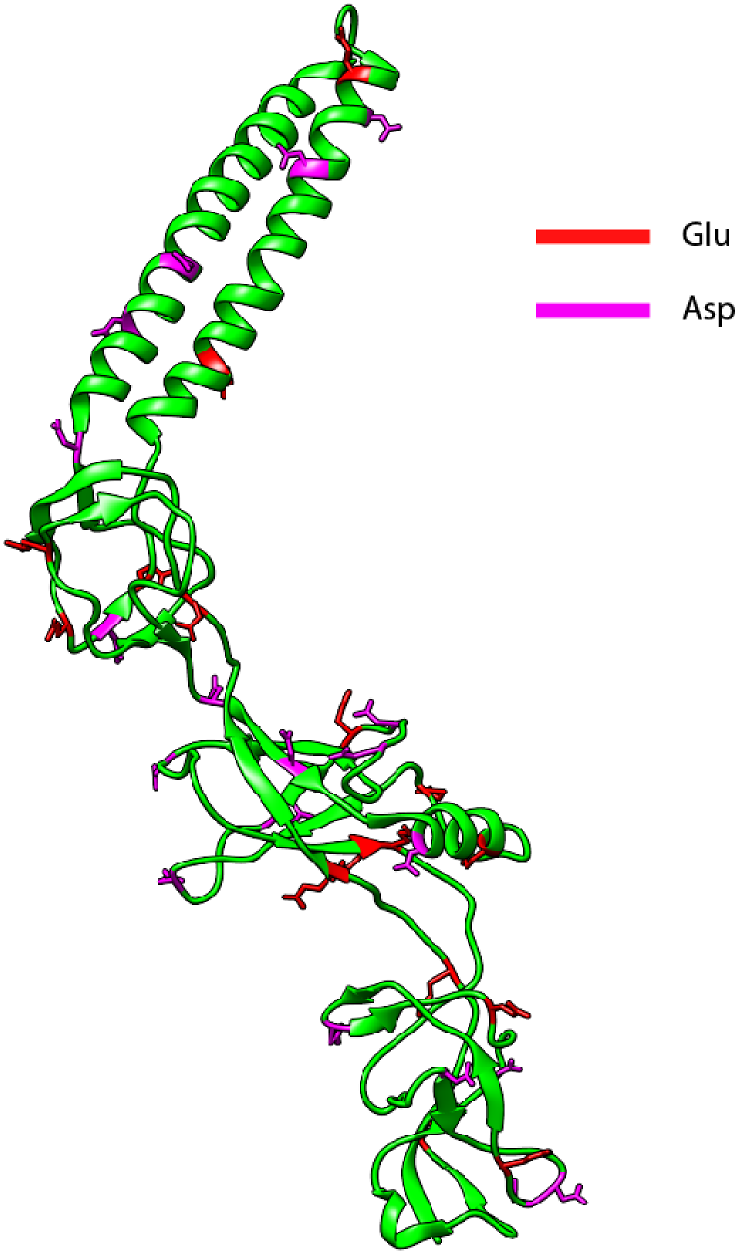

**Figure S1.** ***Location of Asp and Glu residues on AcrA.*** *Mapping of the Asp and Glu residues on AcrA. Asp residues labelled pink, Glu residues labelled red.*

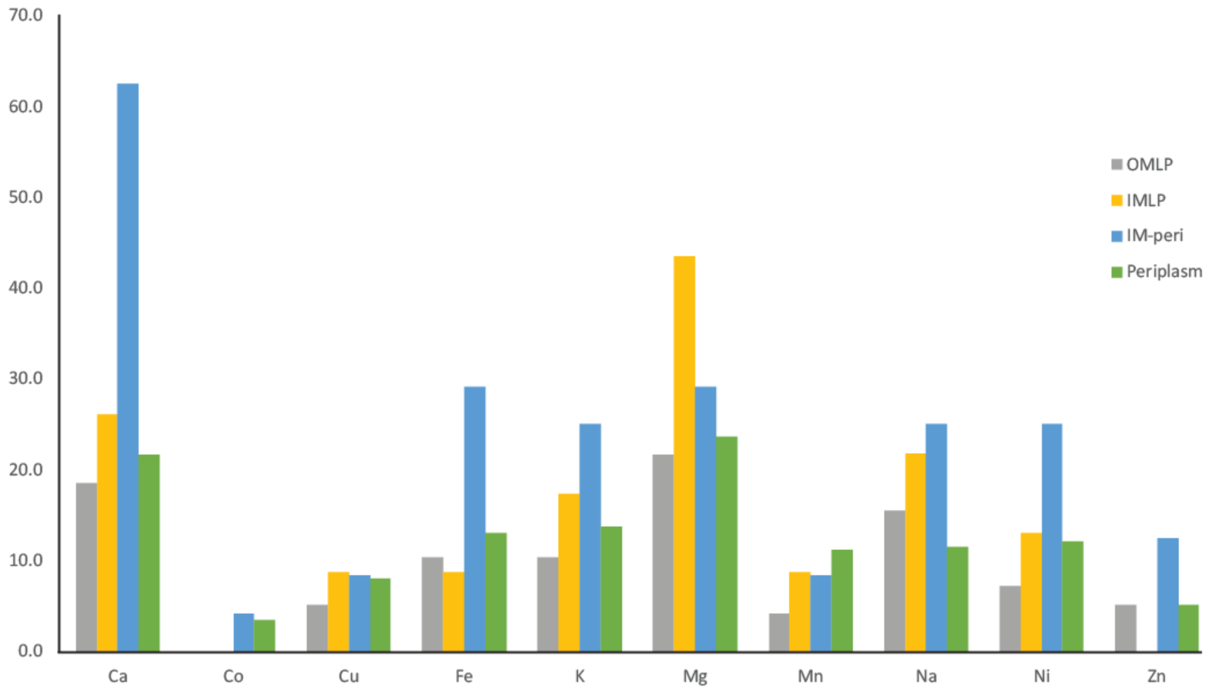

***Figure S2. E. coli proteome within the periplasmic space****. MeBiPred analysis on the entire subcellular E. coli proteome. Plotted are the theroretical total perecentage of proteins predicted to bind different metals. OMLP; outer membrane lipoprotein, IMLP; inner membrane lipoprotein, IM-peri; peripheral inner membrane protein, periplasm; periplasmic protein.*

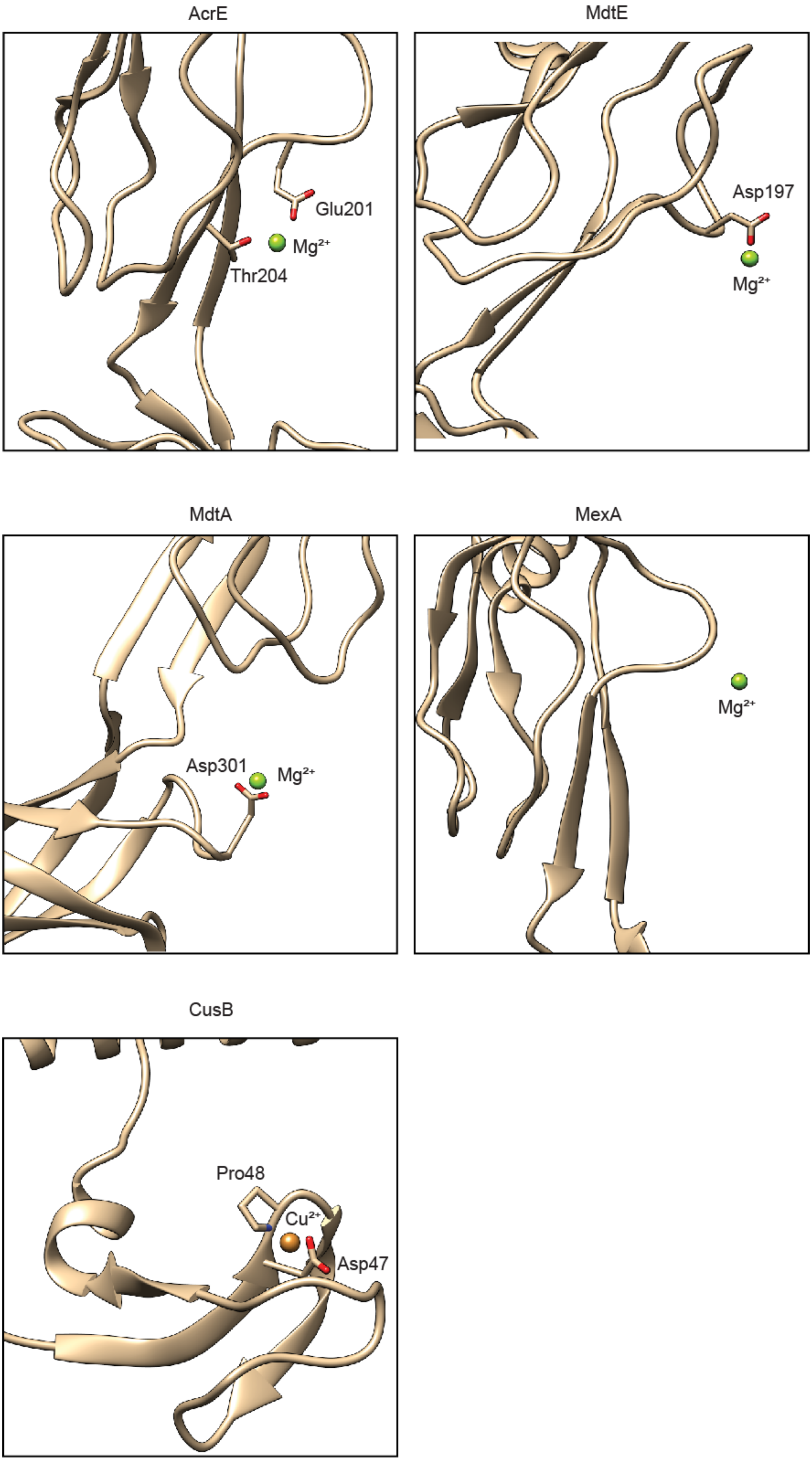
AcrE
Glu201
Mg²⁺
Thr204
MdtE
Asp197
Mg²⁺
MdtA
Asp301
Mg²⁺
MexA
Mg²⁺
CusB
Pro48
Cu²⁺
Asp47

***Figure S3. AlphaFold3 predictions of PAPs and cations.*** *AlphaFold3 predictions of MFPs and cation binding. For AcrE and* $Mg^{2+}$ *the reported predicted template modelling (pTM) score and the interface predicted template modelling (ipTM) score were 0.60 and 0.80, respectively and had a ranking score of 0.82. For MdtE and* $Mg^{2+}$ *the pTM score and the ipTM score were 0.62 and 0.83 respectively and had a ranking score of 0.85. For MdtA and* $Mg^{2+}$ *the pTM score and the ipTM score were 0.55 and 0.74 respectively and had a ranking score of 0.81. For MexA and* $Mg^{2+}$ *the pTM score and the ipTM score were 0.62 and 0.79 respectively and had a ranking score of 0.83. For CusB and* $Cu^{2+}$ *the pTM score and the ipTM score were 0.49 and 0.73 respectively and had a ranking score of 0.75.*

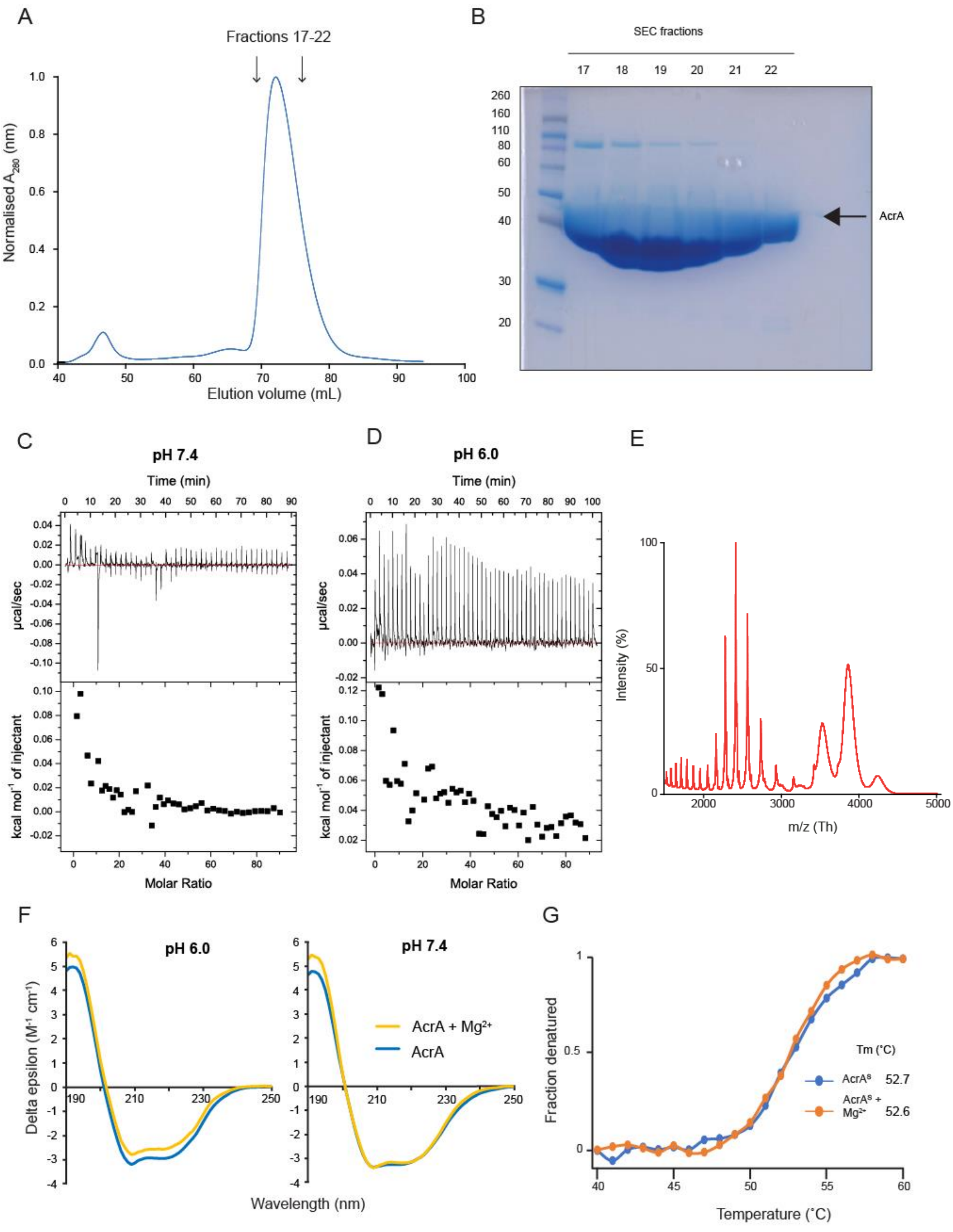

***Figure S4. Biophysical characterisation of AcrA and magnesium. A.*** *Characteristic size exclusion chromatogram (SEC) for AcrA*$^{S}$*. Fractions 17-22 were taken and run on an SDS-PAGE to assess sample purity. Void peak is shown at ~40 mL and the main peak elutes at ~75 mL, suggesting a homogenous monomer.* ***B.*** *Characteristic SDS-PAGE showing the eluted SEC fractions.* ***C.*** *Isothermal titration calorimetry of AcrA and magnesium. micro-ITC (Microcal*

*GE Healthcare) was used to titrate 1.2 mg/ml AcrA with 6.0 mM $MgCl_2$ at 25 °C at pH 7.4.* ***D.*** *Isothermal titration calorimetry repeated at pH 6.0.* ***E****. Native MS of $AcrA^S$ and $Mg^{2+}$. Native-MS characterisation of $AcrA^S$ at pH 6.0 with 100 µM $MgCl_2$. Proteins buffer was exchanged in 100 mM ammonium acetate buffer prior to MS. Two monomeric CSD's observed for $AcrA^S$ + ± $MgCl_2$.* ***F.*** *Full length scans of AcrA +/- ± $MgCl_2$ at pH 6.0 and pH 7.4. Only 190-250 nm analysed due to noisy data at the ends. Proteins loaded at 1 mg/mL and ± $MgCl_2$ at 1 mM. Each scan was repeated in quadruplet, and the averaged scan was analysed. Delta epsilon conversion completed by BeStSel online program[1].* ***G.*** *Circular dichroism thermal melts of $AcrA^S$ ± $MgCl_2$. Scans at 222 nm were taken between 40-60 ˚C with 1 ˚C increments of AcrA ± ± $MgCl_2$ at pH 6.0. Proteins loaded at 0.0075 mg/mL and ± $MgCl_2$ at 1 mM. Plot shows fraction denatured vs temperature. $T_m$'s reported are 52.7 ˚C for $AcrA^S$ and 53.6 for $AcrA^S$ + ± $MgCl_2$.*

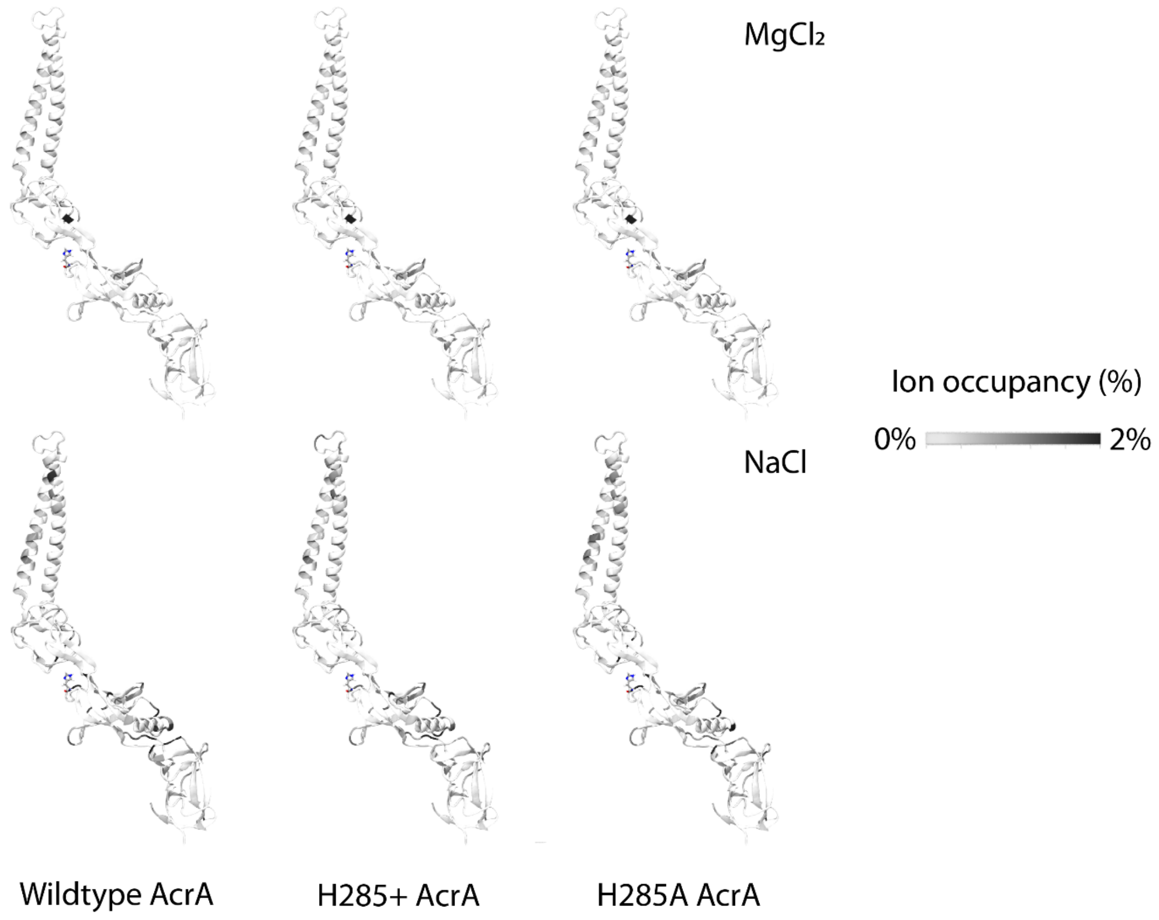

***Figure S5. Ion occupancy (within 7 Å) for $Mg^{2+}$ and $Na^{+}$ on AcrA from MD simulations.*** *Ion occupancy per residue averaged over 4 replicas.*

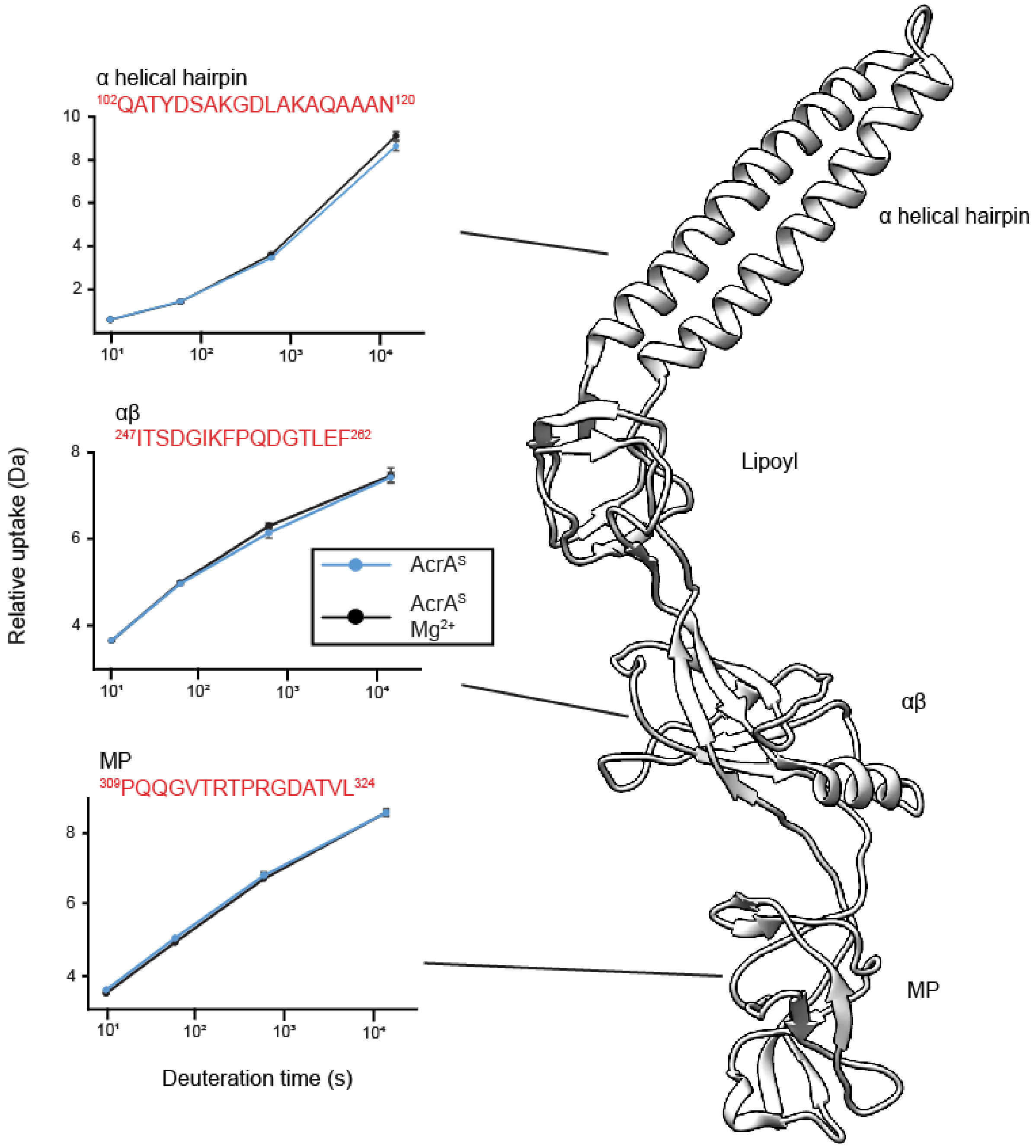

***Figure S6. Structural dynamics of AcrA in the presence of MgCl₂ at pH 7.4.*** *The differential HDX (ΔHDX) plots for ((AcrA$^S$ + Mg$^{2+}$) - AcrA$^S$), at pH 7.4 for the 10-min time point painted onto the structure of AcrA (PDB: 5066) using HDeXplosion and Chimera.[2,3] White signifies areas with no significant change in HDX. Significance was defined to be ≥ 0.5 Da change with a P-value ≤ 0.01 in a Welch's t-test (n=4). Uptake plots for three peptides in different domains of AcrA. Uptake plots are the average deuterium uptake and error bars indicate the standard deviation. Due to the experiment being at pH 7.4, it was more practicably feasible to collect a longer 4-hour time point to see if Mg$^{2+}$ had an effect on dynamics over a longer period of time; again, there were no observed peptides that saw a change in uptake at this time point.*

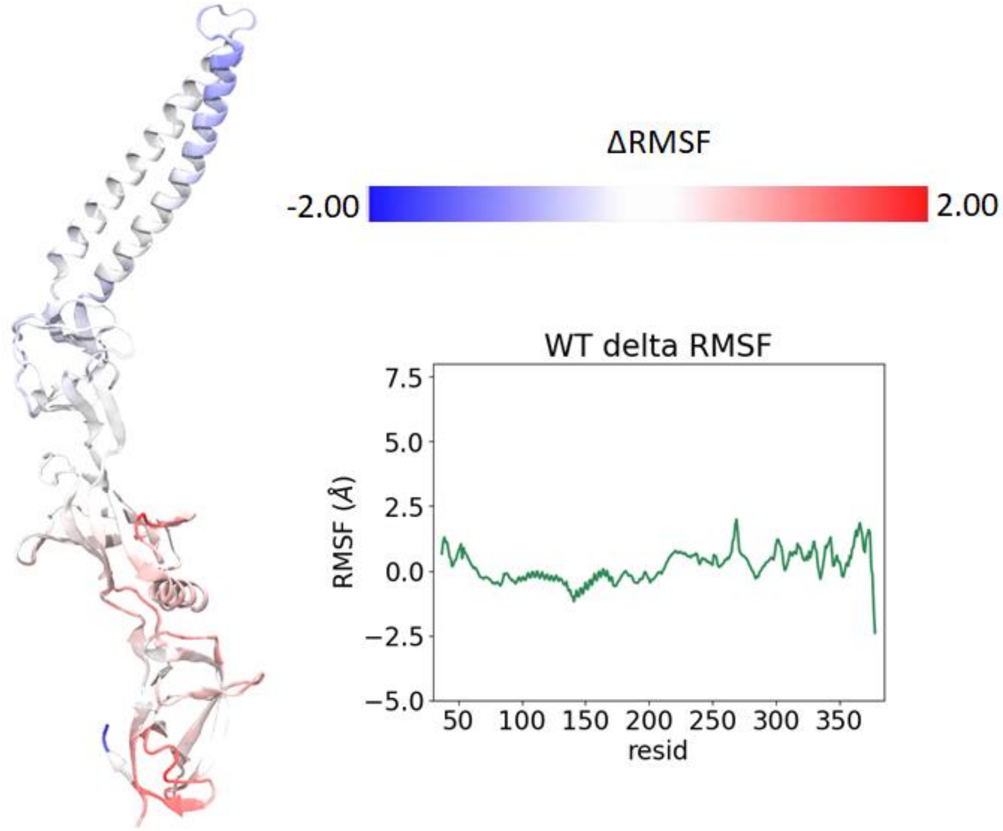

***Figure S7. RMSF from MD simulations of AcrA^S and MgCl₂ at neutral conditions.*** *AcrA coloured according to the difference in root-mean-square fluctuations (RMSF) between simulations of the AcrA$^S$ wild-type state without $MgCl_2$ and with $MgCl_2$ (AcrA$^S$ WT $MgCl_2$ -AcrA$^S$ WT), averaged over four replicas for each. Red indicates that the RMSF has increased whereas blue indicates it has decreased. RMSF was calculated over the last 70 ns of each 100-ns simulation.*

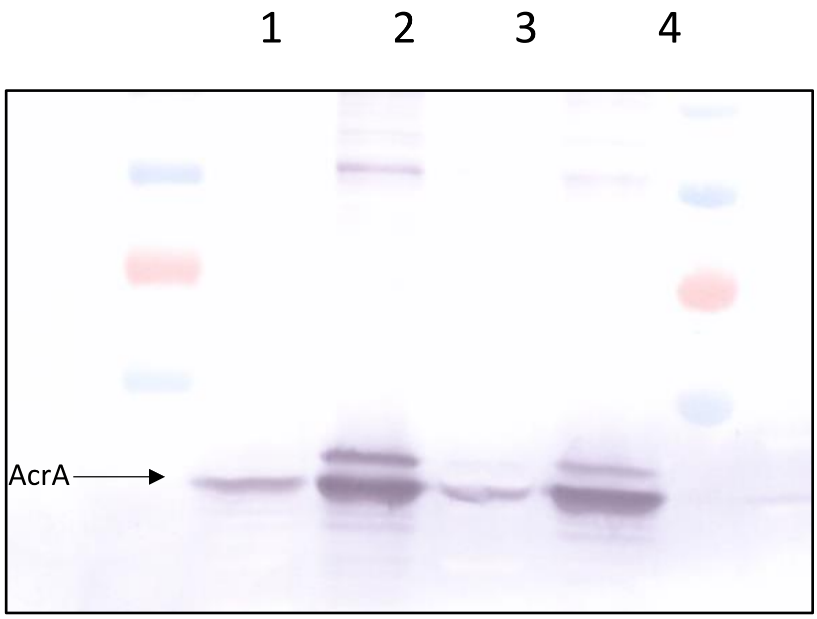

1. E. coli Δ9-pore_pUC18
2. E. coli Δ9-pore_p151
3. E. coli Δ9-pore_p151_AcrA$_{H285A}$ C1
4. E. coli Δ9-pore_p151_AcrA$_{H285A}$ C2

***Figure S8. Protein expression of AcrA mutant.*** *AcrA can be seen for all conditions as labelled on the gel.*

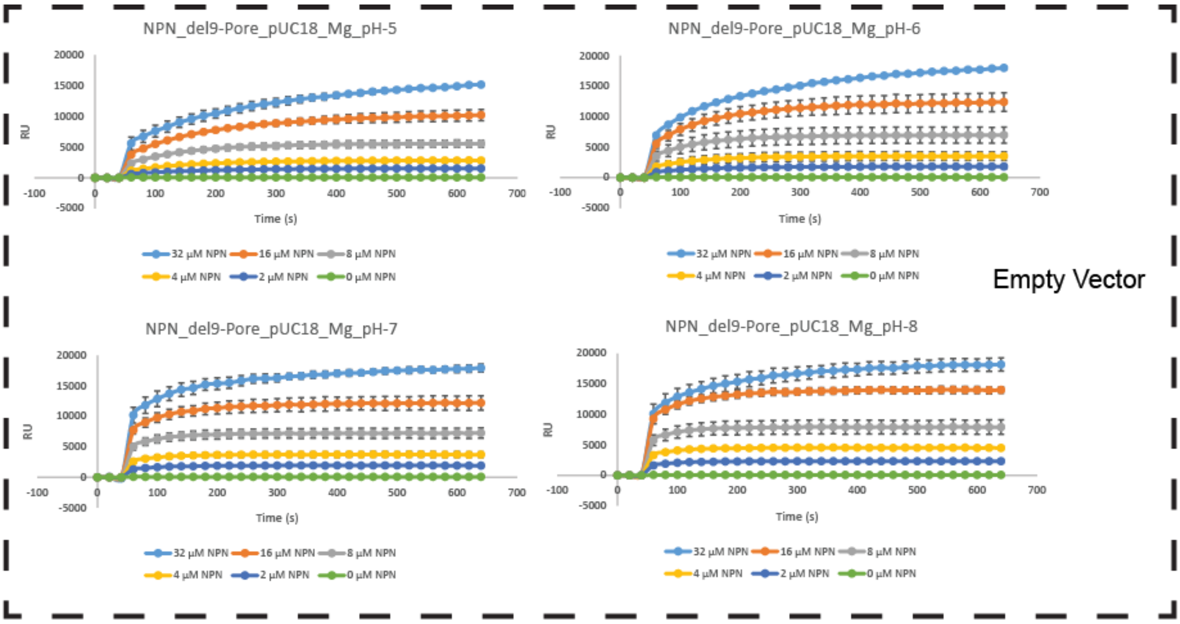
NPN_del9-Pore_pUC18_Mg_pH-5
NPN_del9-Pore_pUC18_Mg_pH-6
NPN_del9-Pore_pUC18_Mg_pH-7
NPN_del9-Pore_pUC18_Mg_pH-8
RU
Time (s)
32 μM NPN 16 μM NPN 8 μM NPN
4 μM NPN 2 μM NPN 0 μM NPN
Empty Vector

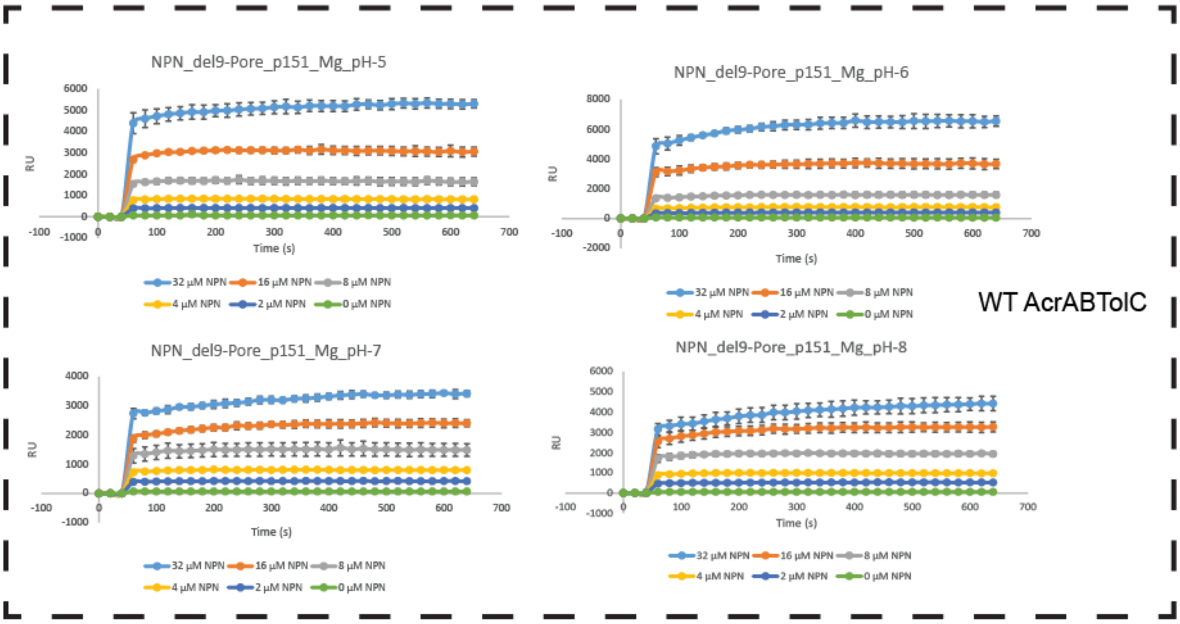
NPN_del9-Pore_p151_Mg_pH-5
NPN_del9-Pore_p151_Mg_pH-6
NPN_del9-Pore_p151_Mg_pH-7
NPN_del9-Pore_p151_Mg_pH-8
RU
Time (s)
32 μM NPN 16 μM NPN 8 μM NPN
4 μM NPN 2 μM NPN 0 μM NPN
WT AcrABTolC

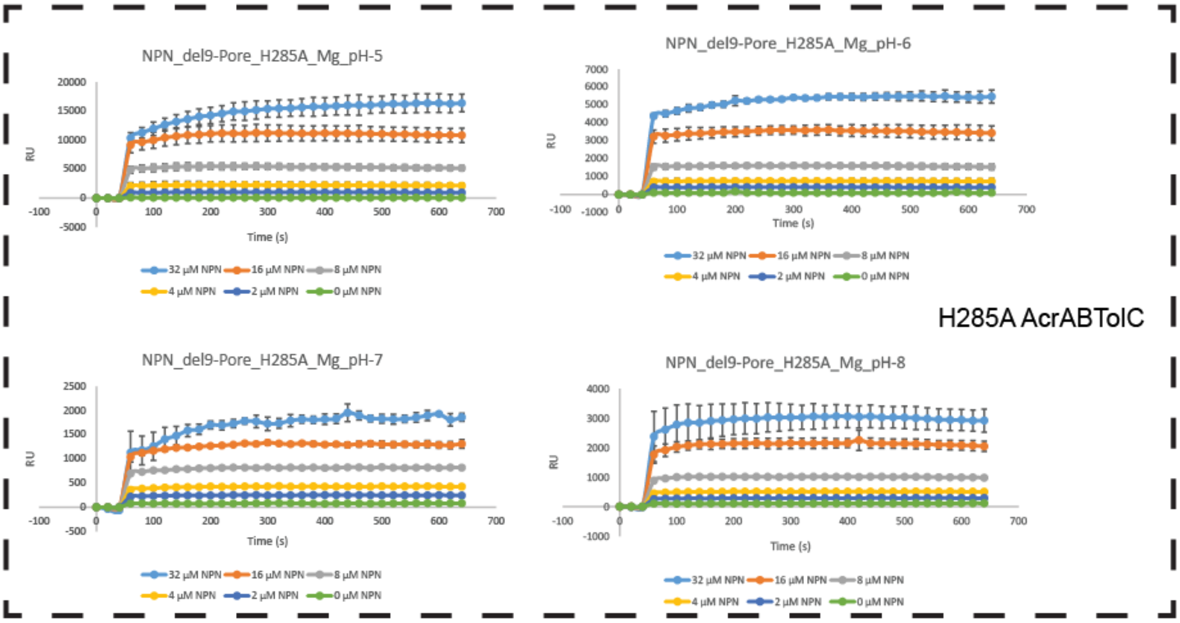
NPN_del9-Pore_H285A_Mg_pH-5
NPN_del9-Pore_H285A_Mg_pH-6
NPN_del9-Pore_H285A_Mg_pH-7
NPN_del9-Pore_H285A_Mg_pH-8
RU
Time (s)
32 μM NPN 16 μM NPN 8 μM NPN
4 μM NPN 2 μM NPN 0 μM NPN
H285A AcrABTolC

***Figure S9. Time course of intracellular NPN accumulation in Δ9-Pore cells.*** *Experiments repeated at pH 5.0, 6.0, 7.0 and 8.0 for three conditions; empty pUC18 vector, AcrAB-TolC and* $AcrA^{H285A}B$*-TolC.*

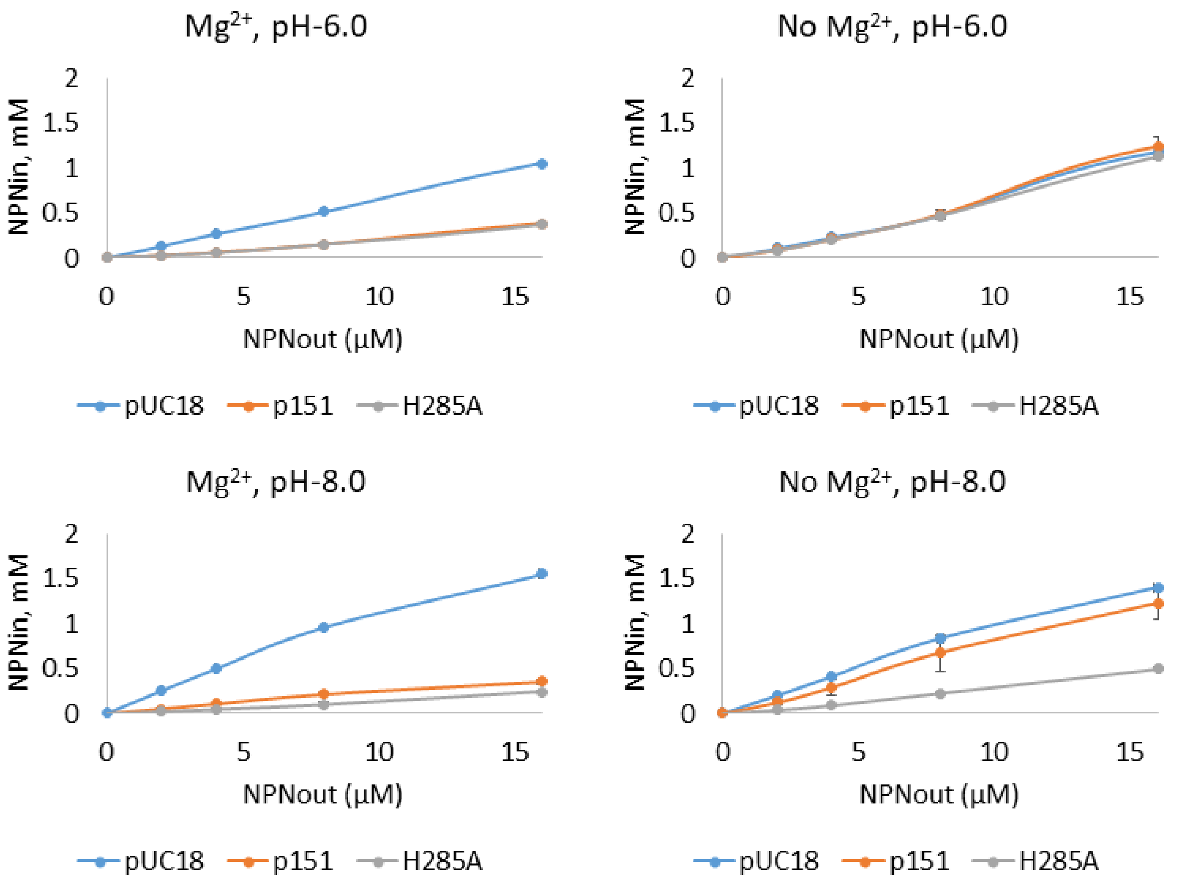

***Figure S10. Effect of $MgCl_2$ on the NPN efflux activity of AcrAB-TolC and its AcrA H285A variant.*** *Steady-state accumulation levels of the fluorescent probe NPN in E. coli Δ9(Pore) cells carrying a AcrAB-TolC or AcrA$^{H285A}$*B-TolC *as a function of the external NPN concentration. Experiments completed ± $MgCl_2$ at pH 6.0 and pH 8.0.*

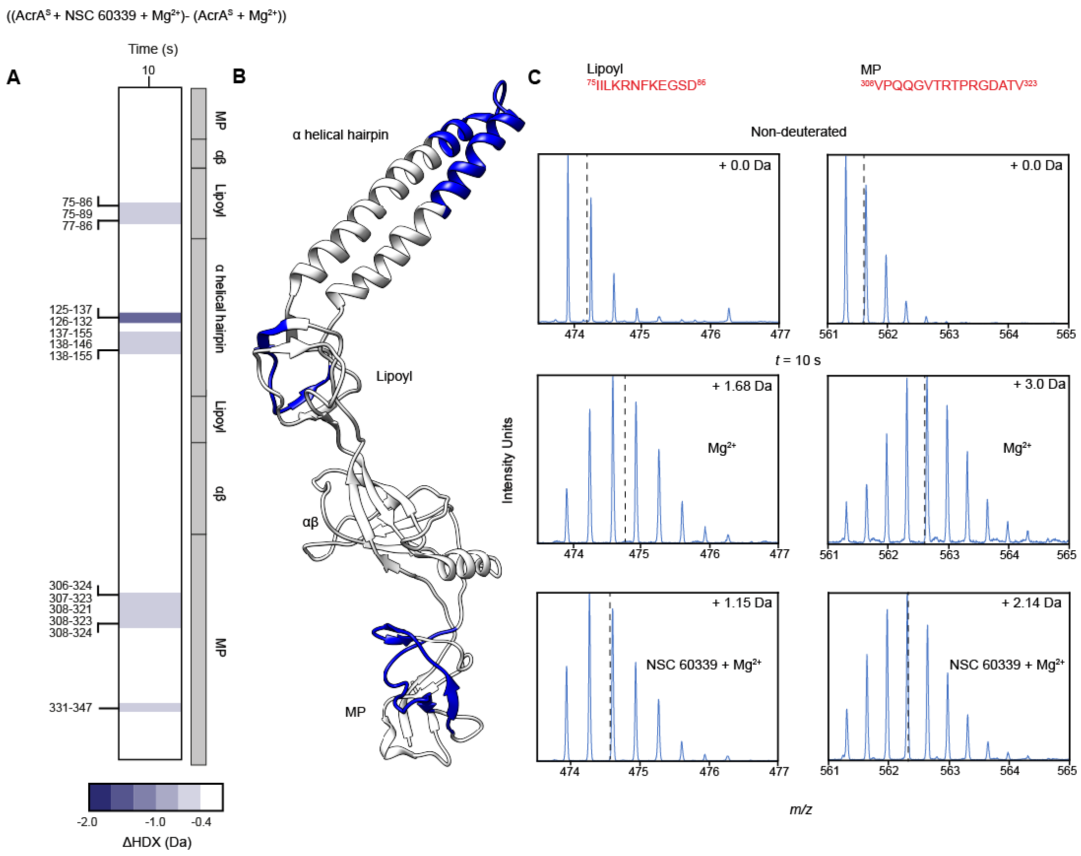

***Figure S11. Inhibition of AcrA by NSC 60339 in the presence of magnesium. A.*** *Chiclet plot displaying the differential HDX (ΔHDX) plots for ((AcrA$^S$ + Mg$^{2+}$) - AcrA$^S$), at pH 6,0 the single time point collected. Blue signifies areas with decreased HDX between states and white signifies areas with no significant change in HDX. Significance was defined to be ≥ 0.4 Da change with a P-value ≤ 0.01 in a Welch's t-test (n=4), as previously reported.*[4] ***B.*** *ΔHDX for ((AcrA$^S$ + Mg$^{2+}$) - AcrA$^S$) for the only time point is painted onto the AcrA structure (PDB:5O66) using HDeXplosion and Chimera.*[2,3] ***C. T****he m/z spectrum for peptides 75-86 and 308-323 under non-deuterating conditions and deuterating conditions with Mg$^{2+}$ and Mg$^{2+}$ + NSC 60339. The centroid is represented by the dotted line, and the mass change of the deuterated samples is written in Daltons.*

*Table S1. BeStSel calculations of AcrA secondary structure*

| Secondary structure | AcrA pH 7.4 (%) | AcrA + $Mg^{2+}$ pH 7.4 (%) | AcrA pH 6.0 (%) | AcrA + $Mg^{2+}$ pH 6.0 (%) |
|---|---|---|---|---|
| **Helix** | 20.4 | 26.7 | 19.8 | 15.6 |
| **Antiparallel** | 26.1 | 23.5 | 30.8 | 35.9 |
| **Parallel** | 2.2 | 0 | 0 | 0 |
| **Turn** | 12.9 | 13.9 | 15.7 | 16.6 |
| **Others** | 38.4 | 35.8 | 33.7 | 31.8 |

*Table S2. Ion occupancy simulations of $Mg^{2+}$ and $Na^{+}$ within 7Å of residue H285*

| **Condition** | **Average number of for Mg2+ or Na+ within 7Å of residue H285** | **Standard deviation** | **Average residence time for $Mg^{2+}$ or $Na^{+}$ within 7Å of residue H285 (ns)** | **Standard deviation** | ***Trans* population** | ***Cis* population** | **Standard deviation** |
|---|---|---|---|---|---|---|---|
| WT AcrA (pH 7.0) NaCl | 0.33 | 0.05 | 1.49 | 0.87 | 0.32 | 0.68 | 0.20 |
| WT AcrA (pH 7.0) $MgCl_2$ | 0.75 | 0.07 | 4.40 | 5.25 | 0.40 | 0.60 | 0.25 |
| HSP AcrA (pH6.0) NaCl | 0.18 | 0.03 | 1.31 | 0.52 | 0.65 | 0.35 | 0.26 |
| HSP AcrA (pH 6.0) $MgCl_2$ | 0.36 | 0.13 | 1.92 | 1.52 | 0.45 | 0.55 | 0.38 |
| H285A AcrA NaCl | 0.20 | 0.06 | 1.36 | 0.70 | 0.54 | 0.46 | 0.28 |
| H285A AcrA $MgCl_2$ | 0.51 | 0.13 | 2.63 | 2.98 | 0.61 | 0.39 | 0.40 |

***pH not modelled outright, pH 6.0 mimicked by doubly protonating His285 (HSP)***

*Table S3. Antibiotic susceptibility assay*

| **Strains** | **SDS (µg/ml)** | **NOV (µg/ml)** | **ERY (µg/ml)** | **VAN (µg/ml)** |
|---|---|---|---|---|
| E. coli Δ9-pore_pUC18 | 7.8 | ≤0.25 | ≤0.25 | 8 |
| E. coli Δ9-pore_p151AcrAB$^{his}$ | ≥1000 | 32 | 4-8 | 8-16 |
| E. coli Δ9-pore_p151_AcrA$_{H285A}$ B$^{his}$ | >1000 | 32 | 8 | 16 |

*Table S4. MD systems simulated*

| Conformation | Ionization | His285 | pH | Time (ns) | # of replicas |
|---|---|---|---|---|---|
| AcrA *cis* | 0.15 M NaCl | Singly protonated His285 | 7 | 500 | 2 |
| AcrA *trans* | 0.15 M NaCl | Singly protonated His285 | 7 | 500 | 2 |
| AcrA *cis* | 0.15 M $MgCl_2$ | Singly protonated His285 | 7 | 500 | 2 |
| AcrA *trans* | 0.15 M $MgCl_2$ | Singly protonated His285 | 7 | 500 | 2 |
| AcrA *cis* | 0.15 M NaCl | Doubly protonated His285 | 6 | 500 | 2 |
| AcrA *trans* | 0.15 M NaCl | Doubly protonated His285 | 6 | 500 | 2 |
| AcrA *cis* | 0.15 M $MgCl_2$ | Doubly protonated His285 | 6 | 500 | 2 |
| AcrA *trans* | 0.15 M $MgCl_2$ | Doubly protonated His285 | 6 | 500 | 2 |
| AcrA *cis* | 0.15 M NaCl | His285Ala | 7 | 500 | 2 |
| AcrA *trans* | 0.15 M NaCl | His285Ala | 7 | 500 | 2 |
| AcrA *cis* | 0.15 M $MgCl_2$ | His285Ala | 7 | 500 | 2 |
| AcrA *trans* | 0.15 M $MgCl_2$ | His285Ala | 7 | 500 | 2 |

### References

1. Micsonai, A. *et al.* BeStSel: A web server for accurate protein secondary structure prediction and fold recognition from the circular dichroism spectra. *Nucleic Acids Res.* **46**, 315–322 (2018).

2. Zhang, N., Yu, X., Zhang, X. & Arcy, S. D. Structural bioinformatics HD-eXplosion: visualization of hydrogen – deuterium exchange data as chiclet and volcano plots with statistical filtering. *Bioinformatics.* **37**, 1926–1927 (2021).

3. Pettersen, E. F. *et al.* UCSF Chimera — A Visualization System for Exploratory Research and Analysis. *J. Comput. Chem.* **25**, 1605–1612 (2004).

4. Lewis, B. R. *et al.* Conformational restriction shapes the inhibition of a multidrug efflux adaptor protein. *Nat. Commun.* **14**, 1–14 (2023).
