## Supplementary Data 3 for "Mg^2+^-dependent mechanism of environmental versatility in a multidrug efflux pump": Uptake plots 3.pdf

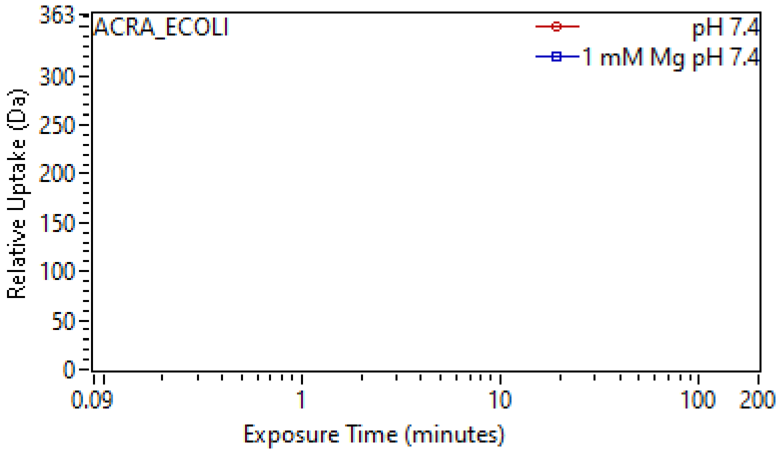
ACRA_ECOLI
pH 7.4
1 mM Mg pH 7.4
Relative Uptake (Da)
363
300
250
200
150
100
50
0
0.09
1
10
100
200
Exposure Time (minutes)

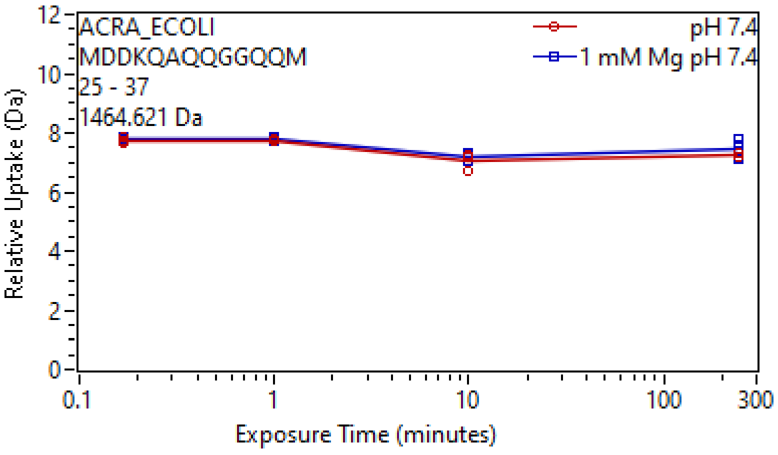
ACRA_ECOLI
MDDKQAQQGGQQM
25 - 37
1464.621 Da
pH 7.4
1 mM Mg pH 7.4
Relative Uptake (Da)
12
10
8
6
4
2
0
0.1
1
10
100
300
Exposure Time (minutes)

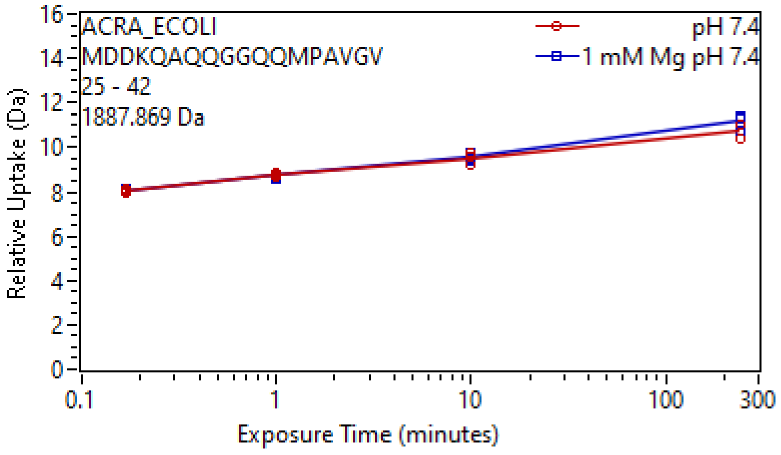
ACRA_ECOLI
MDDKQAQQGGQQMPAVGV
25 - 42
1887.869 Da
pH 7.4
1 mM Mg pH 7.4
Relative Uptake (Da)
16
14
12
10
8
6
4
2
0
0.1
1
10
100
300
Exposure Time (minutes)

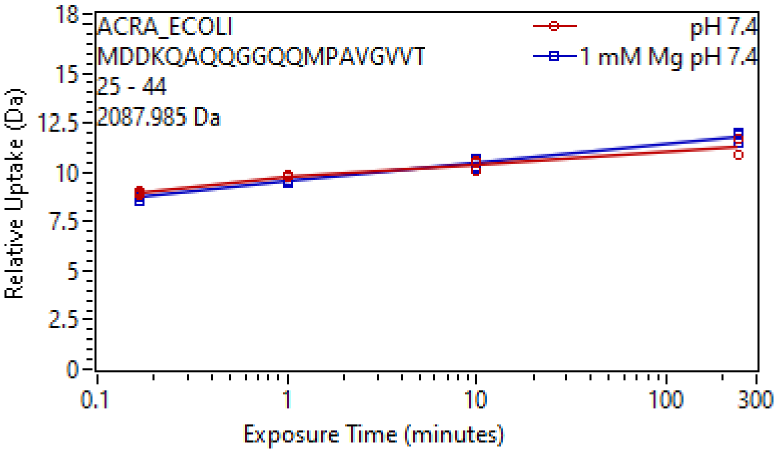
ACRA_ECOLI
MDDKQAQQGGQQMPAVGVVT
25 - 44
2087.985 Da
pH 7.4
1 mM Mg pH 7.4
Relative Uptake (Da)
18
15
12.5
10
7.5
5
2.5
0
0.1
1
10
100
300
Exposure Time (minutes)

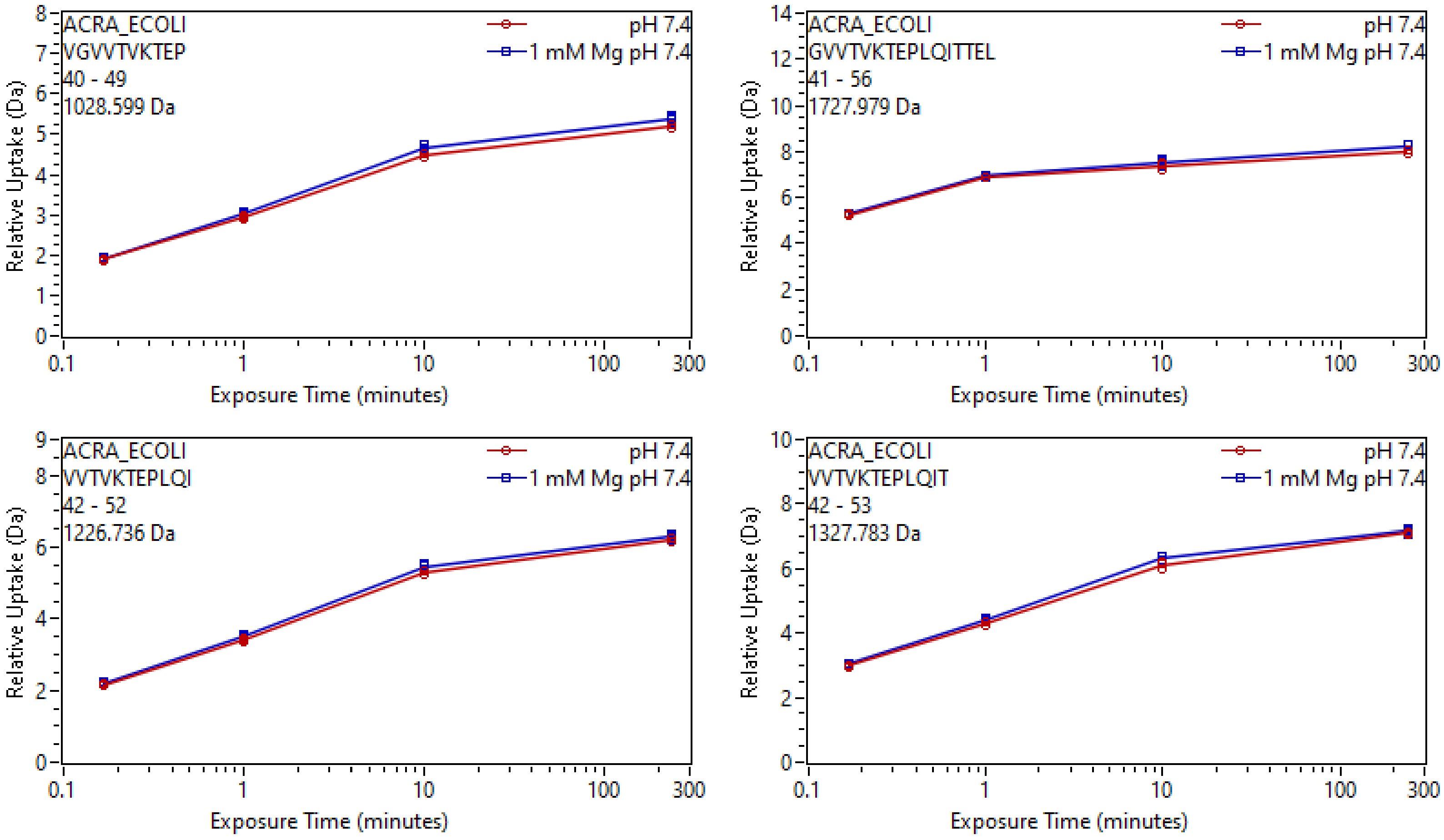

ACRA_ECOLI
VGVVTVKTEP
40 - 49
1028.599 Da
pH 7.4
1 mM Mg pH 7.4
ACRA_ECOLI
GVVTVKTEPLQITTEL
41 - 56
1727.979 Da
ACRA_ECOLI
VVTVKTEPLQI
42 - 52
1226.736 Da
ACRA_ECOLI
VVTVKTEPLQIT
42 - 53
1327.783 Da
Relative Uptake (Da)
Exposure Time (minutes)
0.1
1
10
100
300

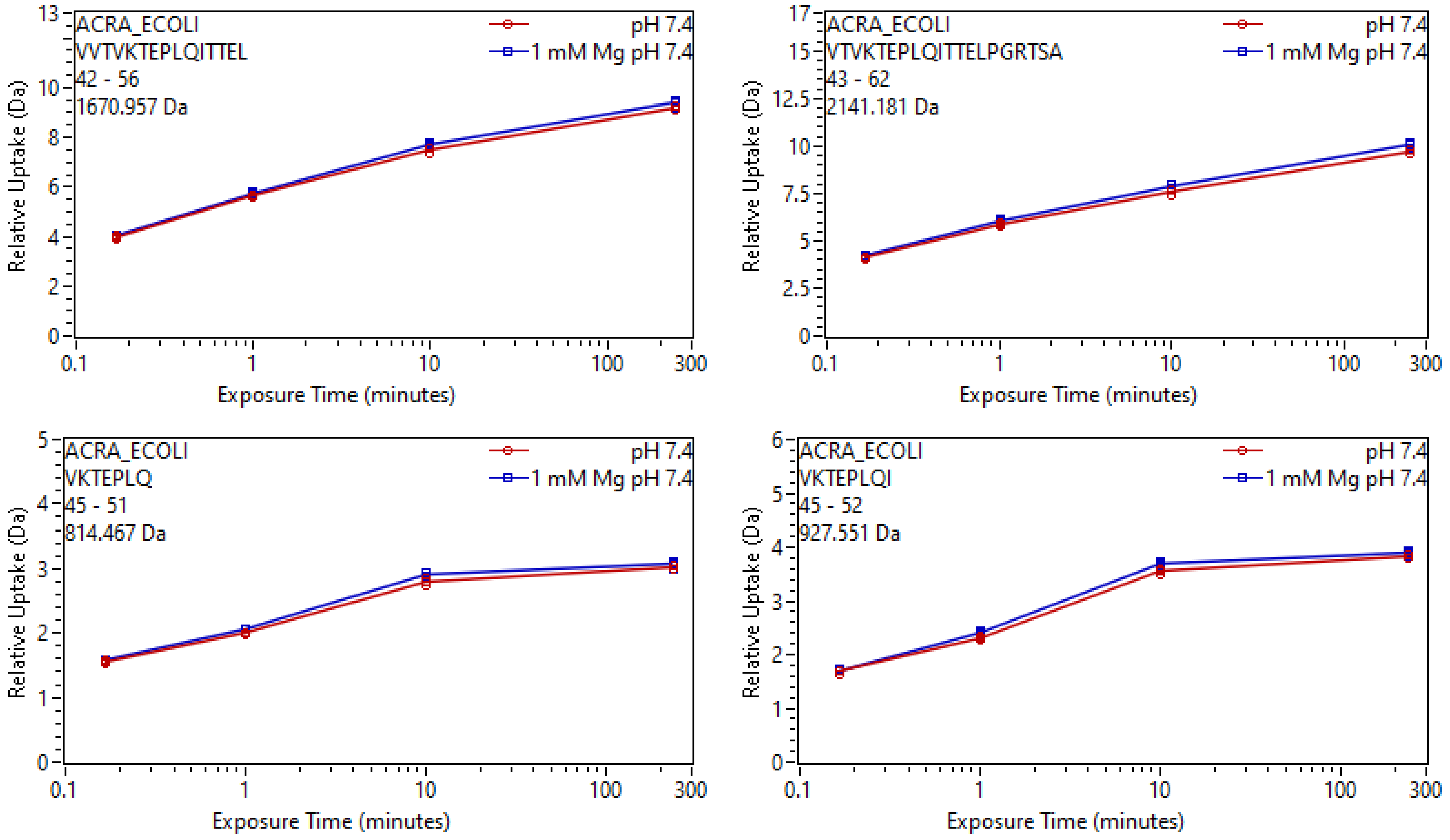
ACRA_ECOLI
VVTVKTEPLQITTEL
42 - 56
1670.957 Da
pH 7.4
1 mM Mg pH 7.4
Relative Uptake (Da)
Exposure Time (minutes)
ACRA_ECOLI
VTVKTEPLQITTELPGRTSA
43 - 62
2141.181 Da
pH 7.4
1 mM Mg pH 7.4
Relative Uptake (Da)
Exposure Time (minutes)
ACRA_ECOLI
VKTEPLQ
45 - 51
814.467 Da
pH 7.4
1 mM Mg pH 7.4
Relative Uptake (Da)
Exposure Time (minutes)
ACRA_ECOLI
VKTEPLQI
45 - 52
927.551 Da
pH 7.4
1 mM Mg pH 7.4
Relative Uptake (Da)
Exposure Time (minutes)

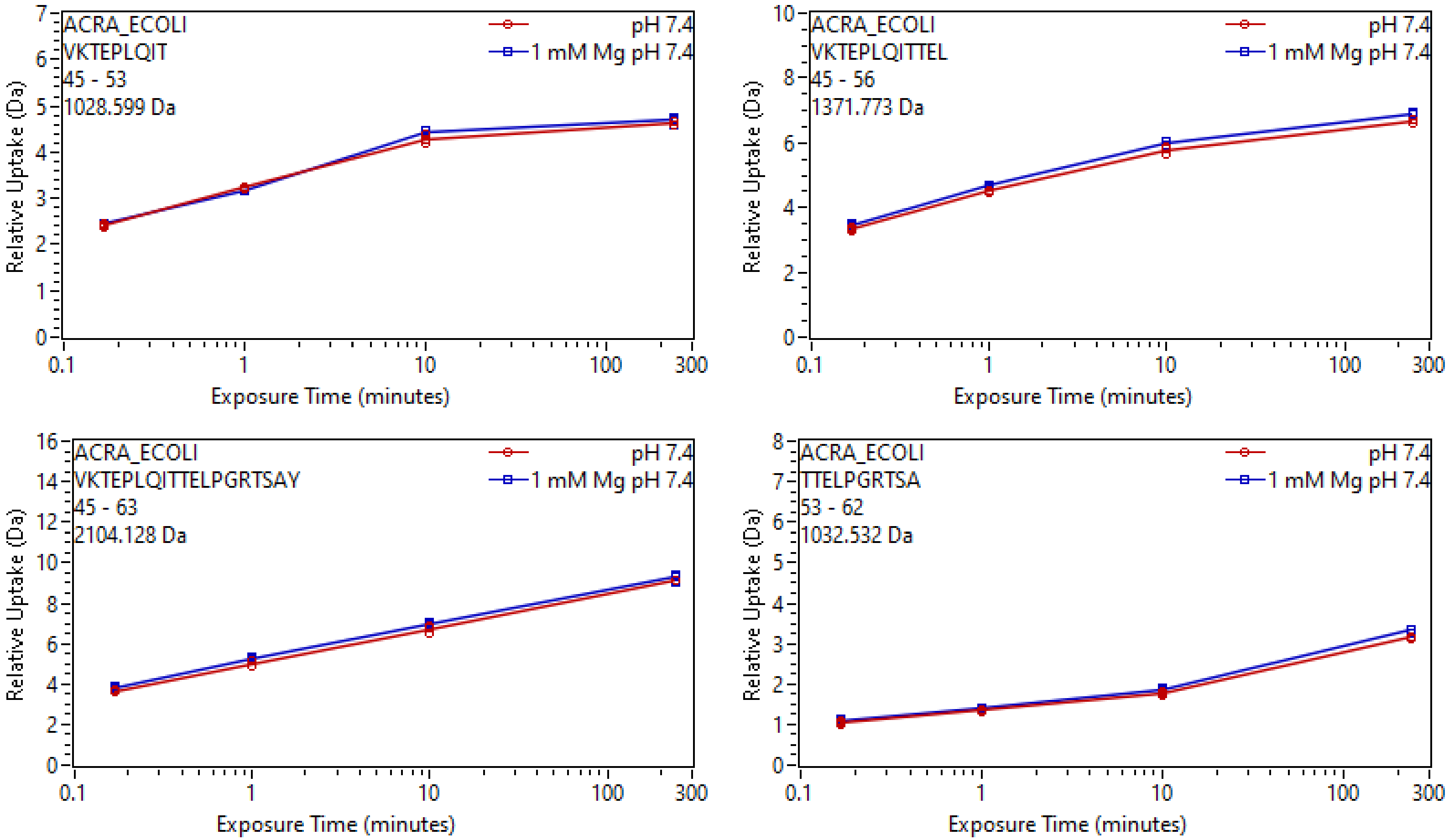

ACRA_ECOLI
VKTEPLQIT
45 - 53
1028.599 Da
pH 7.4
1 mM Mg pH 7.4
Relative Uptake (Da)
Exposure Time (minutes)
ACRA_ECOLI
VKTEPLQITTEL
45 - 56
1371.773 Da
pH 7.4
1 mM Mg pH 7.4
Relative Uptake (Da)
Exposure Time (minutes)
ACRA_ECOLI
VKTEPLQITTELPGRTSAY
45 - 63
2104.128 Da
pH 7.4
1 mM Mg pH 7.4
Relative Uptake (Da)
Exposure Time (minutes)
ACRA_ECOLI
TTELPGRTSA
53 - 62
1032.532 Da
pH 7.4
1 mM Mg pH 7.4
Relative Uptake (Da)
Exposure Time (minutes)

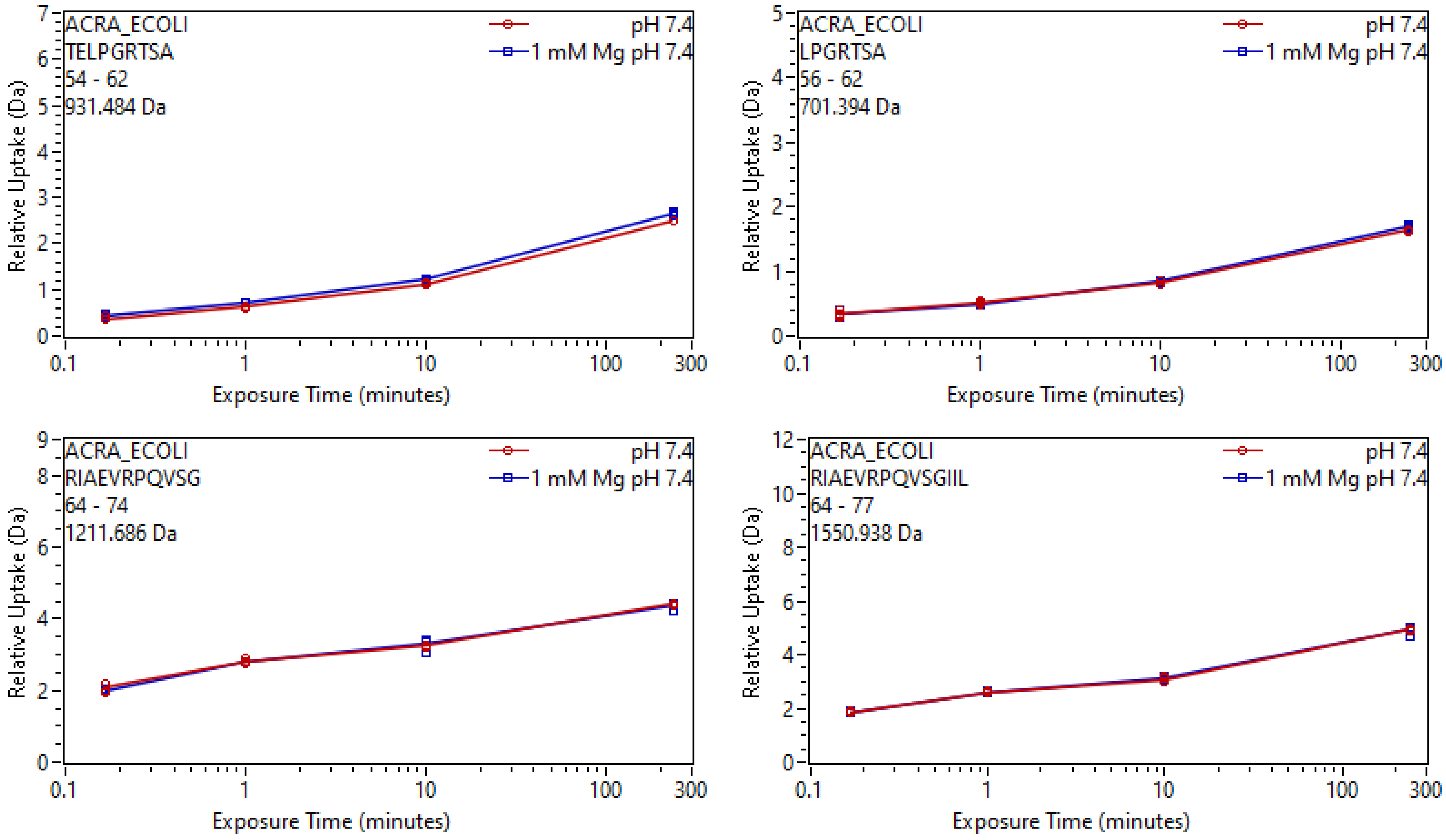

ACRA_ECOLI
TELPGRTSA
54 - 62
931.484 Da
pH 7.4
1 mM Mg pH 7.4
Relative Uptake (Da)
Exposure Time (minutes)
0.1
1
10
100
300
0
1
2
3
4
5
6
7
ACRA_ECOLI
LPGRTSA
56 - 62
701.394 Da
pH 7.4
1 mM Mg pH 7.4
Relative Uptake (Da)
Exposure Time (minutes)
0.1
1
10
100
300
0
1
2
3
4
5
ACRA_ECOLI
RIAEVRPQVSG
64 - 74
1211.686 Da
pH 7.4
1 mM Mg pH 7.4
Relative Uptake (Da)
Exposure Time (minutes)
0.1
1
10
100
300
0
2
4
6
8
9
ACRA_ECOLI
RIAEVRPQVSGIIL
64 - 77
1550.938 Da
pH 7.4
1 mM Mg pH 7.4
Relative Uptake (Da)
Exposure Time (minutes)
0.1
1
10
100
300
0
2
4
6
8
10
12

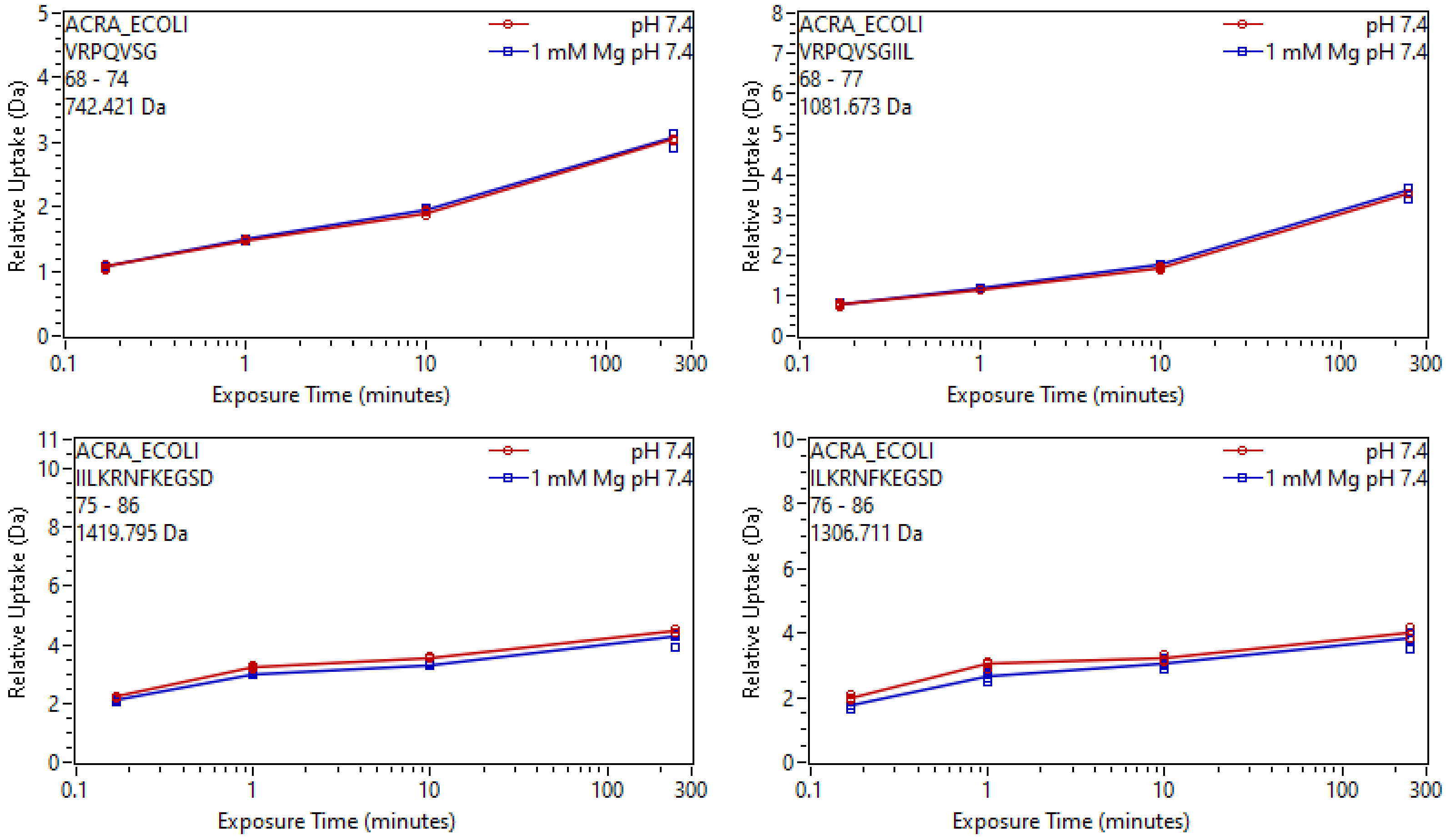

ACRA_ECOLI
VRPQVSG
68 - 74
742.421 Da
pH 7.4
1 mM Mg pH 7.4
Relative Uptake (Da)
Exposure Time (minutes)
0.1
1
10
100
300
ACRA_ECOLI
VRPQVSGIIL
68 - 77
1081.673 Da
pH 7.4
1 mM Mg pH 7.4
Relative Uptake (Da)
Exposure Time (minutes)
ACRA_ECOLI
IILKRNFKEGSD
75 - 86
1419.795 Da
pH 7.4
1 mM Mg pH 7.4
Relative Uptake (Da)
Exposure Time (minutes)
ACRA_ECOLI
ILKRNFKEGSD
76 - 86
1306.711 Da
pH 7.4
1 mM Mg pH 7.4
Relative Uptake (Da)
Exposure Time (minutes)

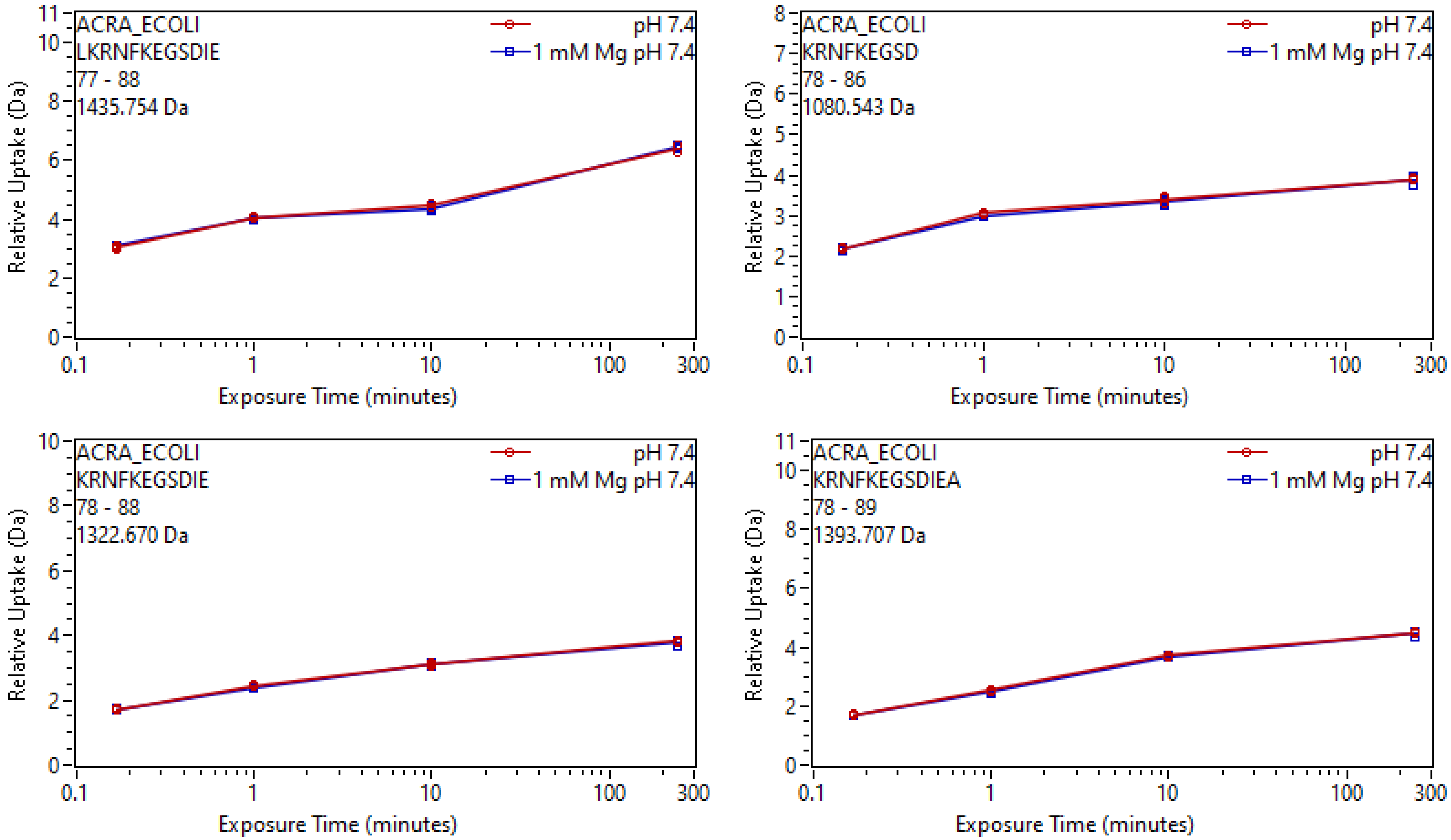
ACRA_ECOLI
LKRNFKEGSDIE
77 - 88
1435.754 Da
pH 7.4
1 mM Mg pH 7.4
Relative Uptake (Da)
Exposure Time (minutes)
ACRA_ECOLI
KRNFKEGSD
78 - 86
1080.543 Da
pH 7.4
1 mM Mg pH 7.4
Relative Uptake (Da)
Exposure Time (minutes)
ACRA_ECOLI
KRNFKEGSDIE
78 - 88
1322.670 Da
pH 7.4
1 mM Mg pH 7.4
Relative Uptake (Da)
Exposure Time (minutes)
ACRA_ECOLI
KRNFKEGSDIEA
78 - 89
1393.707 Da
pH 7.4
1 mM Mg pH 7.4
Relative Uptake (Da)
Exposure Time (minutes)

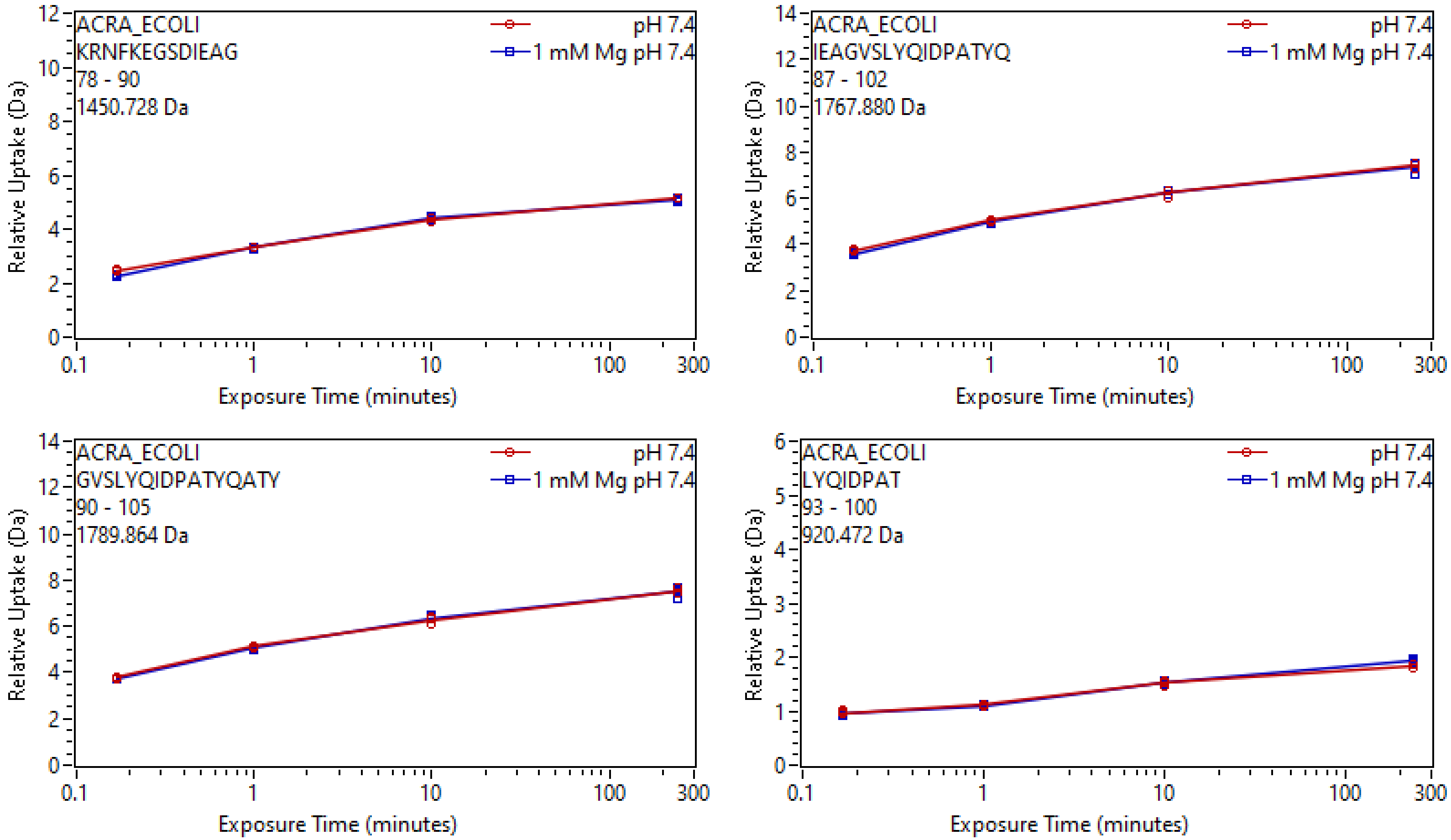
ACRA_ECOLI
KRNFKEGSDIEAG
78 - 90
1450.728 Da
pH 7.4
1 mM Mg pH 7.4
Relative Uptake (Da)
Exposure Time (minutes)
ACRA_ECOLI
IEAGVSLYQIDPATYQ
87 - 102
1767.880 Da
ACRA_ECOLI
GVSLYQIDPATYQATY
90 - 105
1789.864 Da
ACRA_ECOLI
LYQIDPAT
93 - 100
920.472 Da

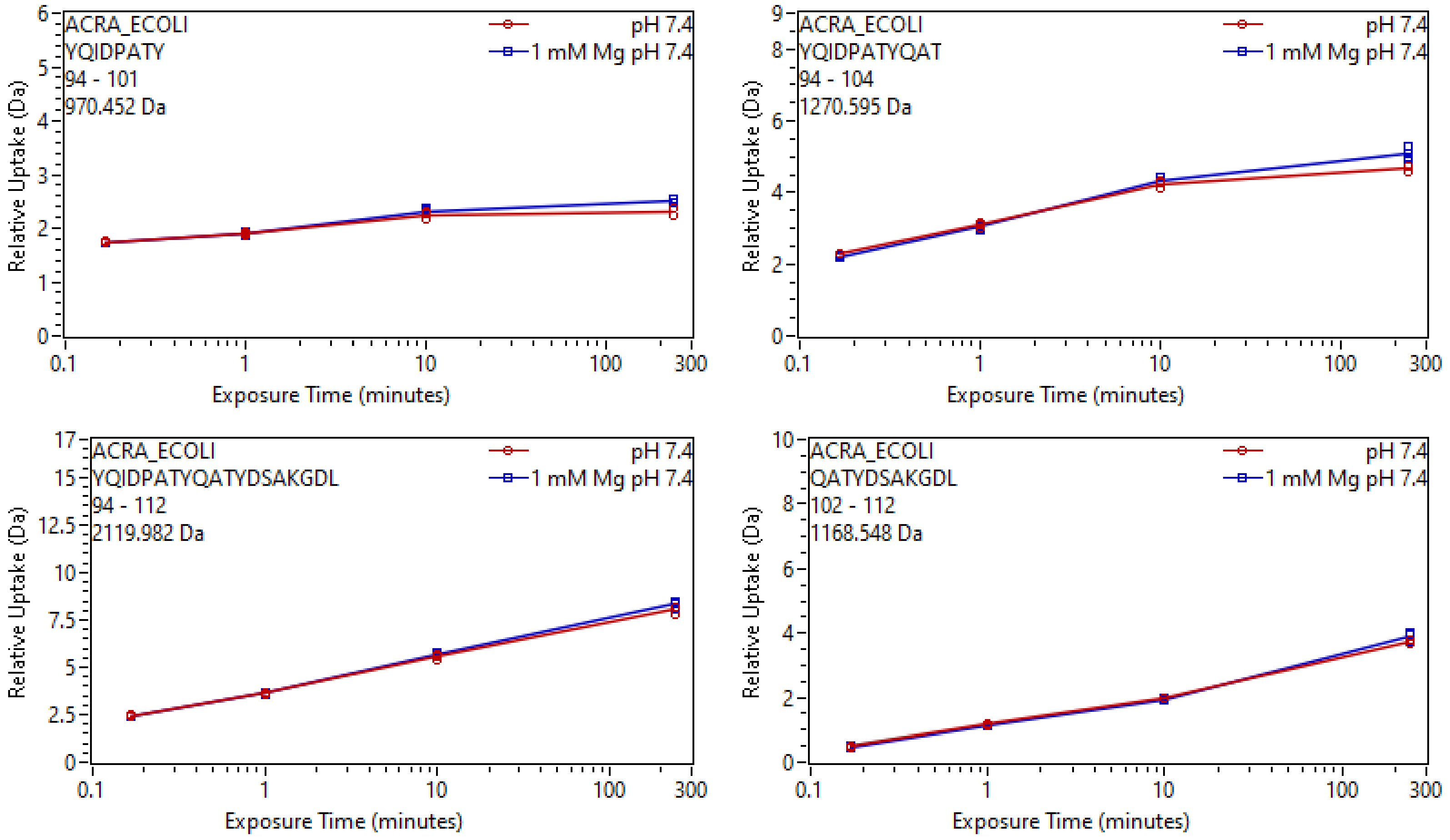

ACRA_ECOLI
YQIDPATY
94 - 101
970.452 Da
pH 7.4
1 mM Mg pH 7.4
Relative Uptake (Da)
Exposure Time (minutes)
ACRA_ECOLI
YQIDPATYQAT
94 - 104
1270.595 Da
pH 7.4
1 mM Mg pH 7.4
Relative Uptake (Da)
Exposure Time (minutes)
ACRA_ECOLI
YQIDPATYQATYDSAKGDL
94 - 112
2119.982 Da
pH 7.4
1 mM Mg pH 7.4
Relative Uptake (Da)
Exposure Time (minutes)
ACRA_ECOLI
QATYDSAKGDL
102 - 112
1168.548 Da
pH 7.4
1 mM Mg pH 7.4
Relative Uptake (Da)
Exposure Time (minutes)

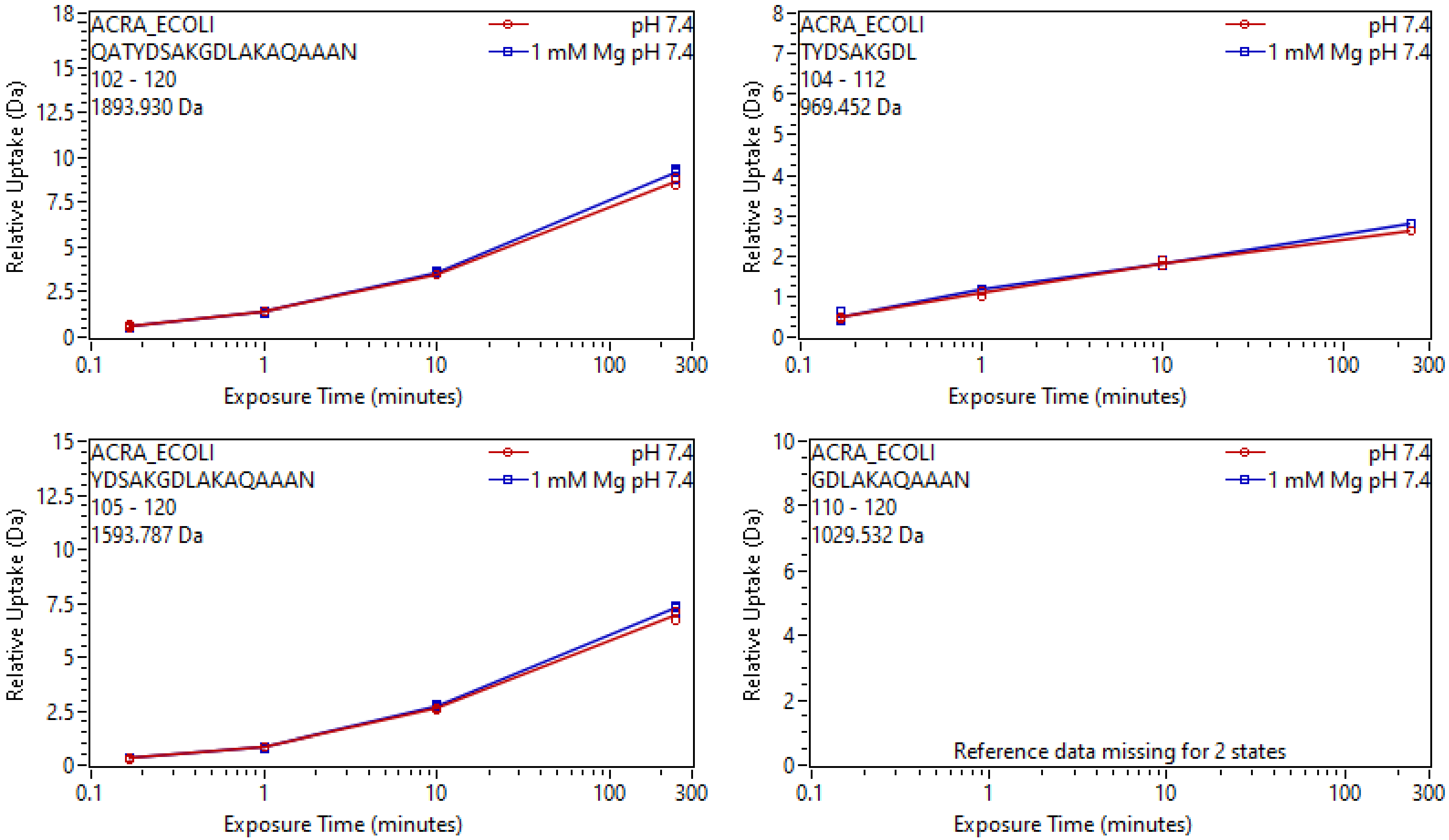
ACRA_ECOLI
QATYDSAKGDLAKAQAAAN
102 - 120
1893.930 Da
pH 7.4
1 mM Mg pH 7.4
Relative Uptake (Da)
Exposure Time (minutes)
ACRA_ECOLI
TYDSAKGDL
104 - 112
969.452 Da
pH 7.4
1 mM Mg pH 7.4
Relative Uptake (Da)
Exposure Time (minutes)
ACRA_ECOLI
YDSAKGDLAKAQAAAN
105 - 120
1593.787 Da
pH 7.4
1 mM Mg pH 7.4
Relative Uptake (Da)
Exposure Time (minutes)
ACRA_ECOLI
GDLAKAQAAAN
110 - 120
1029.532 Da
pH 7.4
1 mM Mg pH 7.4
Reference data missing for 2 states
Relative Uptake (Da)
Exposure Time (minutes)

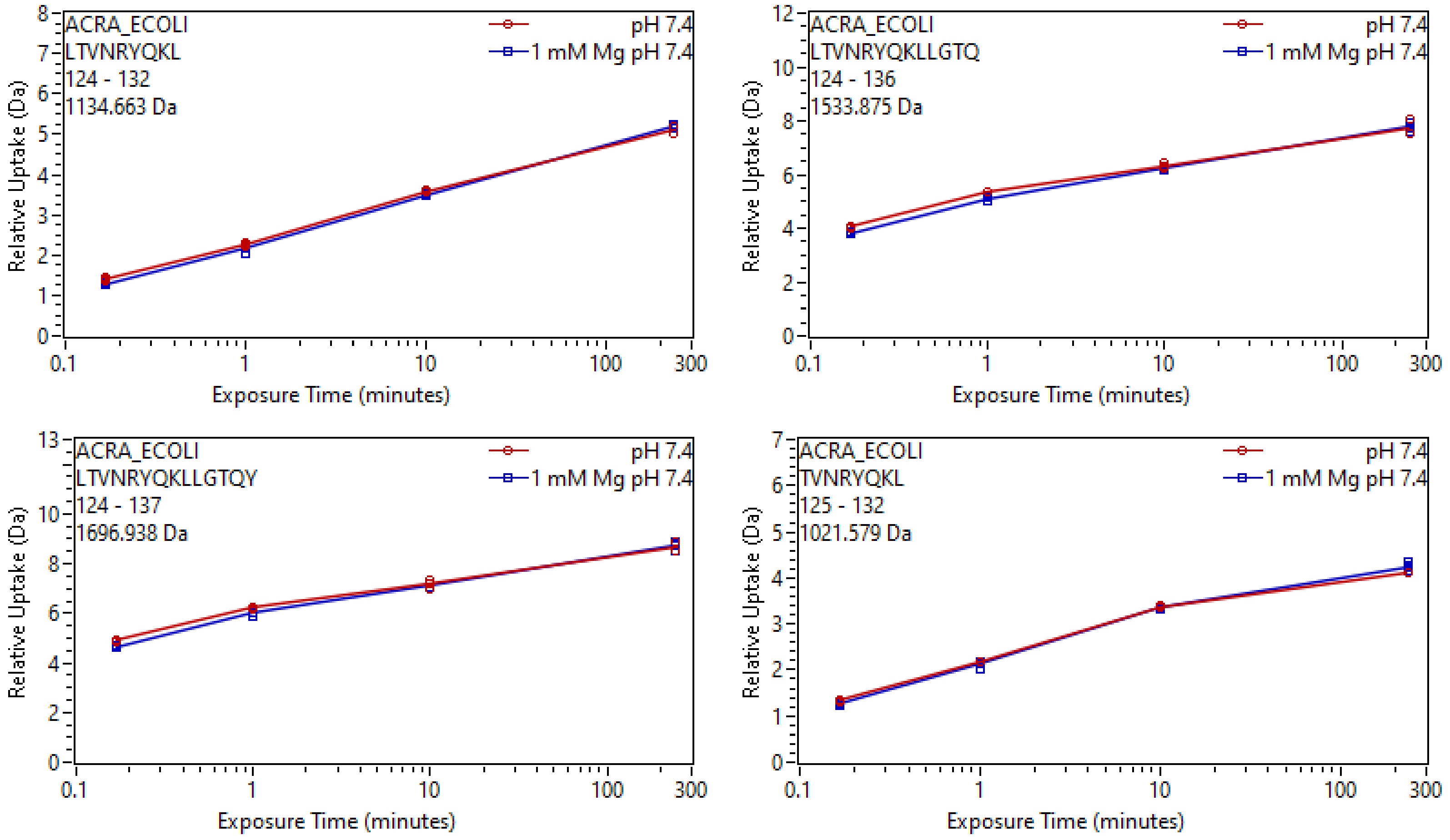
ACRA_ECOLI
LTVNRYQKL
124 - 132
1134.663 Da
pH 7.4
1 mM Mg pH 7.4
Relative Uptake (Da)
Exposure Time (minutes)
ACRA_ECOLI
LTVNRYQKLLGTQ
124 - 136
1533.875 Da
ACRA_ECOLI
LTVNRYQKLLGTQY
124 - 137
1696.938 Da
ACRA_ECOLI
TVNRYQKL
125 - 132
1021.579 Da

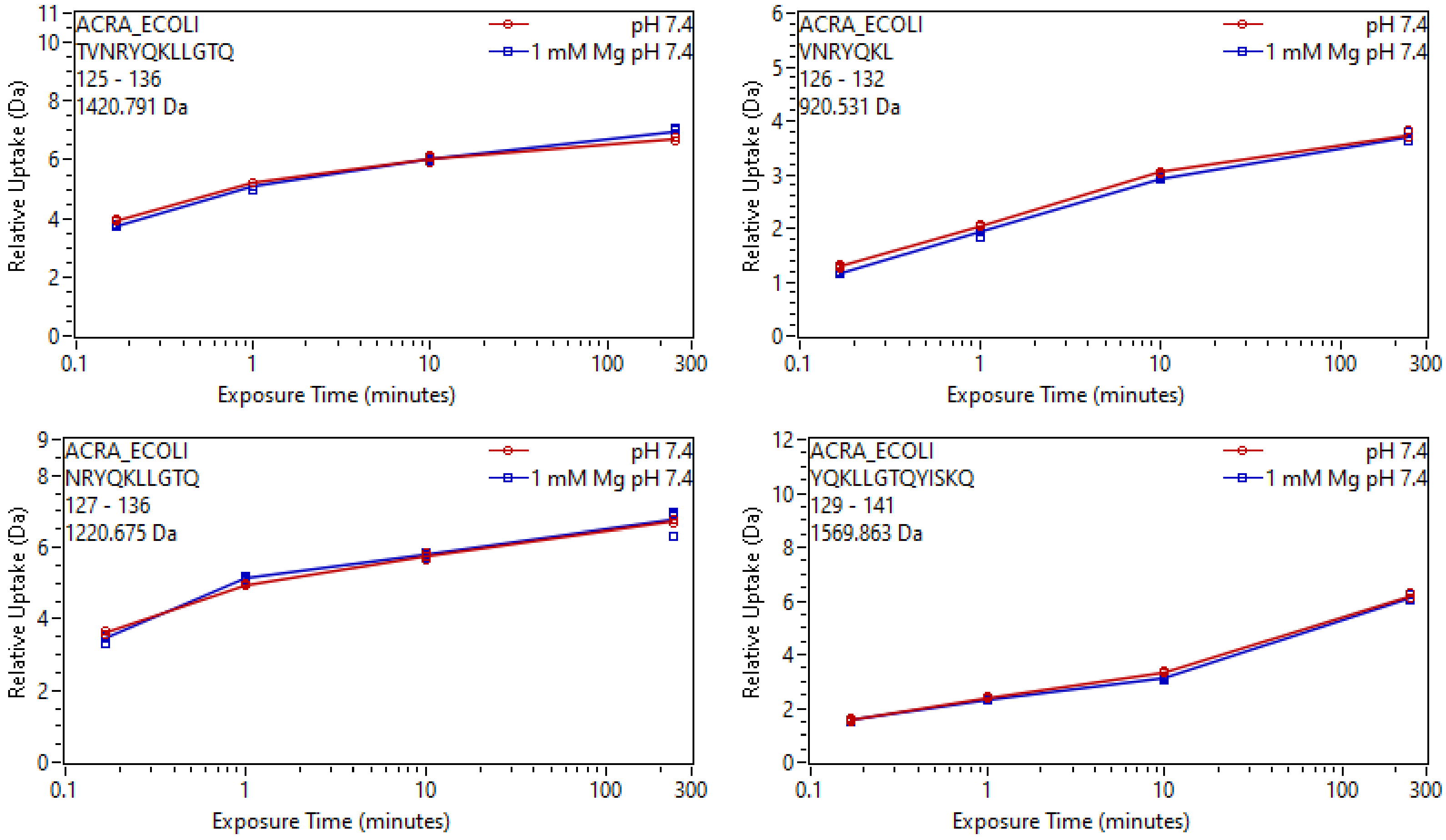
ACRA_ECOLI
TVNRYQKLLGTQ
125 - 136
1420.791 Da
pH 7.4
1 mM Mg pH 7.4
Relative Uptake (Da)
Exposure Time (minutes)
0.1
1
10
100
300
ACRA_ECOLI
VNRYQKL
126 - 132
920.531 Da
pH 7.4
1 mM Mg pH 7.4
Relative Uptake (Da)
Exposure Time (minutes)
ACRA_ECOLI
NRYQKLLGTQ
127 - 136
1220.675 Da
pH 7.4
1 mM Mg pH 7.4
Relative Uptake (Da)
Exposure Time (minutes)
ACRA_ECOLI
YQKLLGTQYISKQ
129 - 141
1569.863 Da
pH 7.4
1 mM Mg pH 7.4
Relative Uptake (Da)
Exposure Time (minutes)

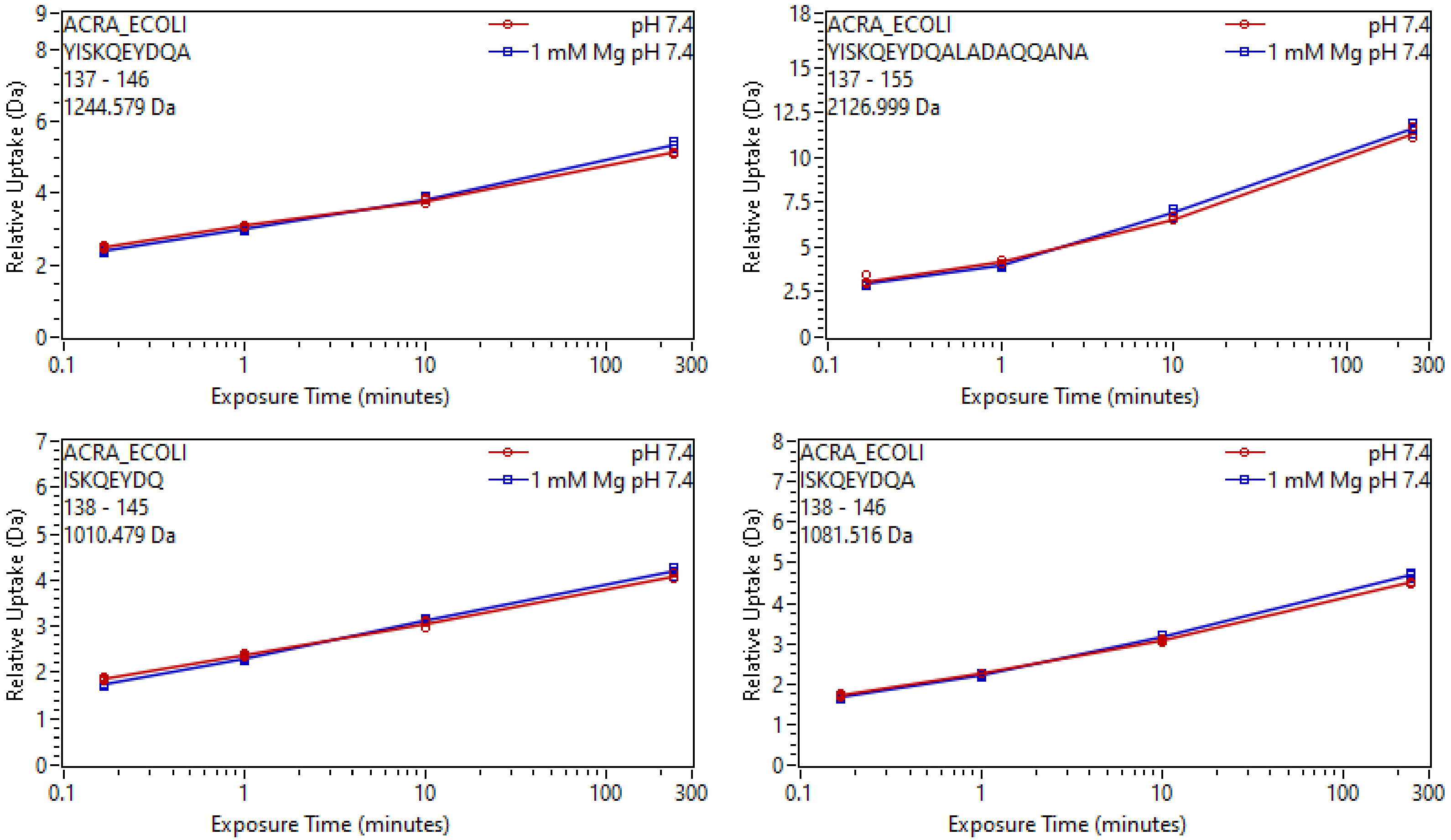
ACRA_ECOLI
YISKQEYDQA
137 - 146
1244.579 Da
pH 7.4
1 mM Mg pH 7.4
Relative Uptake (Da)
Exposure Time (minutes)
ACRA_ECOLI
YISKQEYDQALADAQQANA
137 - 155
2126.999 Da
pH 7.4
1 mM Mg pH 7.4
Relative Uptake (Da)
Exposure Time (minutes)
ACRA_ECOLI
ISKQEYDQ
138 - 145
1010.479 Da
pH 7.4
1 mM Mg pH 7.4
Relative Uptake (Da)
Exposure Time (minutes)
ACRA_ECOLI
ISKQEYDQA
138 - 146
1081.516 Da
pH 7.4
1 mM Mg pH 7.4
Relative Uptake (Da)
Exposure Time (minutes)

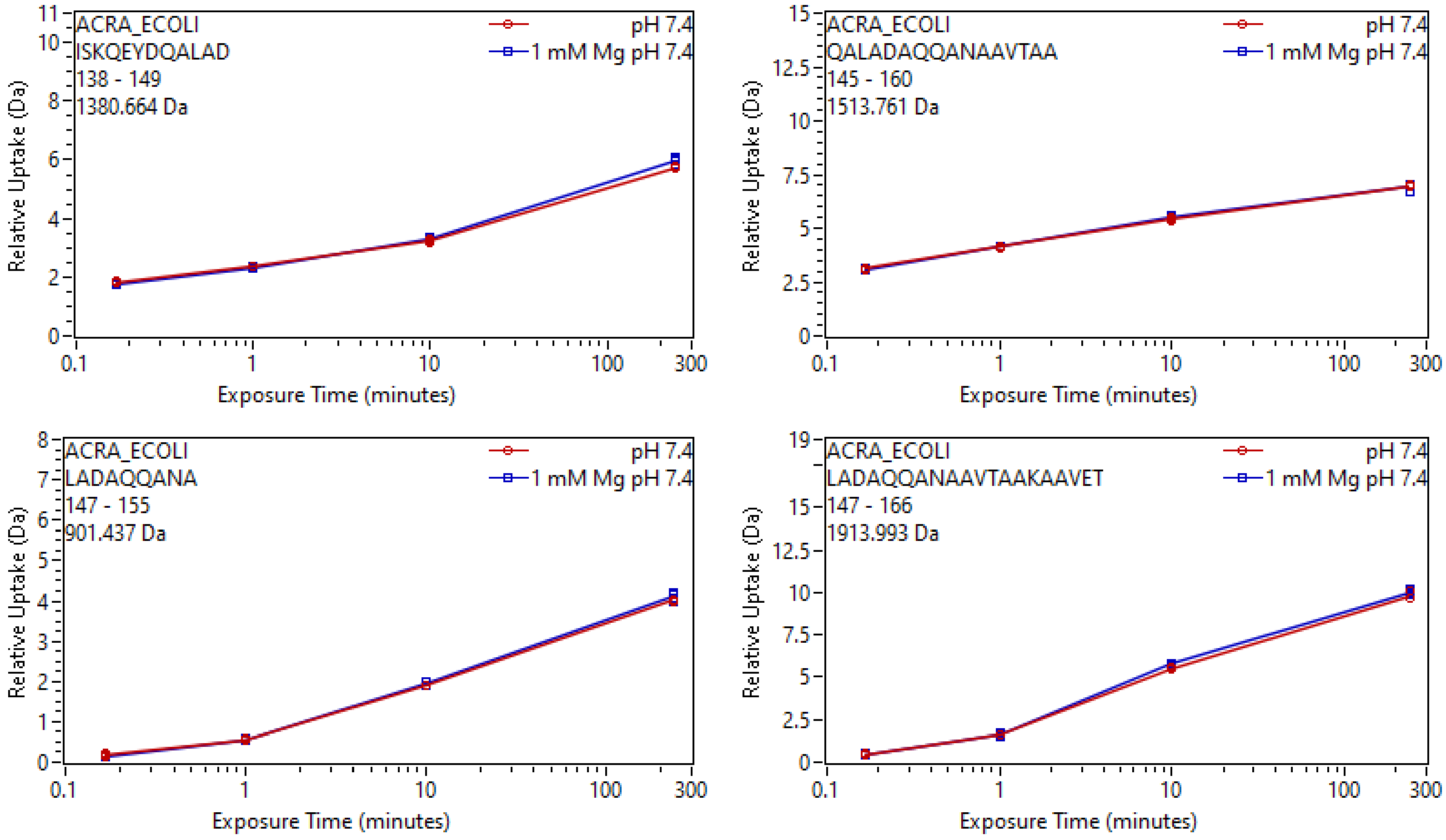
ACRA_ECOLI
ISKQEYDQALAD
138 - 149
1380.664 Da
pH 7.4
1 mM Mg pH 7.4
Relative Uptake (Da)
Exposure Time (minutes)
0.1
1
10
100
300
0
2
4
6
8
10
11
ACRA_ECOLI
QALADAQQANAAVTAA
145 - 160
1513.761 Da
pH 7.4
1 mM Mg pH 7.4
Relative Uptake (Da)
Exposure Time (minutes)
0
2.5
5
7.5
10
12.5
15
ACRA_ECOLI
LADAQQANA
147 - 155
901.437 Da
pH 7.4
1 mM Mg pH 7.4
Relative Uptake (Da)
Exposure Time (minutes)
0
1
2
3
4
5
6
7
8
ACRA_ECOLI
LADAQQANAAVTAAKAAVET
147 - 166
1913.993 Da
pH 7.4
1 mM Mg pH 7.4
Relative Uptake (Da)
Exposure Time (minutes)
0
2.5
5
7.5
10
12.5
15
19

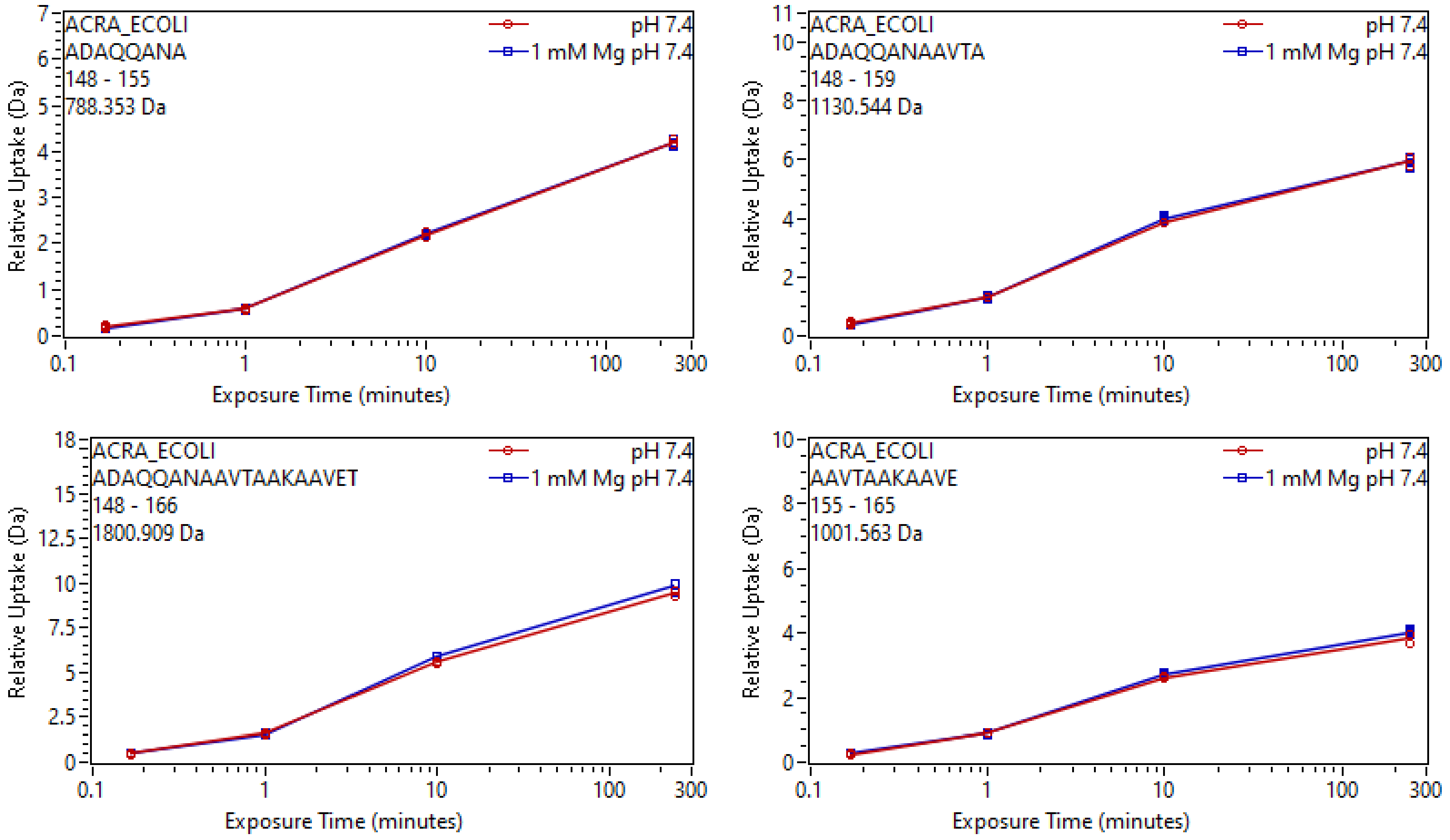

ACRA_ECOLI
ADAQQANA
148 - 155
788.353 Da
pH 7.4
1 mM Mg pH 7.4
Relative Uptake (Da)
Exposure Time (minutes)
ACRA_ECOLI
ADAQQANAAVTA
148 - 159
1130.544 Da
pH 7.4
1 mM Mg pH 7.4
Relative Uptake (Da)
Exposure Time (minutes)
ACRA_ECOLI
ADAQQANAAVTAAKAAVET
148 - 166
1800.909 Da
pH 7.4
1 mM Mg pH 7.4
Relative Uptake (Da)
Exposure Time (minutes)
ACRA_ECOLI
AAVTAAKAAVE
155 - 165
1001.563 Da
pH 7.4
1 mM Mg pH 7.4
Relative Uptake (Da)
Exposure Time (minutes)

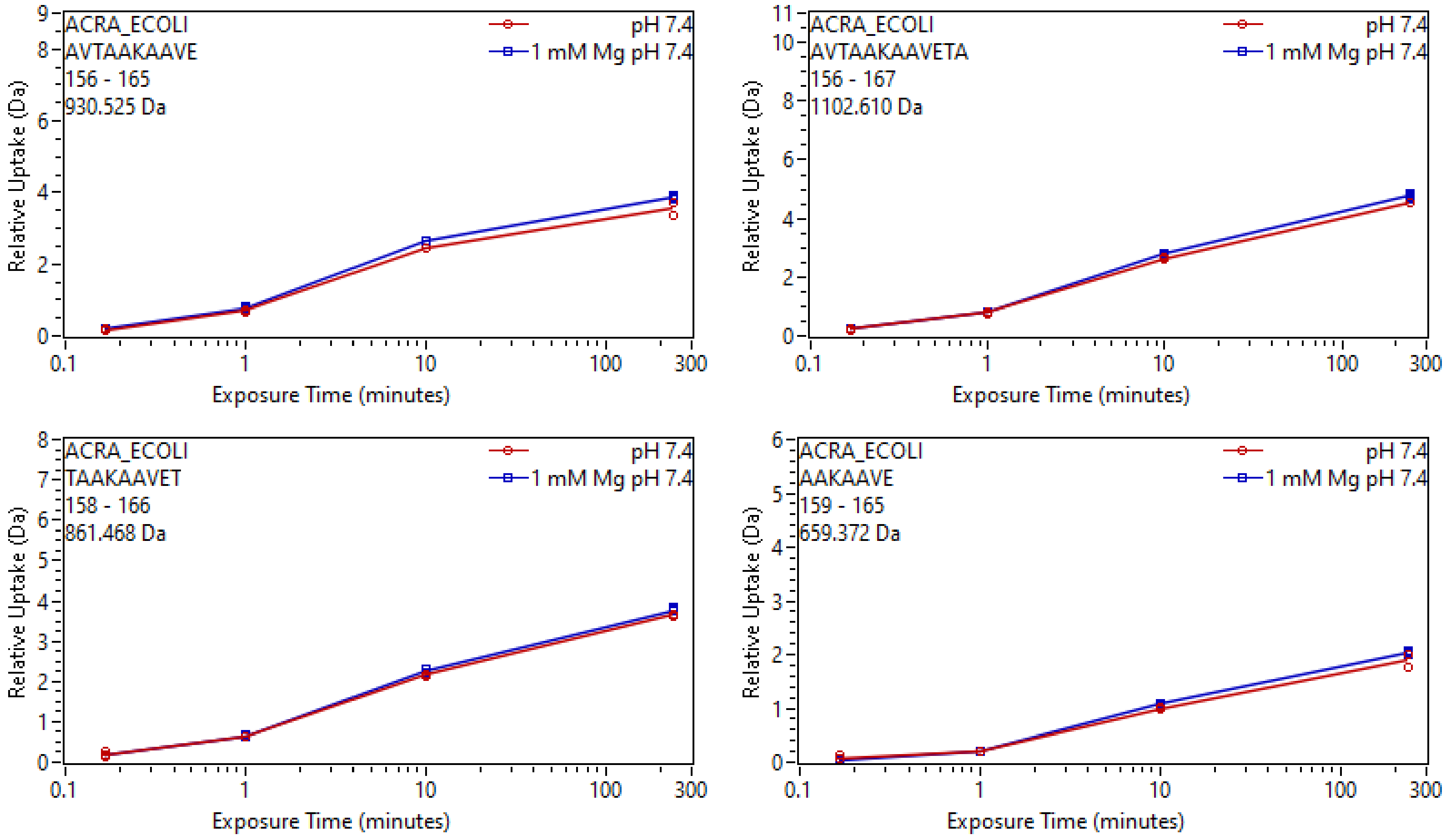
ACRA_ECOLI
AVTAAKAAVE
156 - 165
930.525 Da
pH 7.4
1 mM Mg pH 7.4
Relative Uptake (Da)
Exposure Time (minutes)
ACRA_ECOLI
AVTAAKAAVETA
156 - 167
1102.610 Da
pH 7.4
1 mM Mg pH 7.4
Relative Uptake (Da)
Exposure Time (minutes)
ACRA_ECOLI
TAAKAAVET
158 - 166
861.468 Da
pH 7.4
1 mM Mg pH 7.4
Relative Uptake (Da)
Exposure Time (minutes)
ACRA_ECOLI
AAKAAVE
159 - 165
659.372 Da
pH 7.4
1 mM Mg pH 7.4
Relative Uptake (Da)
Exposure Time (minutes)

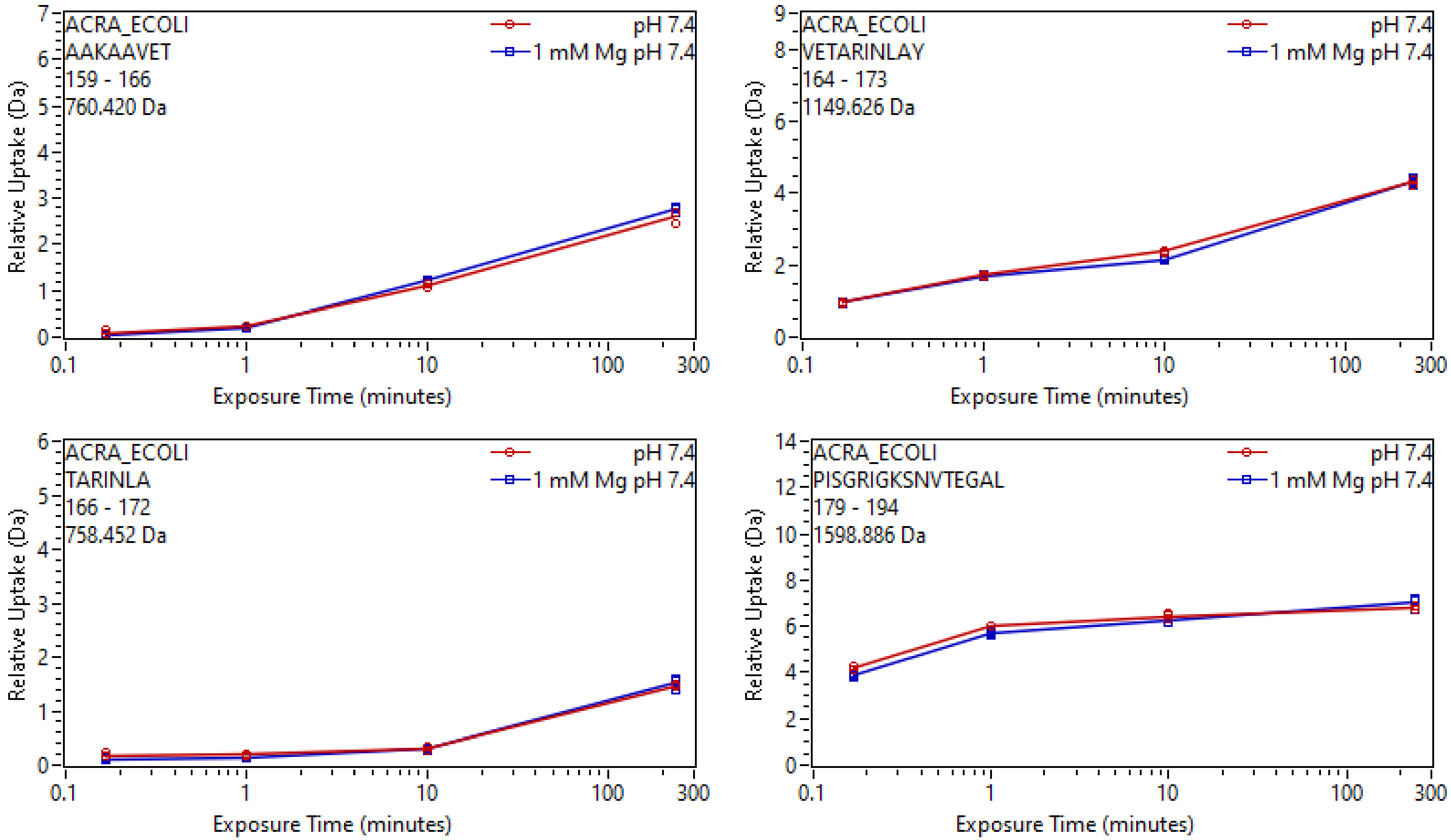

ACRA_ECOLI
AAKAAVET
159 - 166
760.420 Da
pH 7.4
1 mM Mg pH 7.4
Relative Uptake (Da)
Exposure Time (minutes)
ACRA_ECOLI
VETARINLAY
164 - 173
1149.626 Da
pH 7.4
1 mM Mg pH 7.4
Relative Uptake (Da)
Exposure Time (minutes)
ACRA_ECOLI
TARINLA
166 - 172
758.452 Da
pH 7.4
1 mM Mg pH 7.4
Relative Uptake (Da)
Exposure Time (minutes)
ACRA_ECOLI
PISGRIGKSNVTEGAL
179 - 194
1598.886 Da
pH 7.4
1 mM Mg pH 7.4
Relative Uptake (Da)
Exposure Time (minutes)

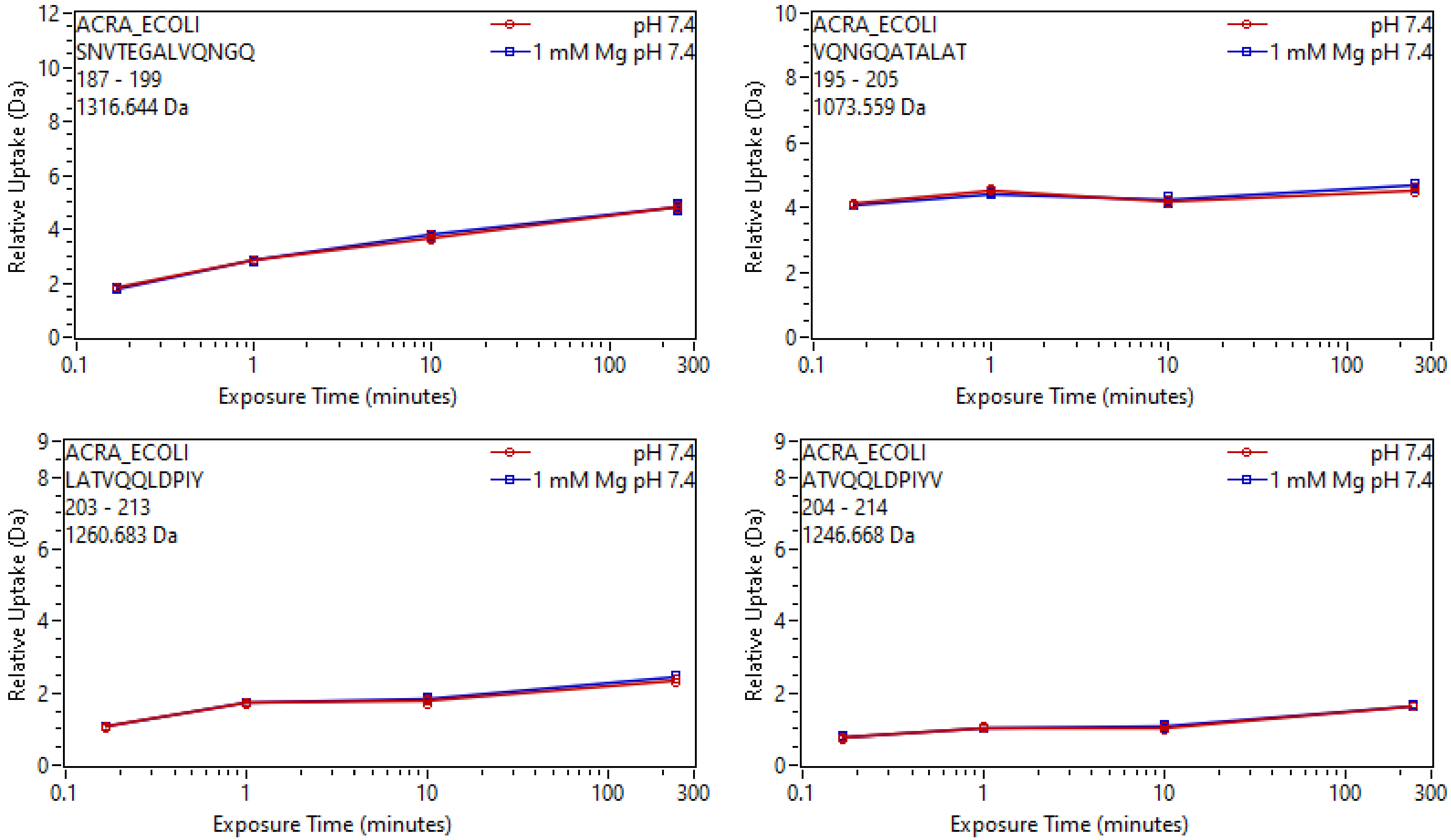

ACRA_ECOLI
SNVTEGALVQNGQ
187 - 199
1316.644 Da
pH 7.4
1 mM Mg pH 7.4
Relative Uptake (Da)
Exposure Time (minutes)
ACRA_ECOLI
VQNGQATALAT
195 - 205
1073.559 Da
pH 7.4
1 mM Mg pH 7.4
Relative Uptake (Da)
Exposure Time (minutes)
ACRA_ECOLI
LATVQQLDPIY
203 - 213
1260.683 Da
pH 7.4
1 mM Mg pH 7.4
Relative Uptake (Da)
Exposure Time (minutes)
ACRA_ECOLI
ATVQQLDPIYV
204 - 214
1246.668 Da
pH 7.4
1 mM Mg pH 7.4
Relative Uptake (Da)
Exposure Time (minutes)

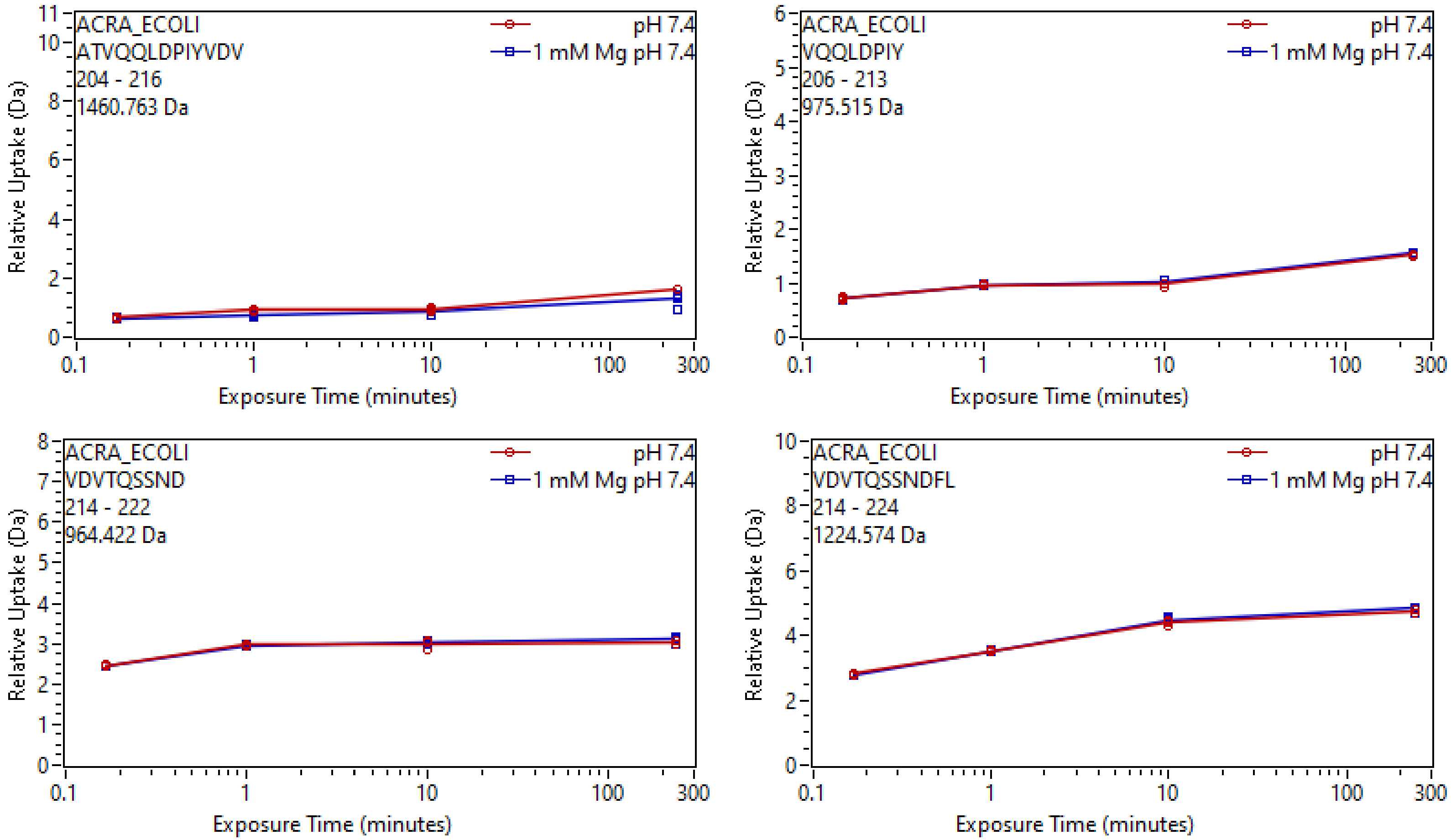
ACRA_ECOLI
ATVQQLDPIYVDV
204 - 216
1460.763 Da
pH 7.4
1 mM Mg pH 7.4
Relative Uptake (Da)
Exposure Time (minutes)
ACRA_ECOLI
VQQLDPIY
206 - 213
975.515 Da
pH 7.4
1 mM Mg pH 7.4
Relative Uptake (Da)
Exposure Time (minutes)
ACRA_ECOLI
VDVTQSSND
214 - 222
964.422 Da
pH 7.4
1 mM Mg pH 7.4
Relative Uptake (Da)
Exposure Time (minutes)
ACRA_ECOLI
VDVTQSSNDFL
214 - 224
1224.574 Da
pH 7.4
1 mM Mg pH 7.4
Relative Uptake (Da)
Exposure Time (minutes)

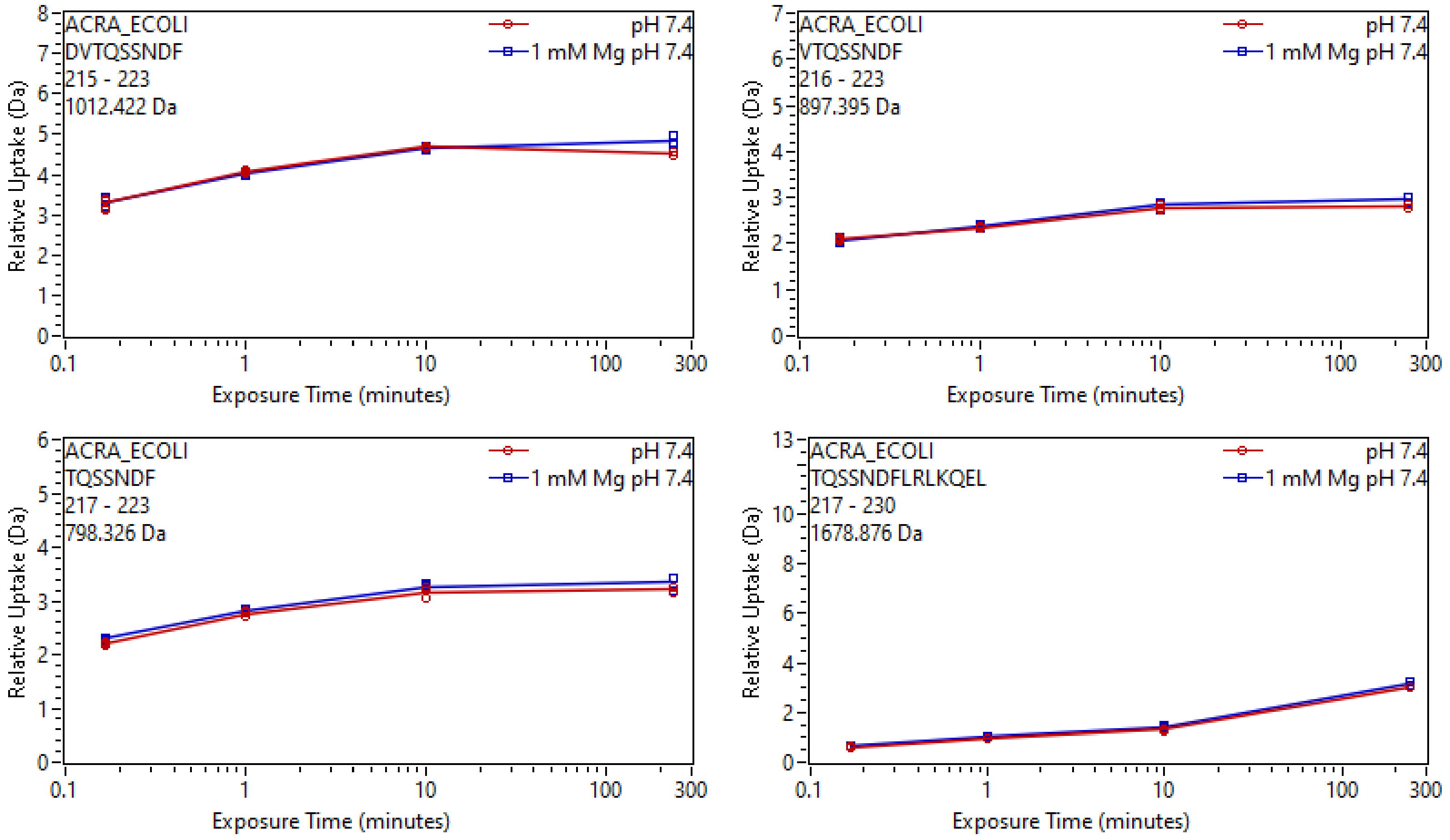

ACRA_ECOLI
DVTQSSNDF
215 - 223
1012.422 Da
pH 7.4
1 mM Mg pH 7.4
Relative Uptake (Da)
Exposure Time (minutes)
ACRA_ECOLI
VTQSSNDF
216 - 223
897.395 Da
ACRA_ECOLI
TQSSNDF
217 - 223
798.326 Da
ACRA_ECOLI
TQSSNDFLRLKQEL
217 - 230
1678.876 Da

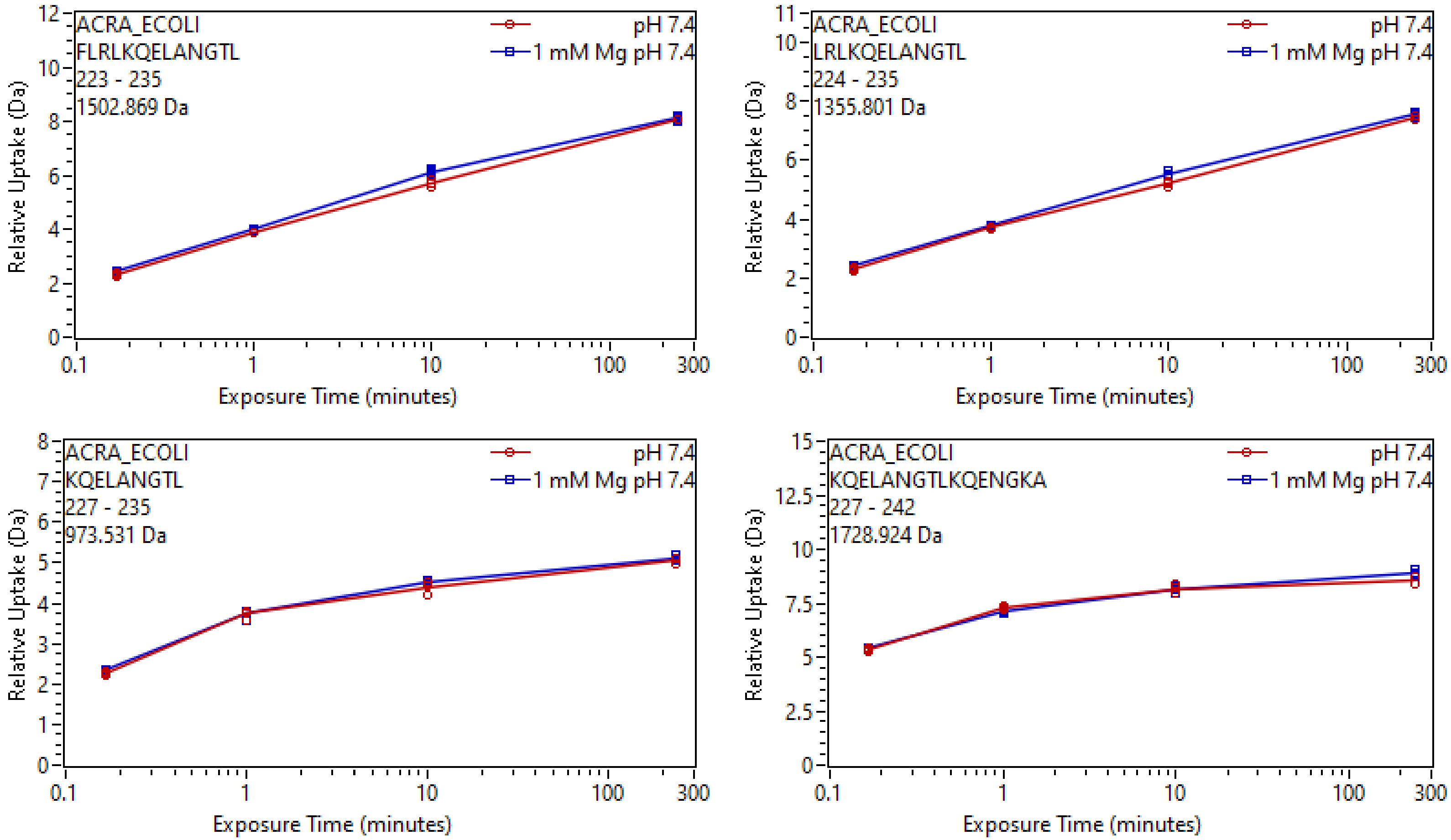
ACRA_ECOLI
FLRLKQELANGTL
223 - 235
1502.869 Da
pH 7.4
1 mM Mg pH 7.4
Relative Uptake (Da)
Exposure Time (minutes)
ACRA_ECOLI
LRLKQELANGTL
224 - 235
1355.801 Da
pH 7.4
1 mM Mg pH 7.4
Relative Uptake (Da)
Exposure Time (minutes)
ACRA_ECOLI
KQELANGTL
227 - 235
973.531 Da
pH 7.4
1 mM Mg pH 7.4
Relative Uptake (Da)
Exposure Time (minutes)
ACRA_ECOLI
KQELANGTLKQENGKA
227 - 242
1728.924 Da
pH 7.4
1 mM Mg pH 7.4
Relative Uptake (Da)
Exposure Time (minutes)

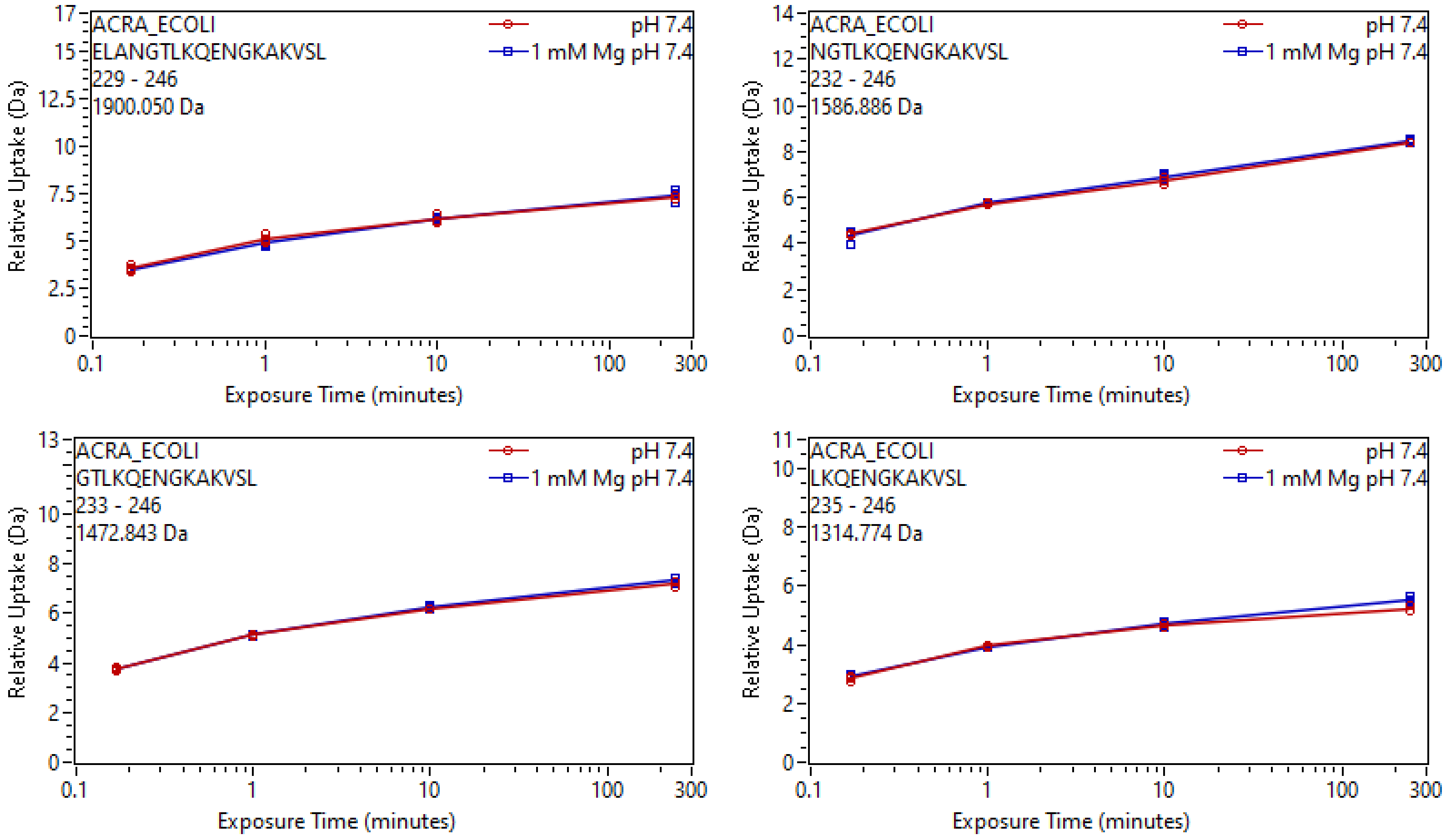

ACRA_ECOLI
ELANGTLKQENGKAKVSL
229 - 246
1900.050 Da
pH 7.4
1 mM Mg pH 7.4
Relative Uptake (Da)
Exposure Time (minutes)
ACRA_ECOLI
NGTLKQENGKAKVSL
232 - 246
1586.886 Da
ACRA_ECOLI
GTLKQENGKAKVSL
233 - 246
1472.843 Da
ACRA_ECOLI
LKQENGKAKVSL
235 - 246
1314.774 Da

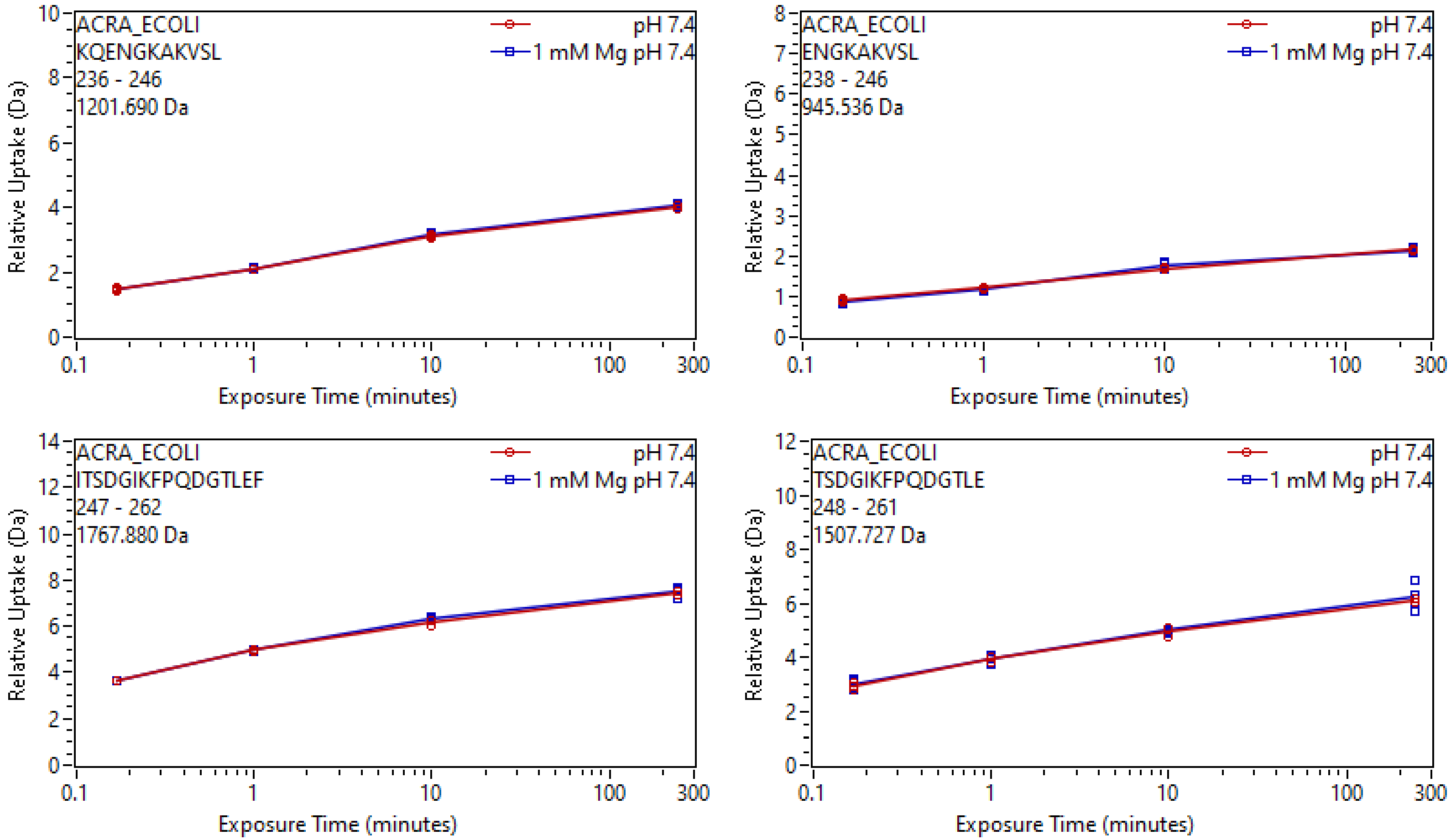

ACRA_ECOLI
KQENGKAKVSL
236 - 246
1201.690 Da
pH 7.4
1 mM Mg pH 7.4
Relative Uptake (Da)
Exposure Time (minutes)
ACRA_ECOLI
ENGKAKVSL
238 - 246
945.536 Da
ACRA_ECOLI
ITSDGIKFPQDGTLEF
247 - 262
1767.880 Da
ACRA_ECOLI
TSDGIKFPQDGTLE
248 - 261
1507.727 Da

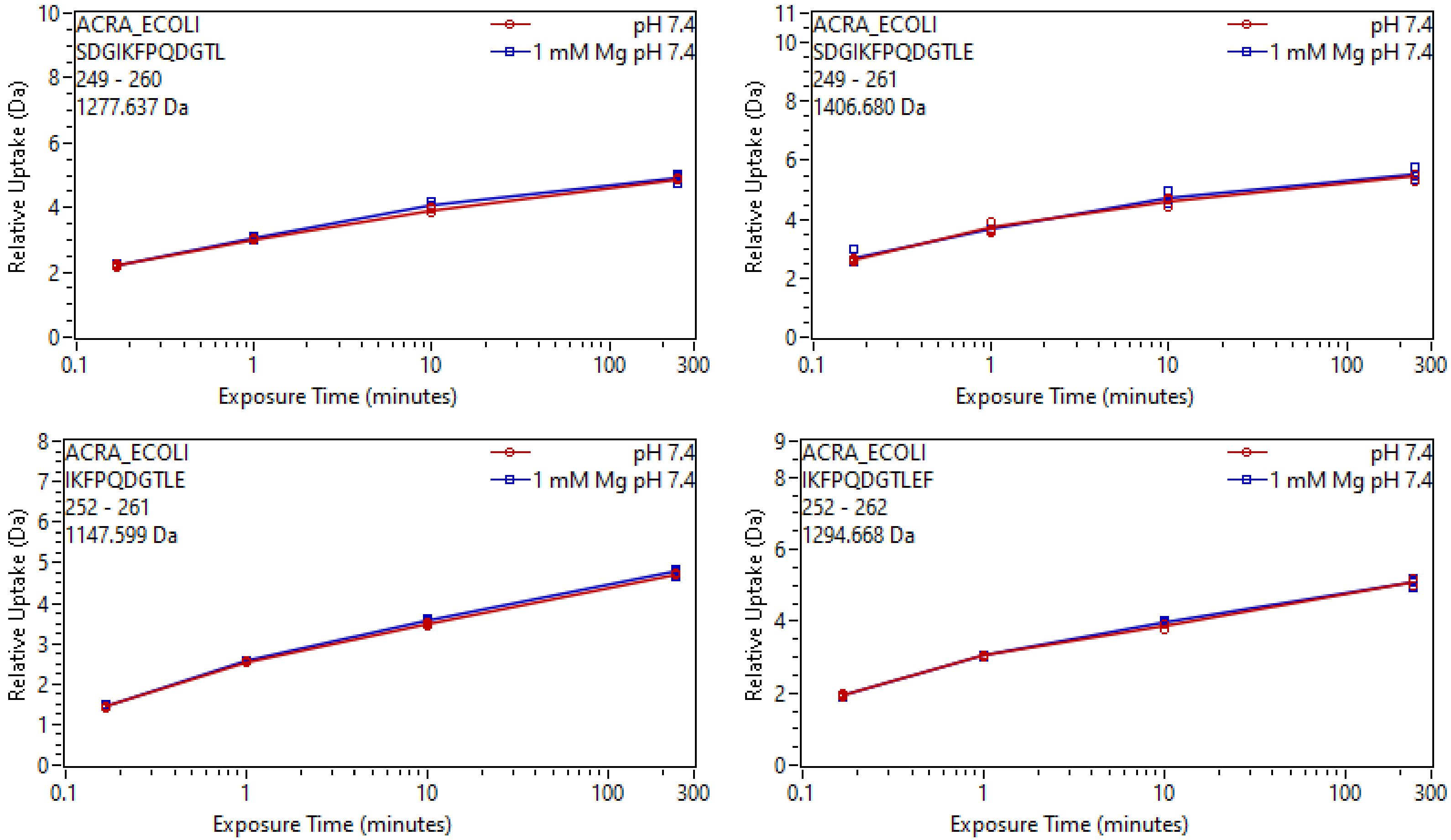
ACRA_ECOLI
SDGIKFPQDGTL
249 - 260
1277.637 Da
pH 7.4
1 mM Mg pH 7.4
Relative Uptake (Da)
Exposure Time (minutes)
ACRA_ECOLI
SDGIKFPQDGTLE
249 - 261
1406.680 Da
pH 7.4
1 mM Mg pH 7.4
Relative Uptake (Da)
Exposure Time (minutes)
ACRA_ECOLI
IKFPQDGTLE
252 - 261
1147.599 Da
pH 7.4
1 mM Mg pH 7.4
Relative Uptake (Da)
Exposure Time (minutes)
ACRA_ECOLI
IKFPQDGTLEF
252 - 262
1294.668 Da
pH 7.4
1 mM Mg pH 7.4
Relative Uptake (Da)
Exposure Time (minutes)

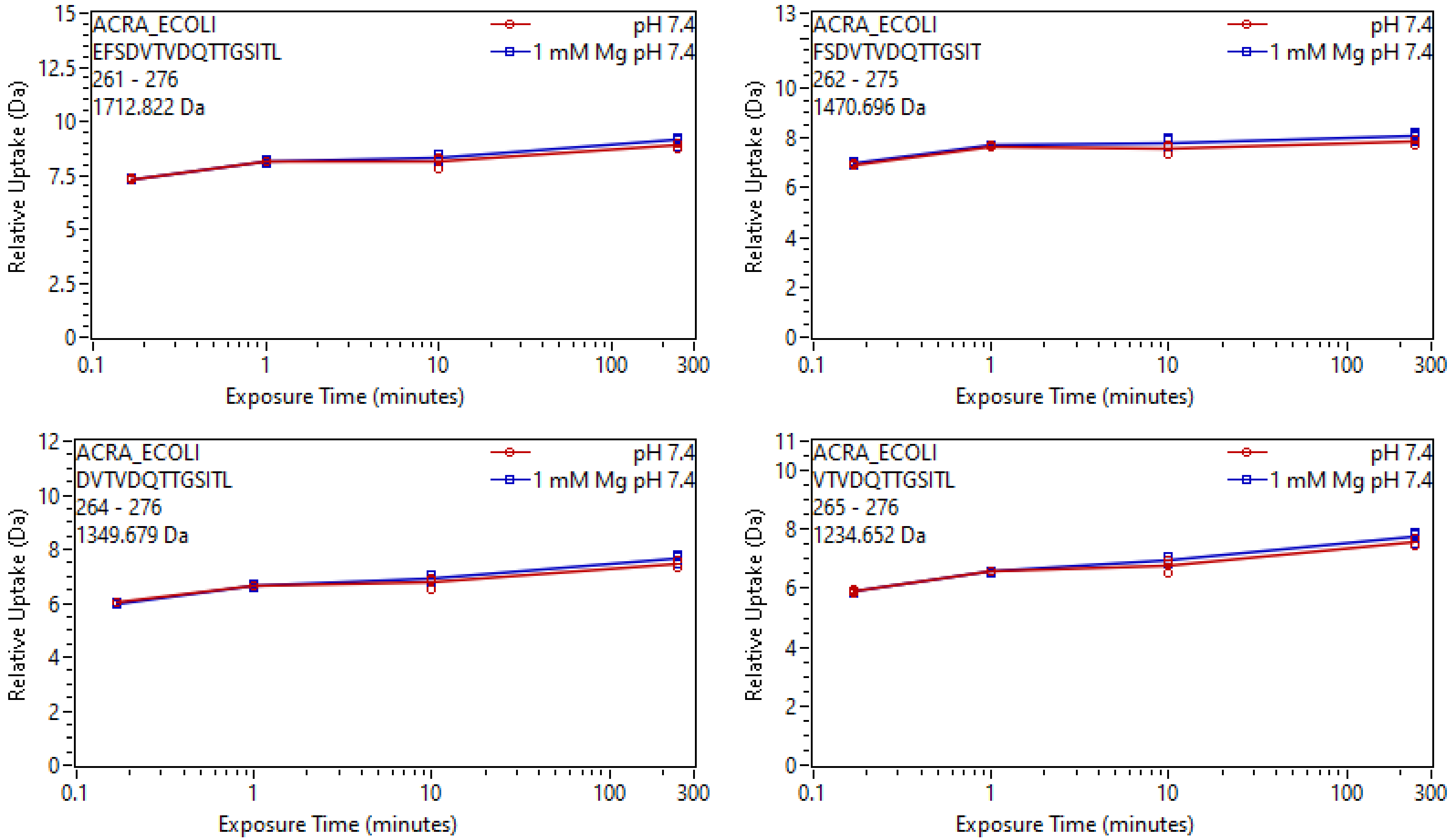

ACRA_ECOLI
EFSDVTVDQTTGSITL
261 - 276
1712.822 Da
pH 7.4
1 mM Mg pH 7.4
Relative Uptake (Da)
Exposure Time (minutes)
ACRA_ECOLI
FSDVTVDQTTGSIT
262 - 275
1470.696 Da
pH 7.4
1 mM Mg pH 7.4
Relative Uptake (Da)
Exposure Time (minutes)
ACRA_ECOLI
DVTVDQTTGSITL
264 - 276
1349.679 Da
pH 7.4
1 mM Mg pH 7.4
Relative Uptake (Da)
Exposure Time (minutes)
ACRA_ECOLI
VTVDQTTGSITL
265 - 276
1234.652 Da
pH 7.4
1 mM Mg pH 7.4
Relative Uptake (Da)
Exposure Time (minutes)

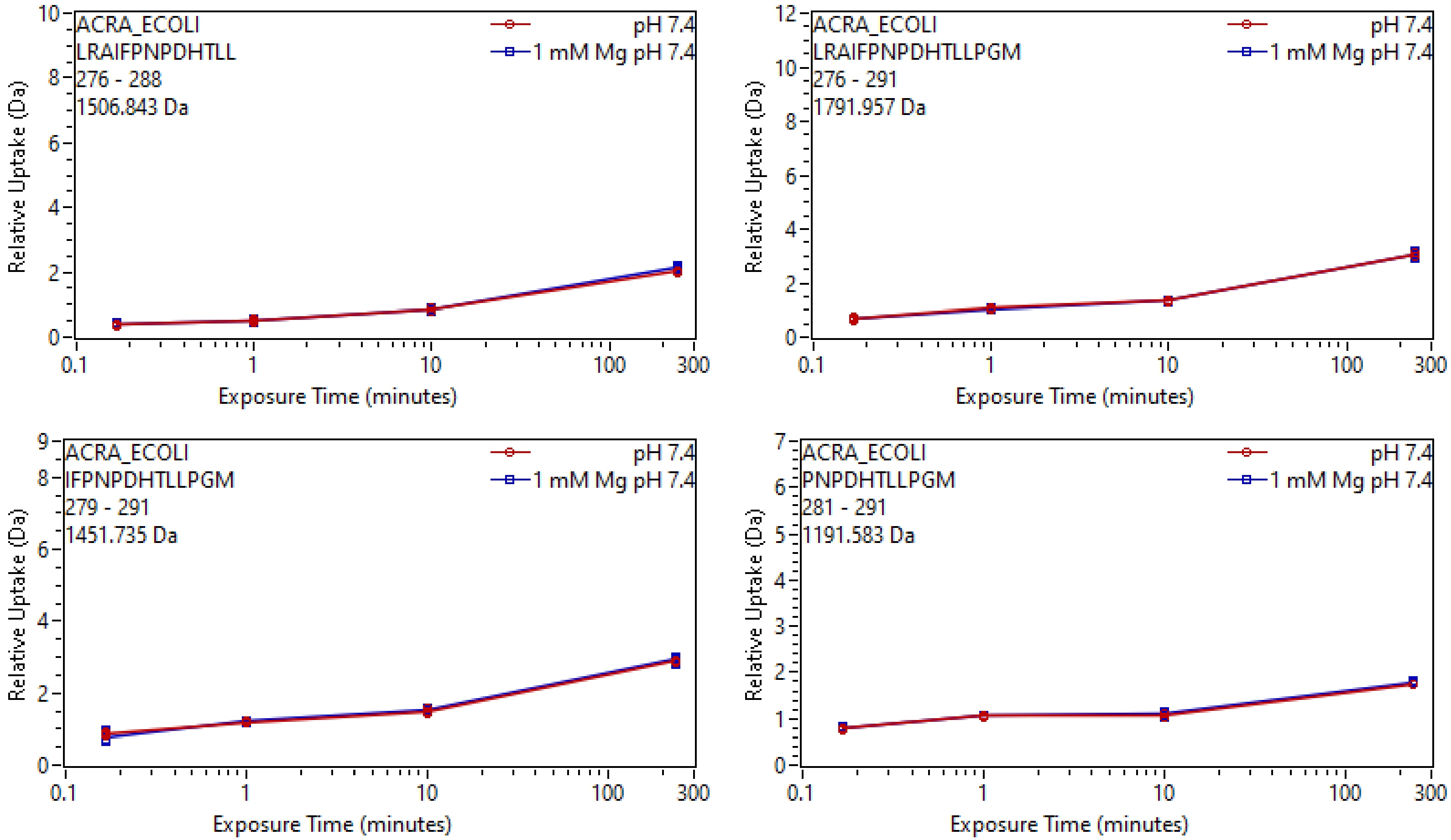

ACRA_ECOLI
LRAIFPNPDHTLL
276 - 288
1506.843 Da
pH 7.4
1 mM Mg pH 7.4
Relative Uptake (Da)
Exposure Time (minutes)
0.1
1
10
100
300
ACRA_ECOLI
LRAIFPNPDHTLLPGM
276 - 291
1791.957 Da
pH 7.4
1 mM Mg pH 7.4
Relative Uptake (Da)
Exposure Time (minutes)
ACRA_ECOLI
IFPNPDHTLLPGM
279 - 291
1451.735 Da
pH 7.4
1 mM Mg pH 7.4
Relative Uptake (Da)
Exposure Time (minutes)
ACRA_ECOLI
PNPDHTLLPGM
281 - 291
1191.583 Da
pH 7.4
1 mM Mg pH 7.4
Relative Uptake (Da)
Exposure Time (minutes)

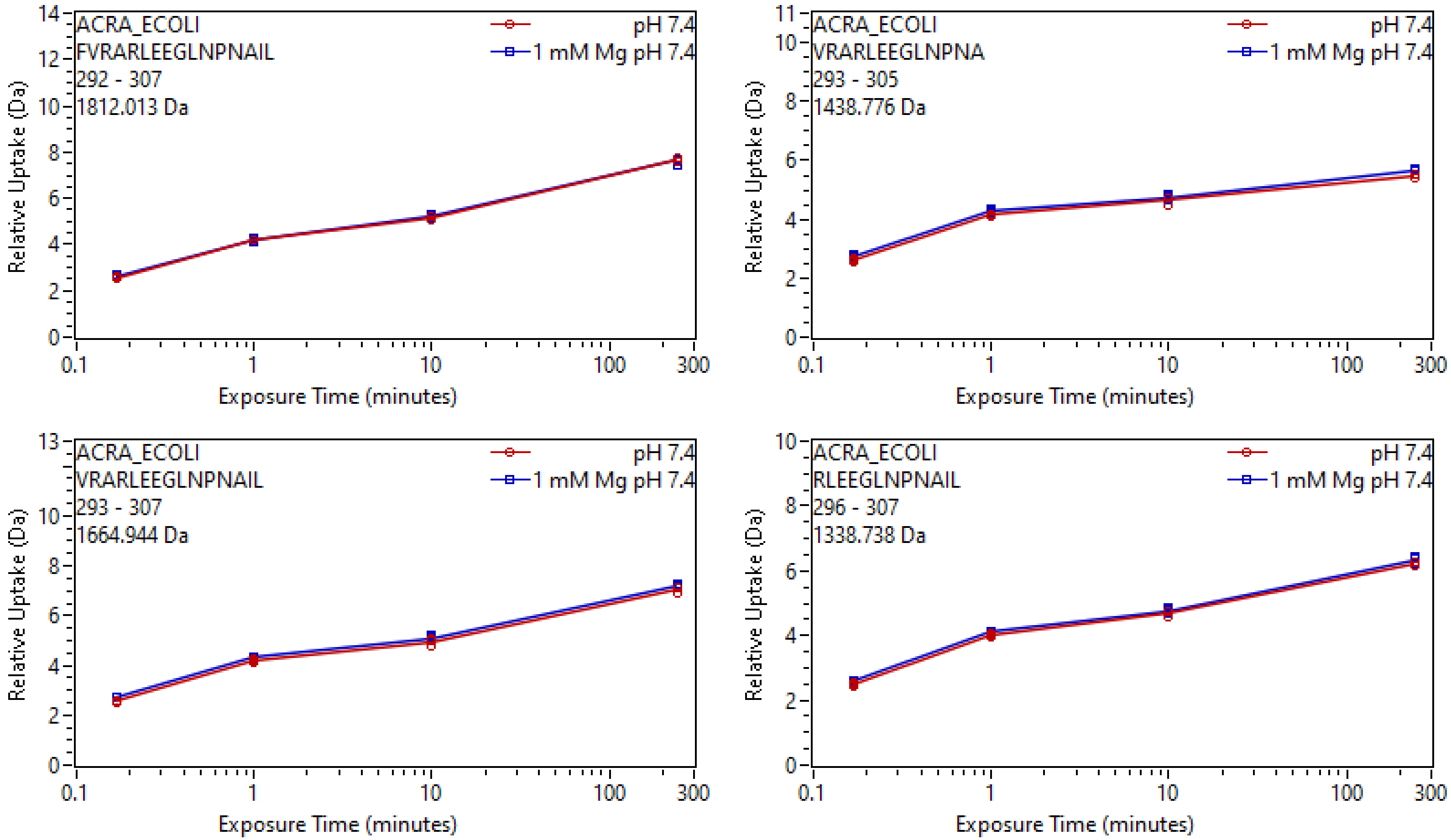
ACRA_ECOLI
FVRARLEEGLNPNAIL
292 - 307
1812.013 Da
ACRA_ECOLI
VRARLEEGLNPNA
293 - 305
1438.776 Da
ACRA_ECOLI
VRARLEEGLNPNAIL
293 - 307
1664.944 Da
ACRA_ECOLI
RLEEGLNPNAIL
296 - 307
1338.738 Da
pH 7.4
1 mM Mg pH 7.4
Relative Uptake (Da)
Exposure Time (minutes)
0.1
1
10
100
300

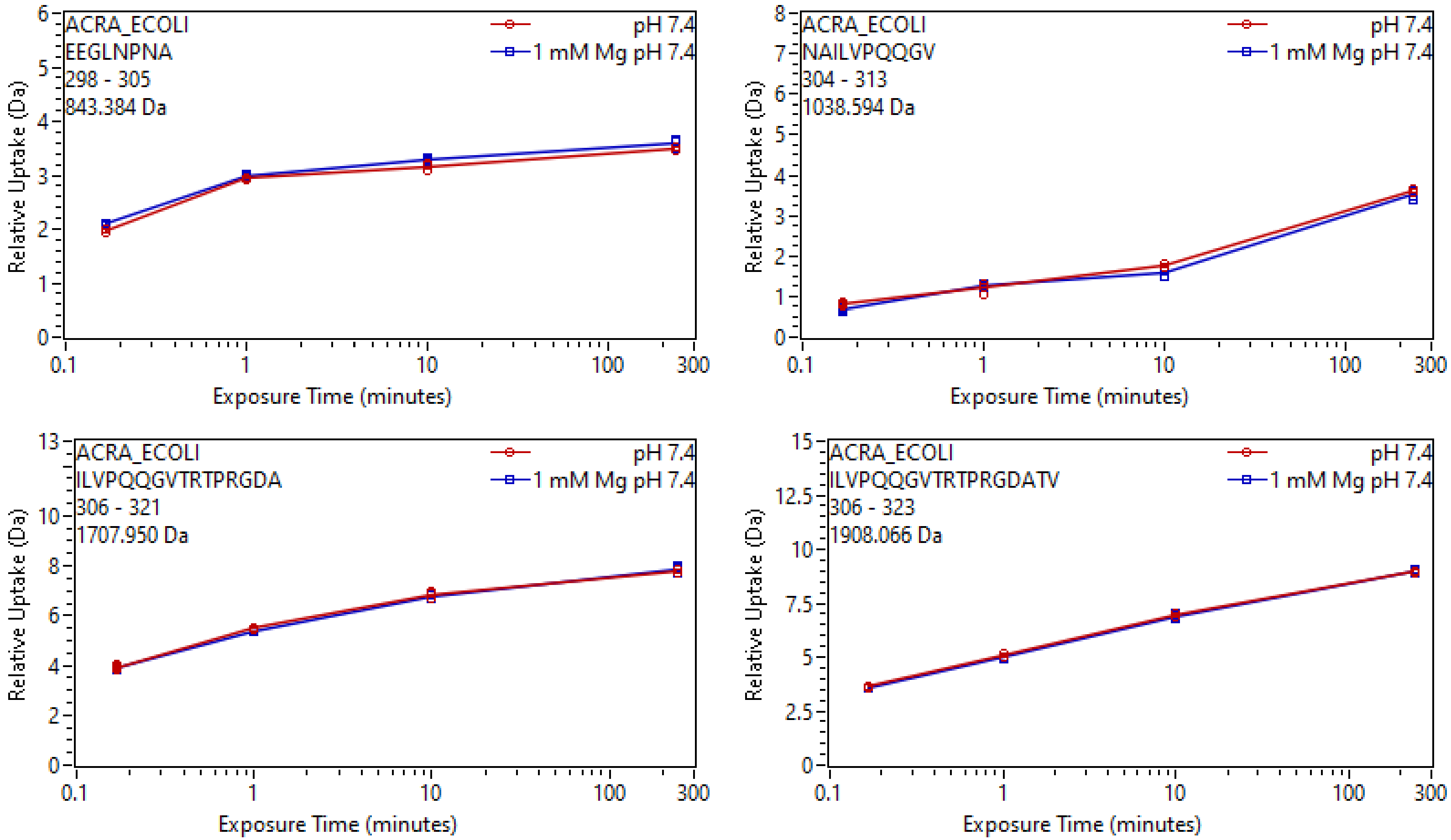
ACRA_ECOLI
EEGLNPNA
298 - 305
843.384 Da
pH 7.4
1 mM Mg pH 7.4
Relative Uptake (Da)
Exposure Time (minutes)
ACRA_ECOLI
NAILVPQQGV
304 - 313
1038.594 Da
ACRA_ECOLI
ILVPQQGVTRTPRGDA
306 - 321
1707.950 Da
ACRA_ECOLI
ILVPQQGVTRTPRGDATV
306 - 323
1908.066 Da

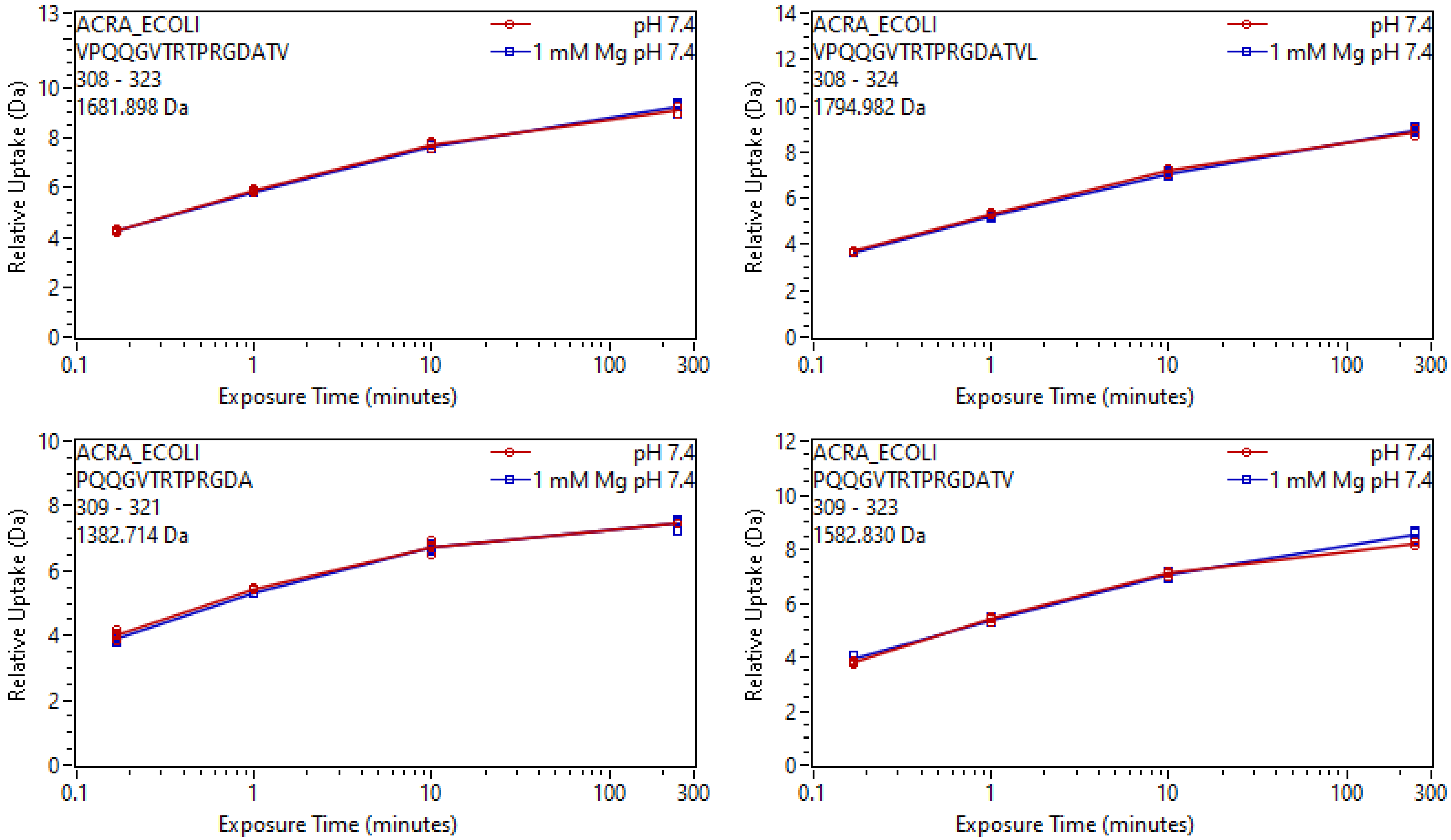
ACRA_ECOLI
VPQQGVTRTPRGDATV
308 - 323
1681.898 Da
pH 7.4
1 mM Mg pH 7.4
Relative Uptake (Da)
Exposure Time (minutes)
ACRA_ECOLI
VPQQGVTRTPRGDATVL
308 - 324
1794.982 Da
ACRA_ECOLI
PQQGVTRTPRGDA
309 - 321
1382.714 Da
ACRA_ECOLI
PQQGVTRTPRGDATV
309 - 323
1582.830 Da

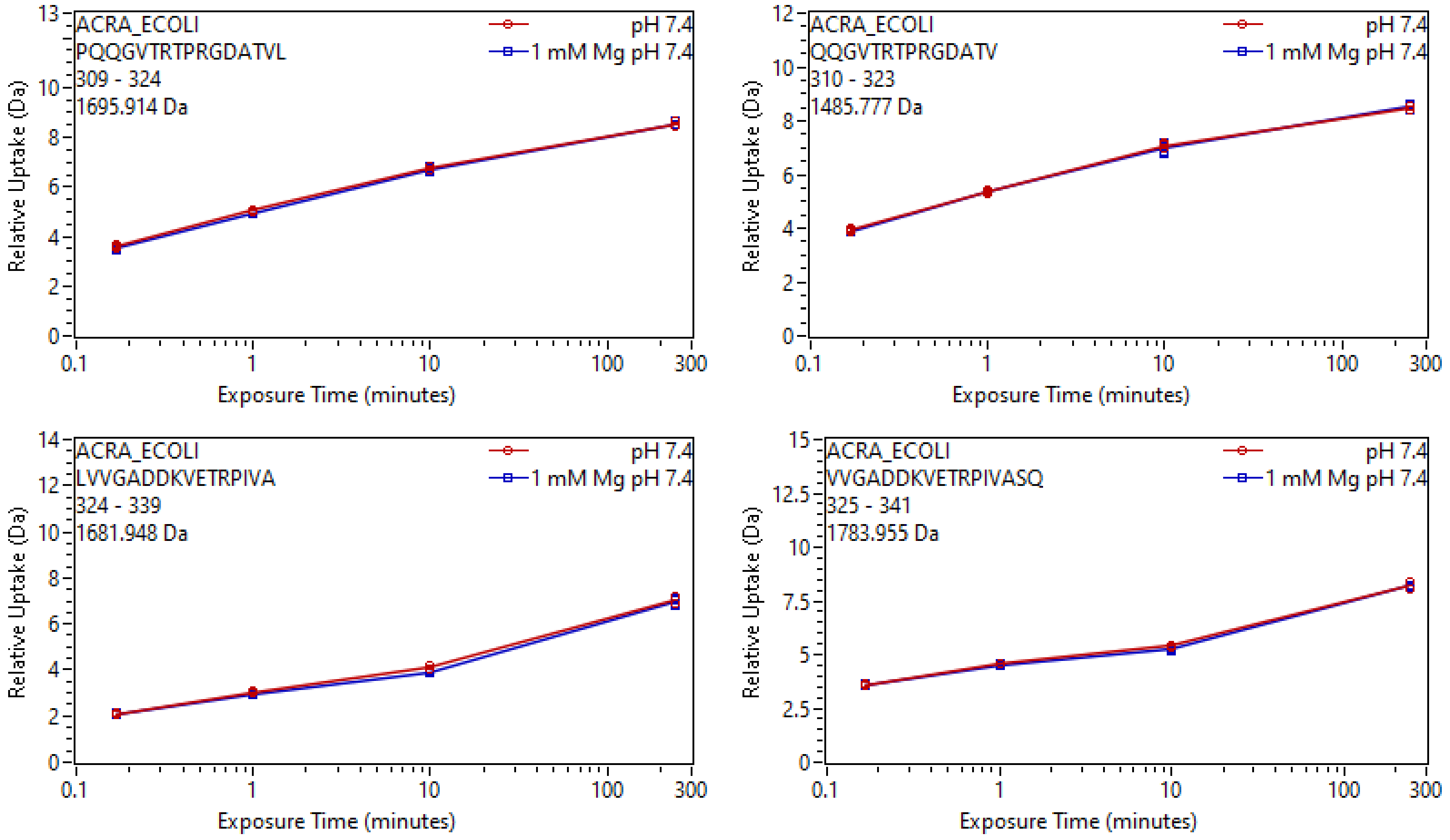

ACRA_ECOLI
PQQGVTRTPRGDATVL
309 - 324
1695.914 Da
ACRA_ECOLI
QQGVTRTPRGDATV
310 - 323
1485.777 Da
ACRA_ECOLI
LVVGADDKVETRPIVA
324 - 339
1681.948 Da
ACRA_ECOLI
VVGADDKVETRPIVASQ
325 - 341
1783.955 Da
pH 7.4
1 mM Mg pH 7.4
Relative Uptake (Da)
Exposure Time (minutes)

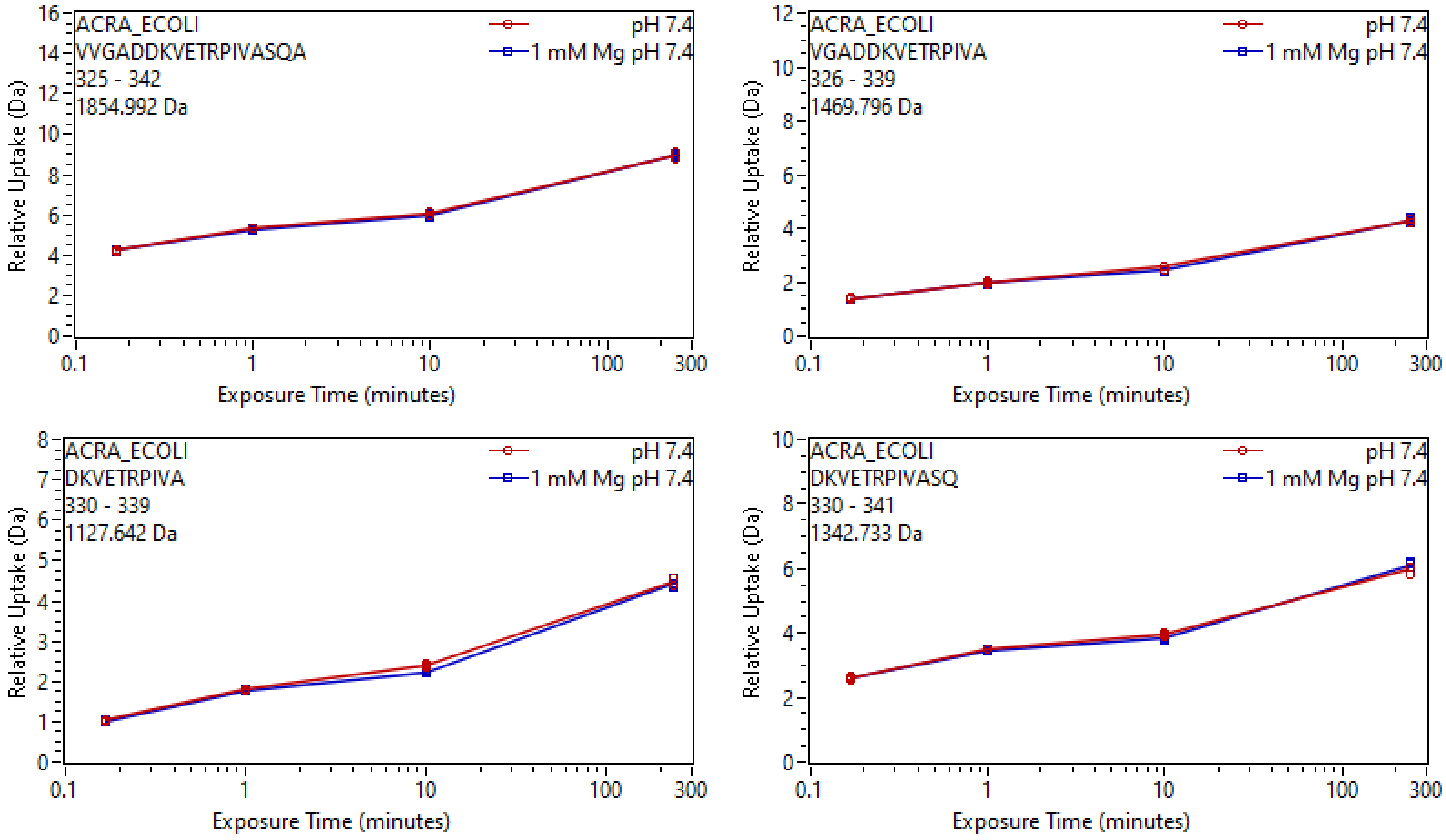
ACRA_ECOLI
VVGADDKVETRPIVASQA
325 - 342
1854.992 Da
ACRA_ECOLI
VGADDKVETRPIVA
326 - 339
1469.796 Da
ACRA_ECOLI
DKVETRPIVA
330 - 339
1127.642 Da
ACRA_ECOLI
DKVETRPIVASQ
330 - 341
1342.733 Da
pH 7.4
1 mM Mg pH 7.4
Relative Uptake (Da)
Exposure Time (minutes)

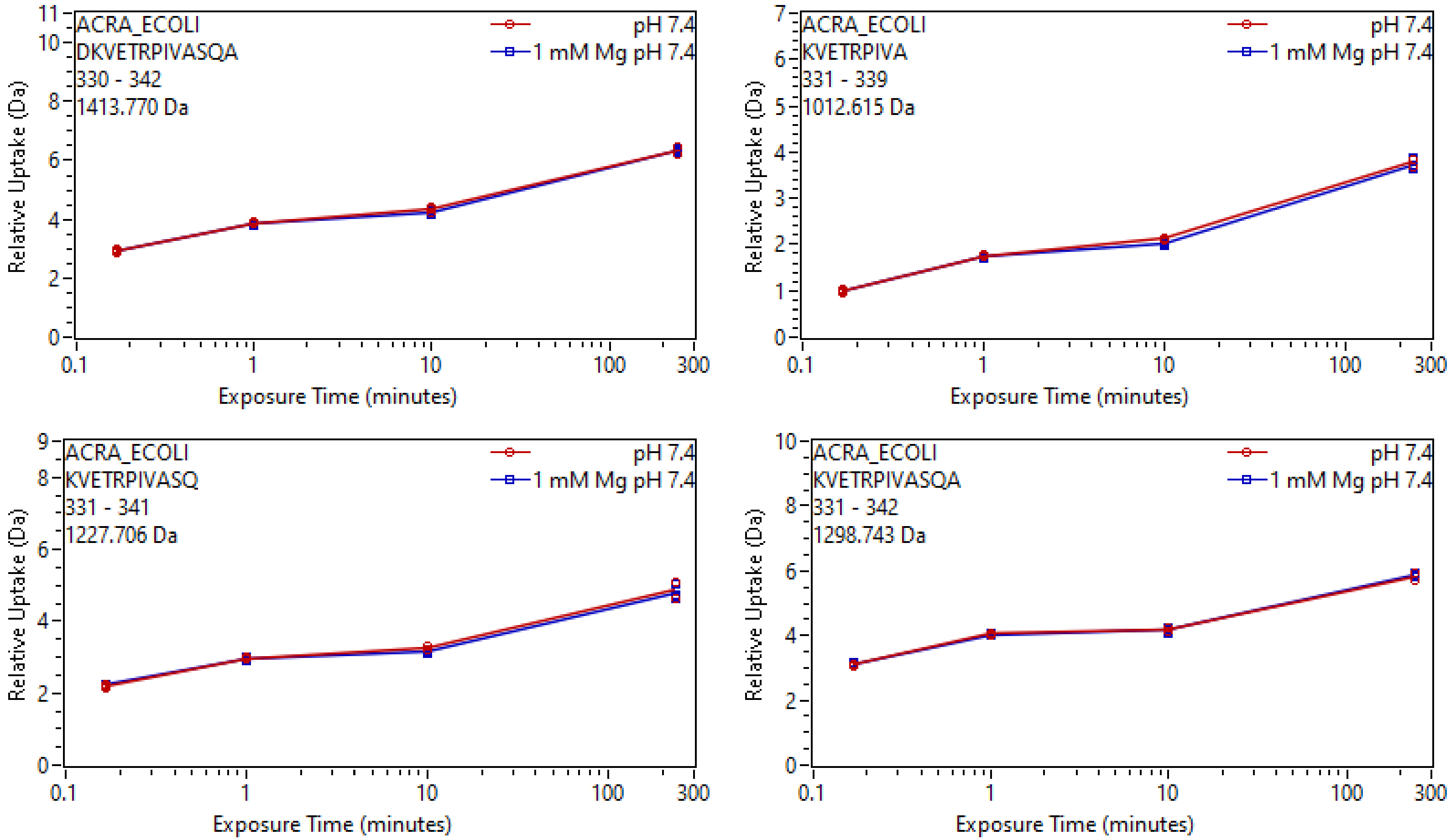
ACRA_ECOLI
DKVETRPIVASQA
330 - 342
1413.770 Da
pH 7.4
1 mM Mg pH 7.4
Relative Uptake (Da)
Exposure Time (minutes)
0
2
4
6
8
10
11
0.1
1
10
100
300
ACRA_ECOLI
KVETRPIVA
331 - 339
1012.615 Da
pH 7.4
1 mM Mg pH 7.4
Relative Uptake (Da)
Exposure Time (minutes)
0
1
2
3
4
5
6
7
0.1
1
10
100
300
ACRA_ECOLI
KVETRPIVASQ
331 - 341
1227.706 Da
pH 7.4
1 mM Mg pH 7.4
Relative Uptake (Da)
Exposure Time (minutes)
0
2
4
6
8
9
0.1
1
10
100
300
ACRA_ECOLI
KVETRPIVASQA
331 - 342
1298.743 Da
pH 7.4
1 mM Mg pH 7.4
Relative Uptake (Da)
Exposure Time (minutes)
0
2
4
6
8
10
0.1
1
10
100
300

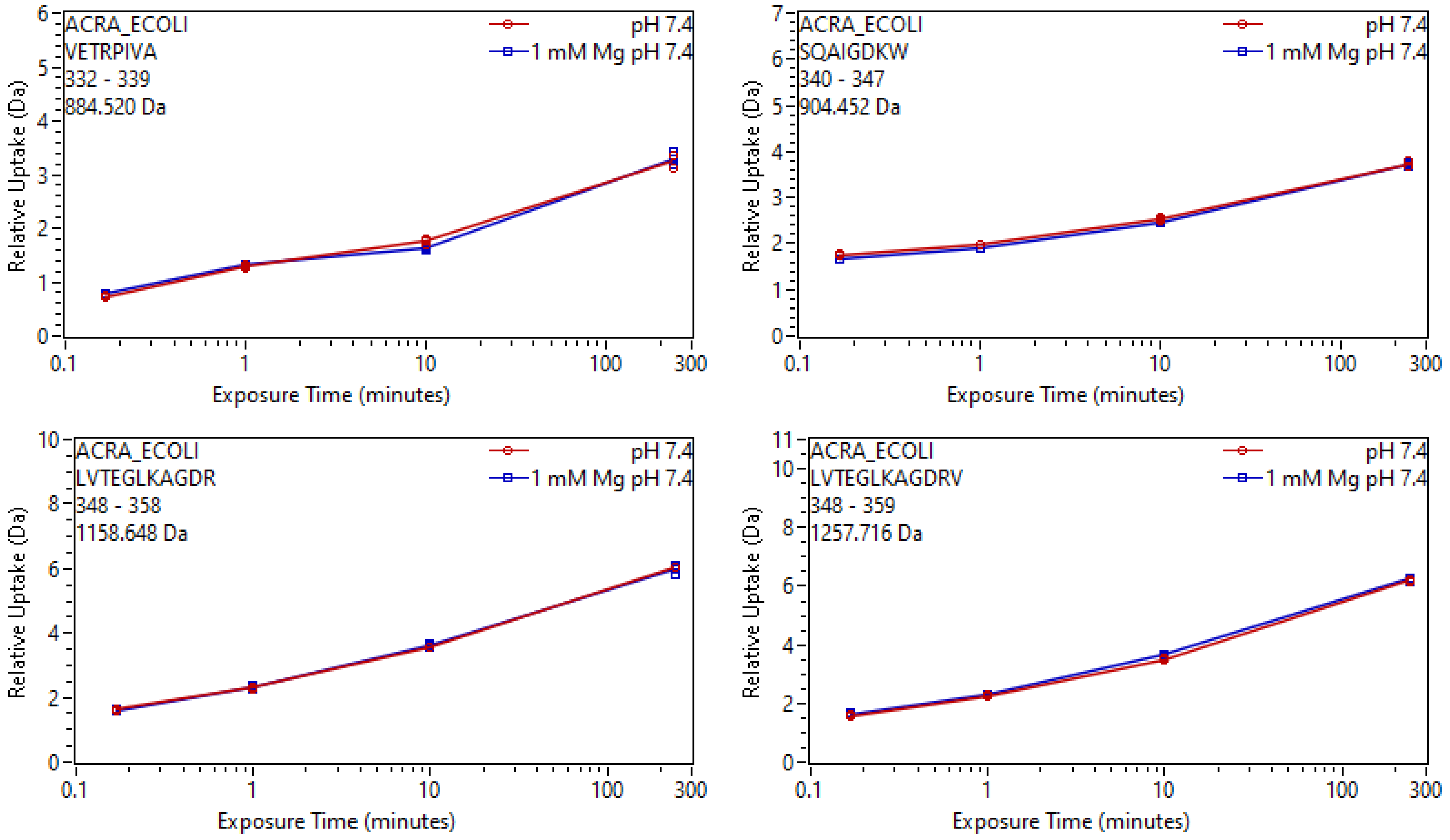
ACRA_ECOLI
VETRPIVA
332 - 339
884.520 Da
pH 7.4
1 mM Mg pH 7.4
Relative Uptake (Da)
Exposure Time (minutes)
ACRA_ECOLI
SQAIGDKW
340 - 347
904.452 Da
pH 7.4
1 mM Mg pH 7.4
Relative Uptake (Da)
Exposure Time (minutes)
ACRA_ECOLI
LVTEGLKAGDR
348 - 358
1158.648 Da
pH 7.4
1 mM Mg pH 7.4
Relative Uptake (Da)
Exposure Time (minutes)
ACRA_ECOLI
LVTEGLKAGDRV
348 - 359
1257.716 Da
pH 7.4
1 mM Mg pH 7.4
Relative Uptake (Da)
Exposure Time (minutes)

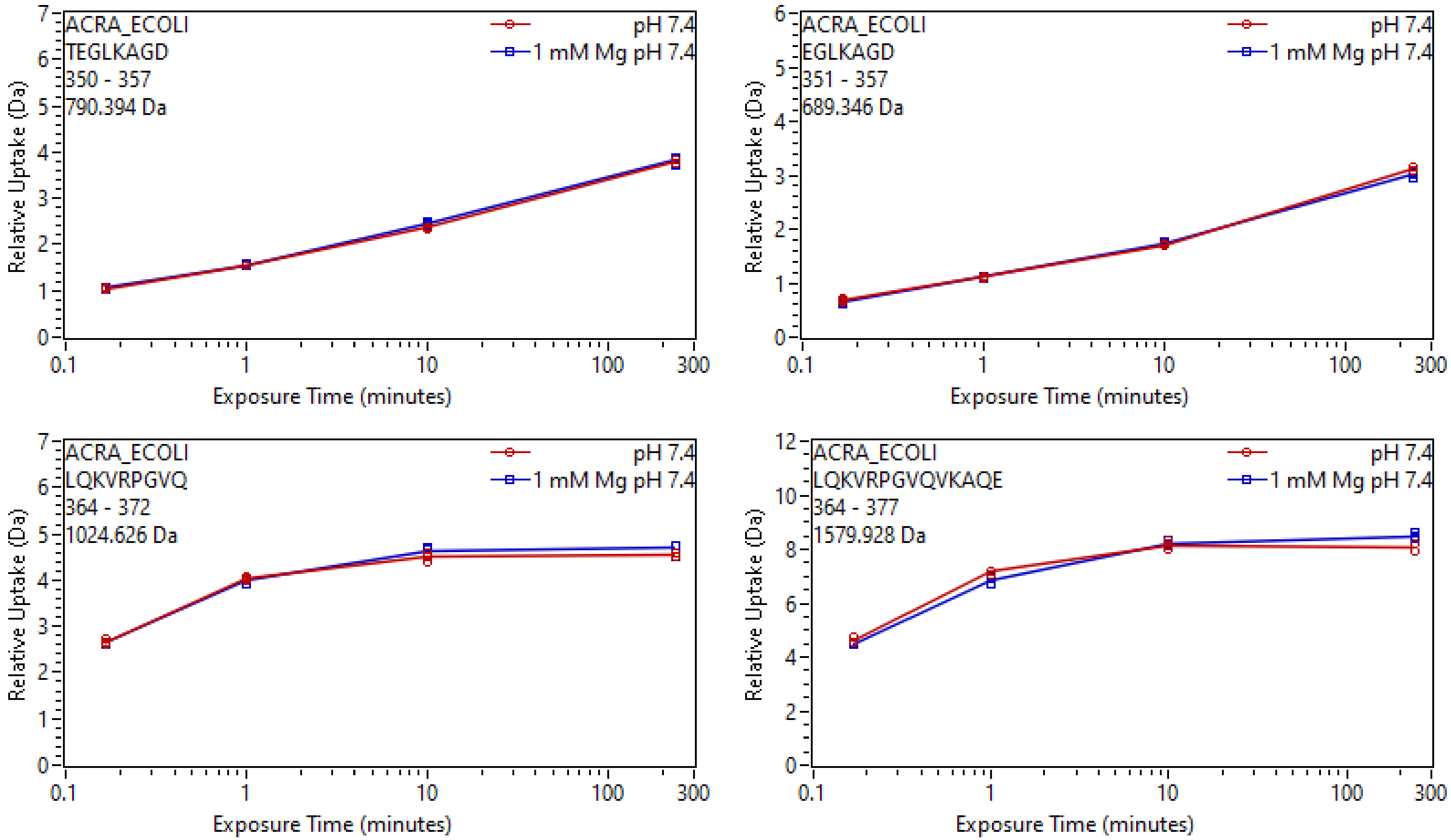
ACRA_ECOLI
TEGLKAGD
350 - 357
790.394 Da
pH 7.4
1 mM Mg pH 7.4
Relative Uptake (Da)
Exposure Time (minutes)
ACRA_ECOLI
EGLKAGD
351 - 357
689.346 Da
pH 7.4
1 mM Mg pH 7.4
Relative Uptake (Da)
Exposure Time (minutes)
ACRA_ECOLI
LQKVRPGVQ
364 - 372
1024.626 Da
pH 7.4
1 mM Mg pH 7.4
Relative Uptake (Da)
Exposure Time (minutes)
ACRA_ECOLI
LQKVRPGVQVKAQE
364 - 377
1579.928 Da
pH 7.4
1 mM Mg pH 7.4
Relative Uptake (Da)
Exposure Time (minutes)

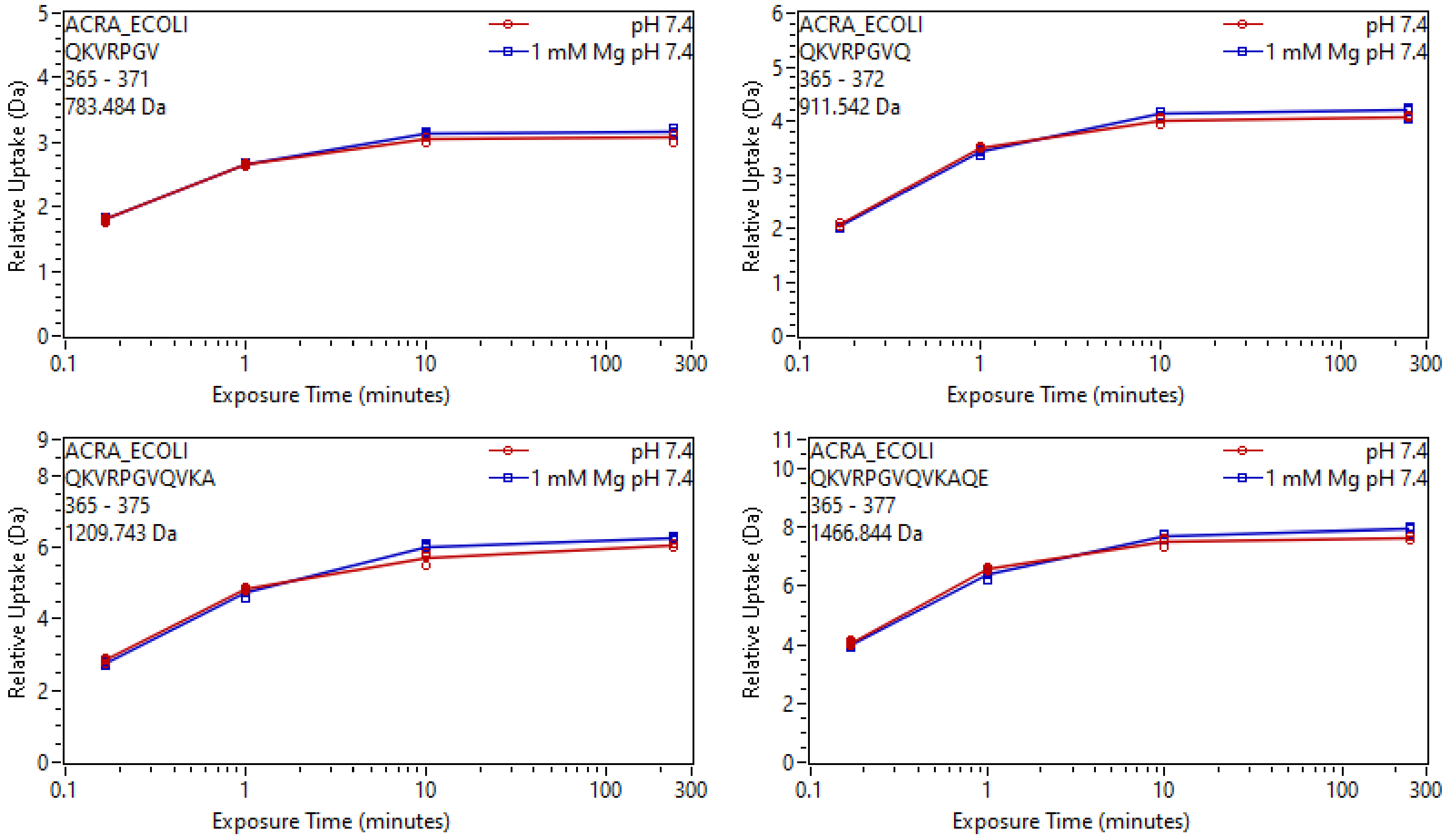
ACRA_ECOLI
QKVRPGV
365 - 371
783.484 Da
pH 7.4
1 mM Mg pH 7.4
Relative Uptake (Da)
Exposure Time (minutes)
ACRA_ECOLI
QKVRPGVQ
365 - 372
911.542 Da
pH 7.4
1 mM Mg pH 7.4
Relative Uptake (Da)
Exposure Time (minutes)
ACRA_ECOLI
QKVRPGVQVKA
365 - 375
1209.743 Da
pH 7.4
1 mM Mg pH 7.4
Relative Uptake (Da)
Exposure Time (minutes)
ACRA_ECOLI
QKVRPGVQVKAQE
365 - 377
1466.844 Da
pH 7.4
1 mM Mg pH 7.4
Relative Uptake (Da)
Exposure Time (minutes)

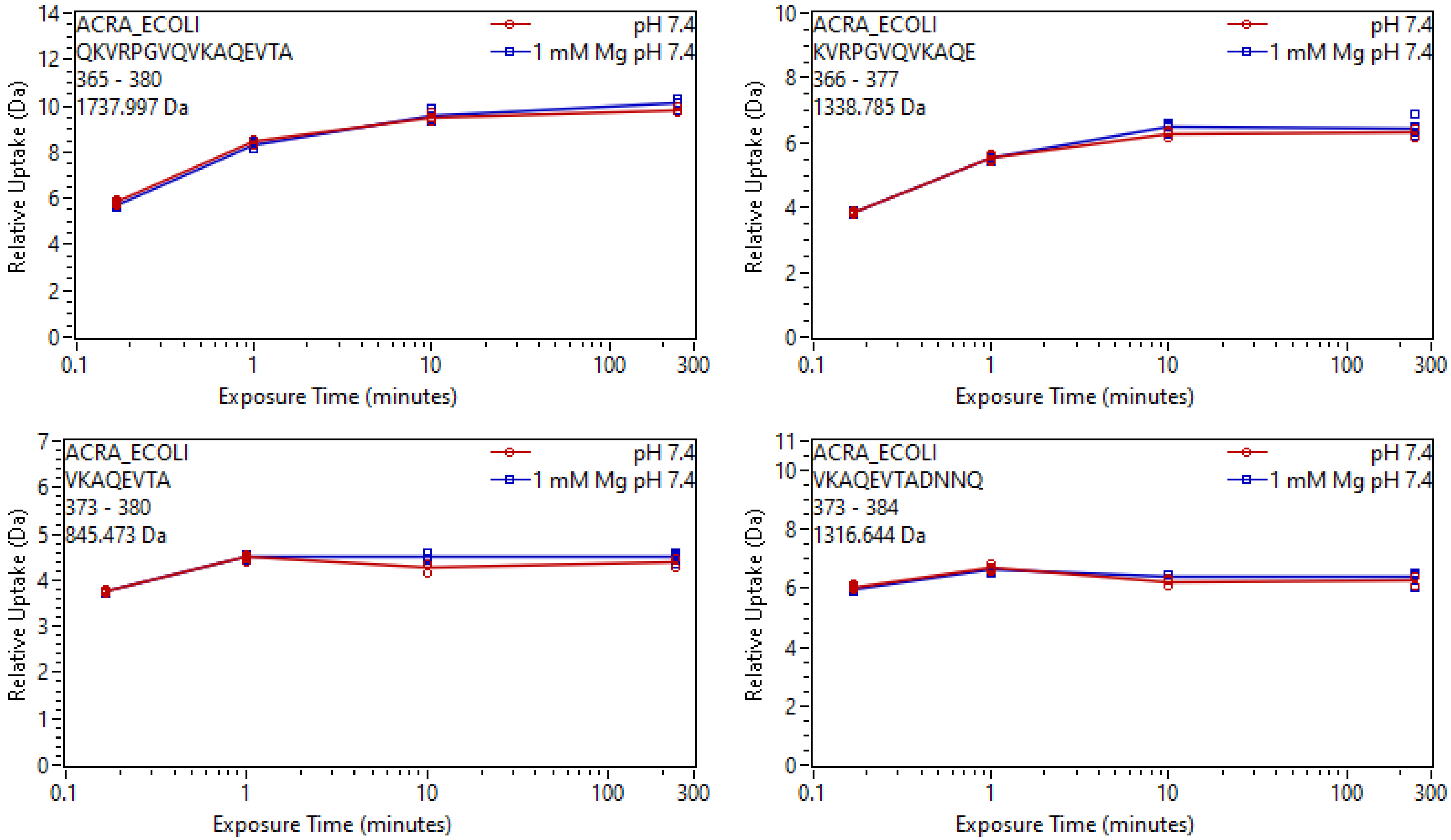
ACRA_ECOLI
QKVRPGVQVKAQEVTA
365 - 380
1737.997 Da
pH 7.4
1 mM Mg pH 7.4
Relative Uptake (Da)
Exposure Time (minutes)
ACRA_ECOLI
KVRPGVQVKAQE
366 - 377
1338.785 Da
pH 7.4
1 mM Mg pH 7.4
Relative Uptake (Da)
Exposure Time (minutes)
ACRA_ECOLI
VKAQEVTA
373 - 380
845.473 Da
pH 7.4
1 mM Mg pH 7.4
Relative Uptake (Da)
Exposure Time (minutes)
ACRA_ECOLI
VKAQEVTADNNQ
373 - 384
1316.644 Da
pH 7.4
1 mM Mg pH 7.4
Relative Uptake (Da)
Exposure Time (minutes)

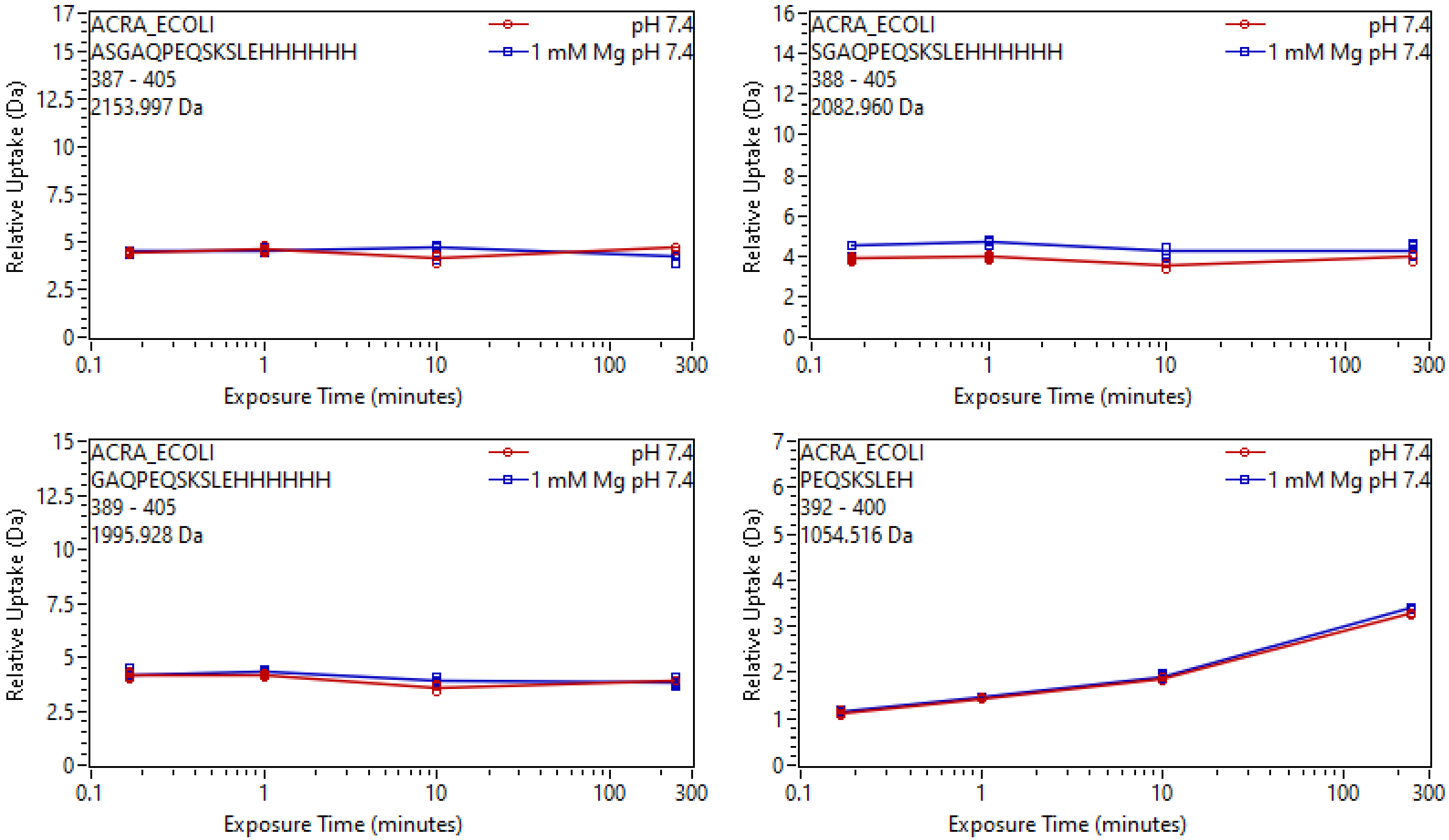
ACRA_ECOLI
ASGAQPEQSKSLEHHHHHH
387 - 405
2153.997 Da
pH 7.4
1 mM Mg pH 7.4
Relative Uptake (Da)
Exposure Time (minutes)
ACRA_ECOLI
SGAQPEQSKSLEHHHHHH
388 - 405
2082.960 Da
pH 7.4
1 mM Mg pH 7.4
Relative Uptake (Da)
Exposure Time (minutes)
ACRA_ECOLI
GAQPEQSKSLEHHHHHH
389 - 405
1995.928 Da
pH 7.4
1 mM Mg pH 7.4
Relative Uptake (Da)
Exposure Time (minutes)
ACRA_ECOLI
PEQSKSLEH
392 - 400
1054.516 Da
pH 7.4
1 mM Mg pH 7.4
Relative Uptake (Da)
Exposure Time (minutes)

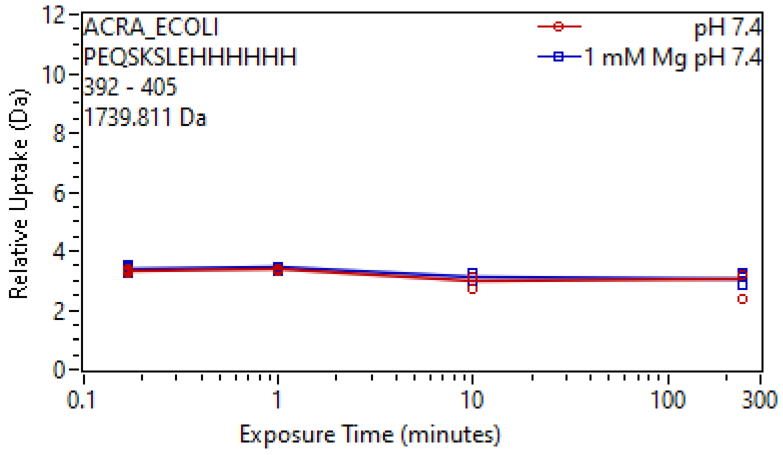
ACRA_ECOLI
PEQSKSLEHHHHHH
392 - 405
1739.811 Da
pH 7.4
1 mM Mg pH 7.4
Relative Uptake (Da)
Exposure Time (minutes)
0
2
4
6
8
10
12
0.1
1
10
100
300
