## Supplementary Data 3 for "Mg^2+^-dependent mechanism of environmental versatility in a multidrug efflux pump": Uptake plots 5.pdf

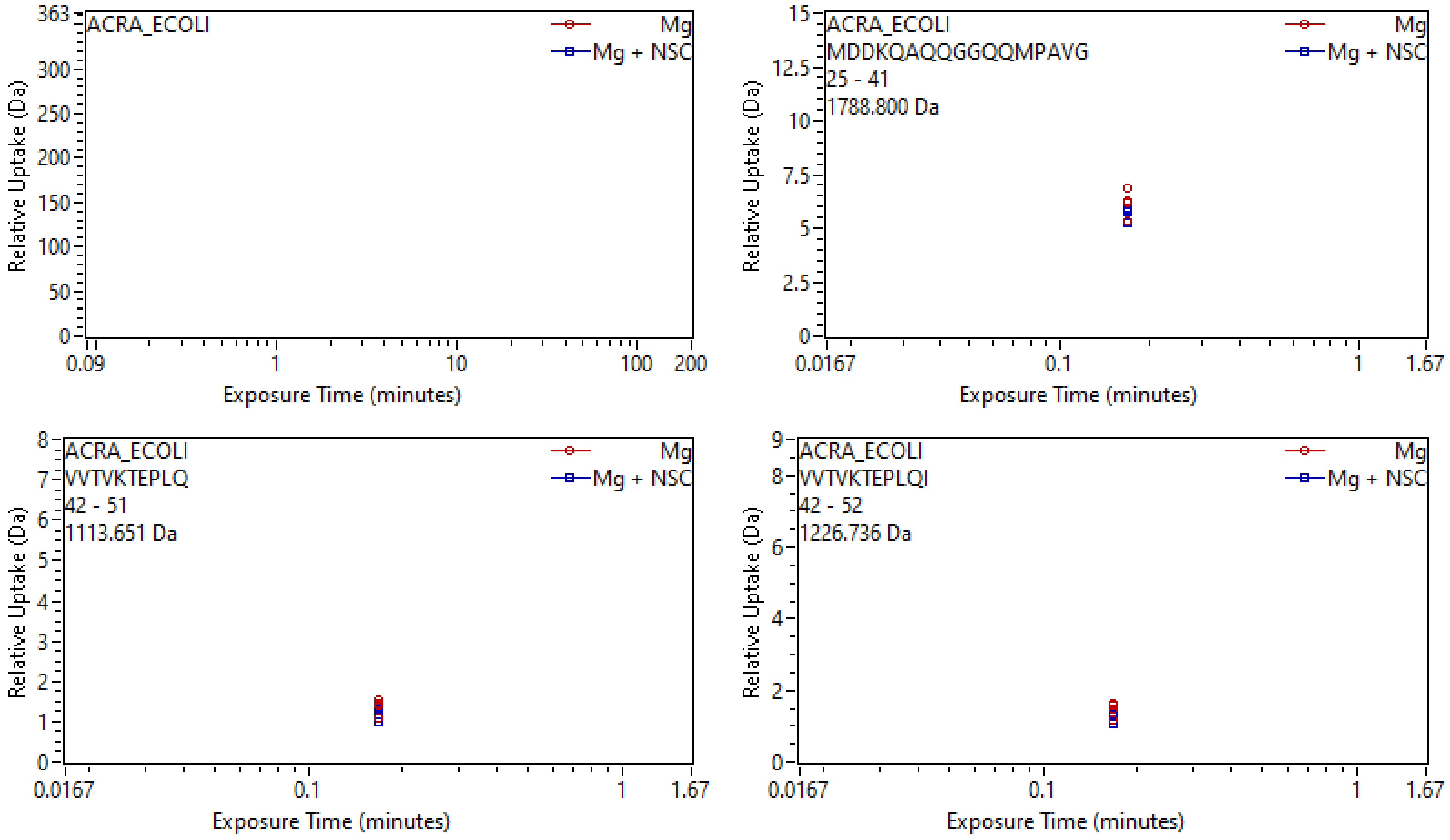
ACRA_ECOLI
Mg
Mg + NSC
Relative Uptake (Da)
Exposure Time (minutes)
0
50
100
150
200
250
300
363
0.09
1
10
100
200
ACRA_ECOLI
MDDKQAQQGGQQMPAVG
25 - 41
1788.800 Da
Mg
Mg + NSC
Relative Uptake (Da)
Exposure Time (minutes)
0
2.5
5
7.5
10
12.5
15
0.0167
0.1
1
1.67
ACRA_ECOLI
VVTVKTEPLQ
42 - 51
1113.651 Da
Mg
Mg + NSC
Relative Uptake (Da)
Exposure Time (minutes)
0
1
2
3
4
5
6
7
8
0.0167
0.1
1
1.67
ACRA_ECOLI
VVTVKTEPLQI
42 - 52
1226.736 Da
Mg
Mg + NSC
Relative Uptake (Da)
Exposure Time (minutes)
0
2
4
6
8
9
0.0167
0.1
1
1.67

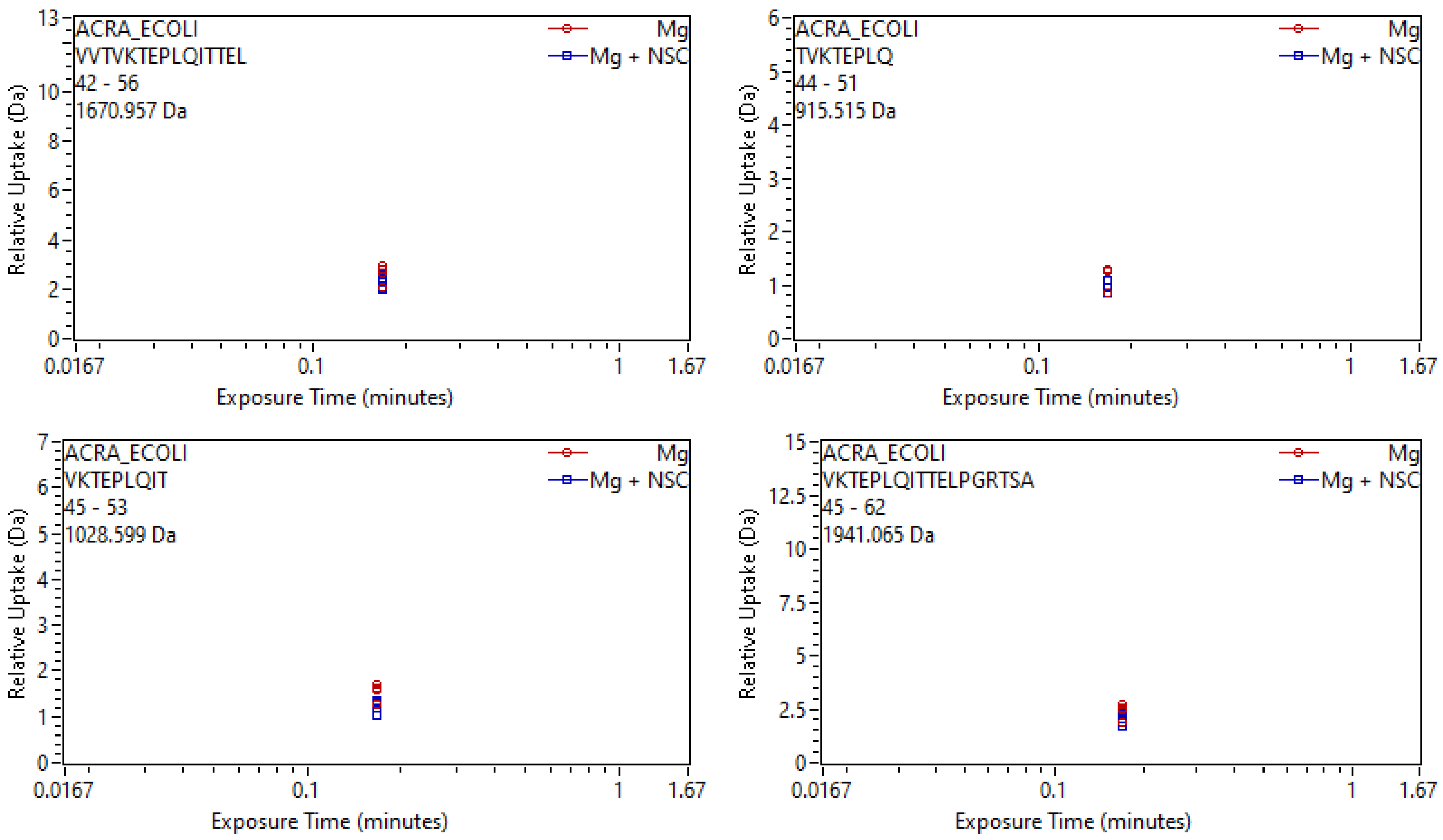
ACRA_ECOLI
VVTVKTEPLQITTEL
42 - 56
1670.957 Da
Mg
Mg + NSC
Relative Uptake (Da)
Exposure Time (minutes)
0.0167
0.1
1
1.67
0
2
4
6
8
10
13
ACRA_ECOLI
TVKTEPLQ
44 - 51
915.515 Da
Mg
Mg + NSC
Relative Uptake (Da)
Exposure Time (minutes)
ACRA_ECOLI
VKTEPLQIT
45 - 53
1028.599 Da
Mg
Mg + NSC
Relative Uptake (Da)
Exposure Time (minutes)
ACRA_ECOLI
VKTEPLQITTELPGRTSA
45 - 62
1941.065 Da
Mg
Mg + NSC
Relative Uptake (Da)
Exposure Time (minutes)
0
2.5
5
7.5
10
12.5
15

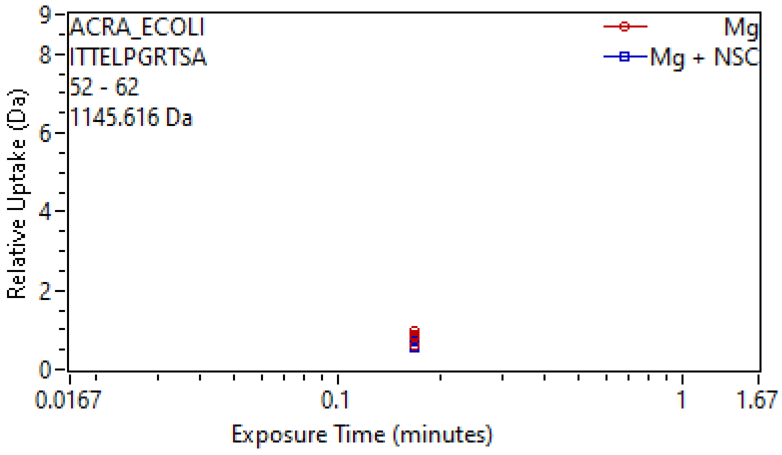
ACRA_ECOLI
ITTELPGRTSA
52 - 62
1145.616 Da
Mg
Mg + NSC
Relative Uptake (Da)
Exposure Time (minutes)
0.0167
0.1
1
1.67

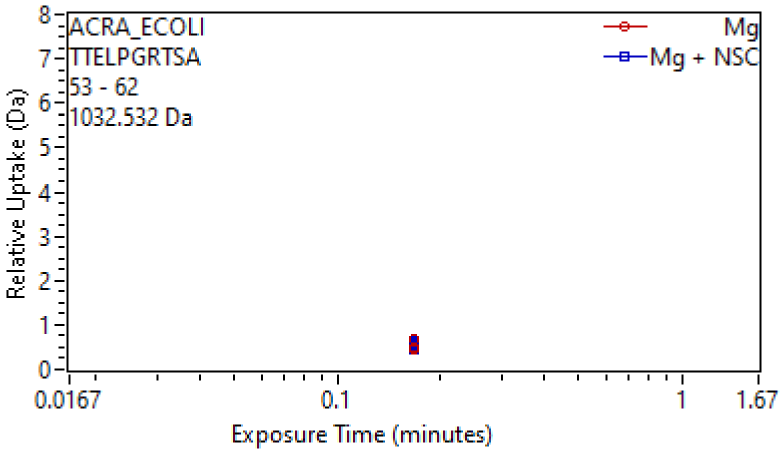
ACRA_ECOLI
TTELPGRTSA
53 - 62
1032.532 Da
Mg
Mg + NSC
Relative Uptake (Da)
Exposure Time (minutes)
0.0167
0.1
1
1.67

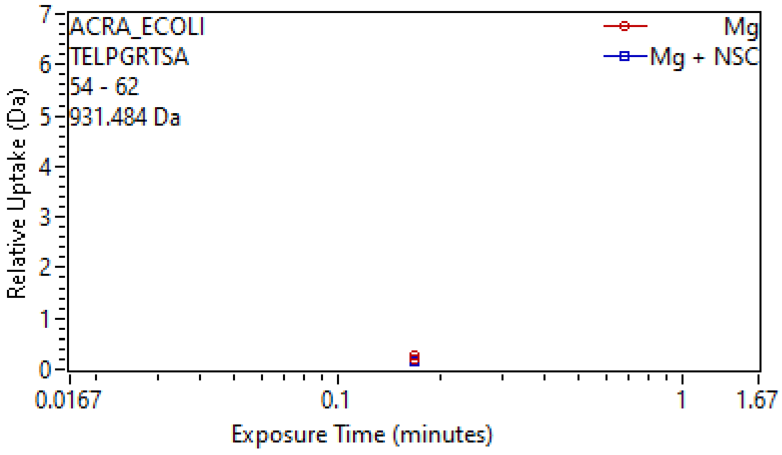
ACRA_ECOLI
TELPGRTSA
54 - 62
931.484 Da
Mg
Mg + NSC
Relative Uptake (Da)
Exposure Time (minutes)
0.0167
0.1
1
1.67

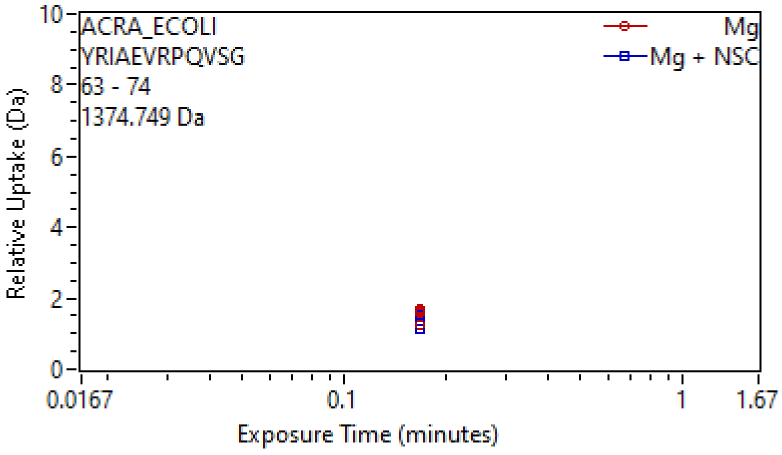
ACRA_ECOLI
YRIAEVRPQVSG
63 - 74
1374.749 Da
Mg
Mg + NSC
Relative Uptake (Da)
Exposure Time (minutes)
0.0167
0.1
1
1.67

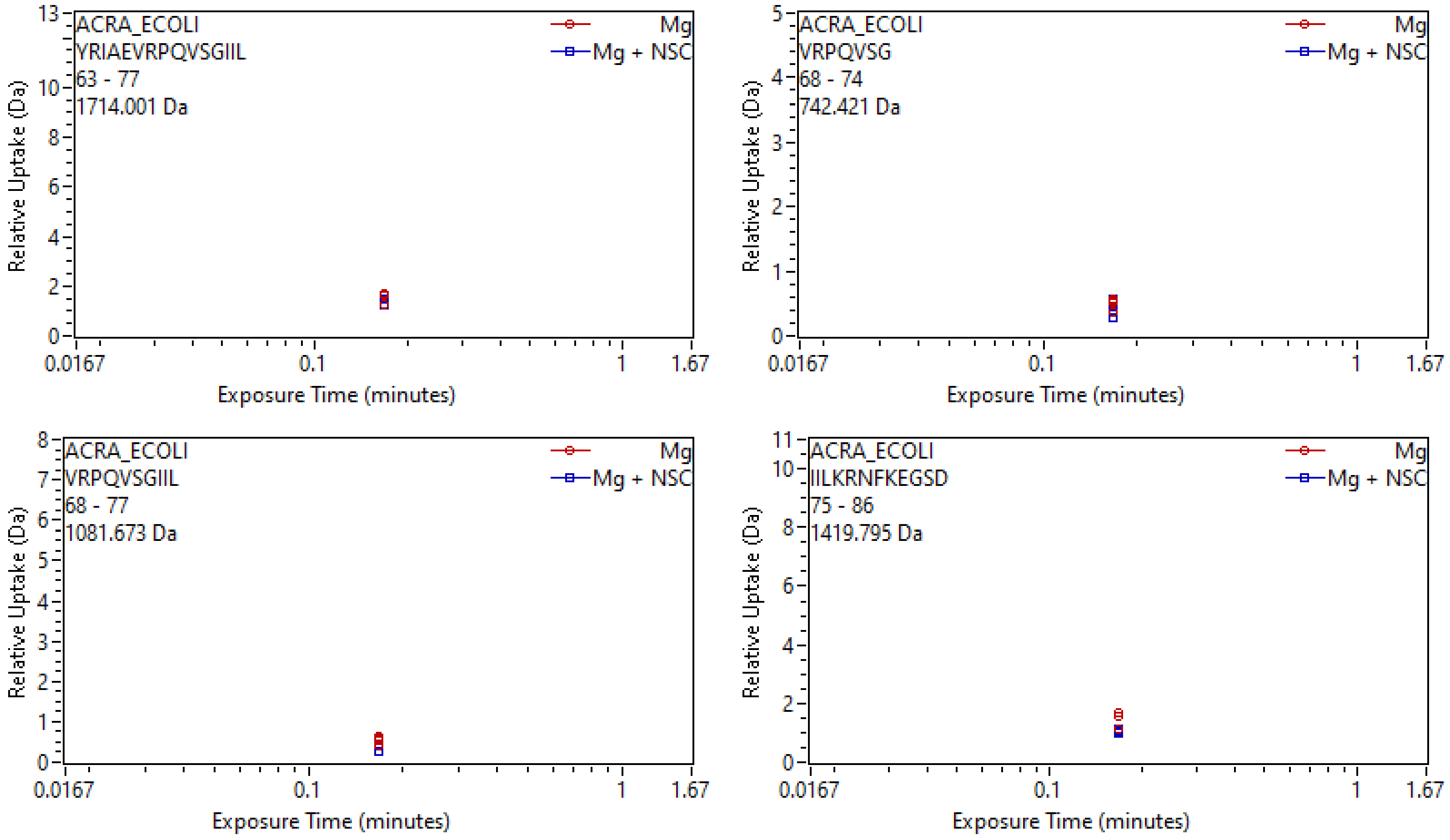
ACRA_ECOLI
YRIAEVRPQVSGIIL
63 - 77
1714.001 Da
Mg
Mg + NSC
Relative Uptake (Da)
Exposure Time (minutes)
0.0167
0.1
1
1.67
0
2
4
6
8
10
13
ACRA_ECOLI
VRPQVSG
68 - 74
742.421 Da
Mg
Mg + NSC
Relative Uptake (Da)
Exposure Time (minutes)
0.0167
0.1
1
1.67
0
1
2
3
4
5
ACRA_ECOLI
VRPQVSGIIL
68 - 77
1081.673 Da
Mg
Mg + NSC
Relative Uptake (Da)
Exposure Time (minutes)
0.0167
0.1
1
1.67
0
1
2
3
4
5
6
7
8
ACRA_ECOLI
IILKRNFKEGSD
75 - 86
1419.795 Da
Mg
Mg + NSC
Relative Uptake (Da)
Exposure Time (minutes)
0.0167
0.1
1
1.67
0
2
4
6
8
10
11

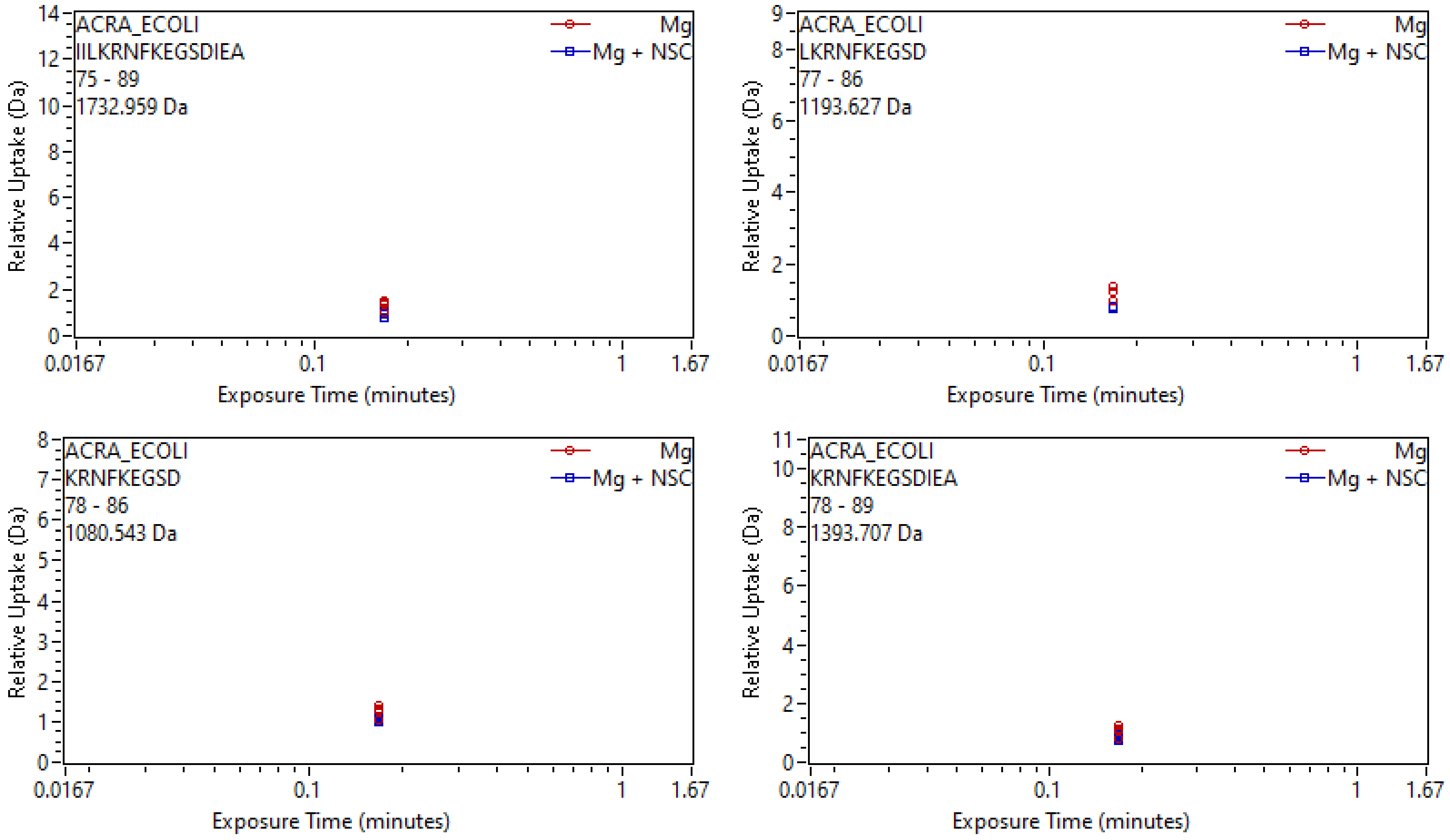

ACRA_ECOLI
IILKRNFKEGSDIEA
75 - 89
1732.959 Da
Mg
Mg + NSC
Relative Uptake (Da)
Exposure Time (minutes)
0.0167
0.1
1
1.67
ACRA_ECOLI
LKRNFKEGSD
77 - 86
1193.627 Da
ACRA_ECOLI
KRNFKEGSD
78 - 86
1080.543 Da
ACRA_ECOLI
KRNFKEGSDIEA
78 - 89
1393.707 Da

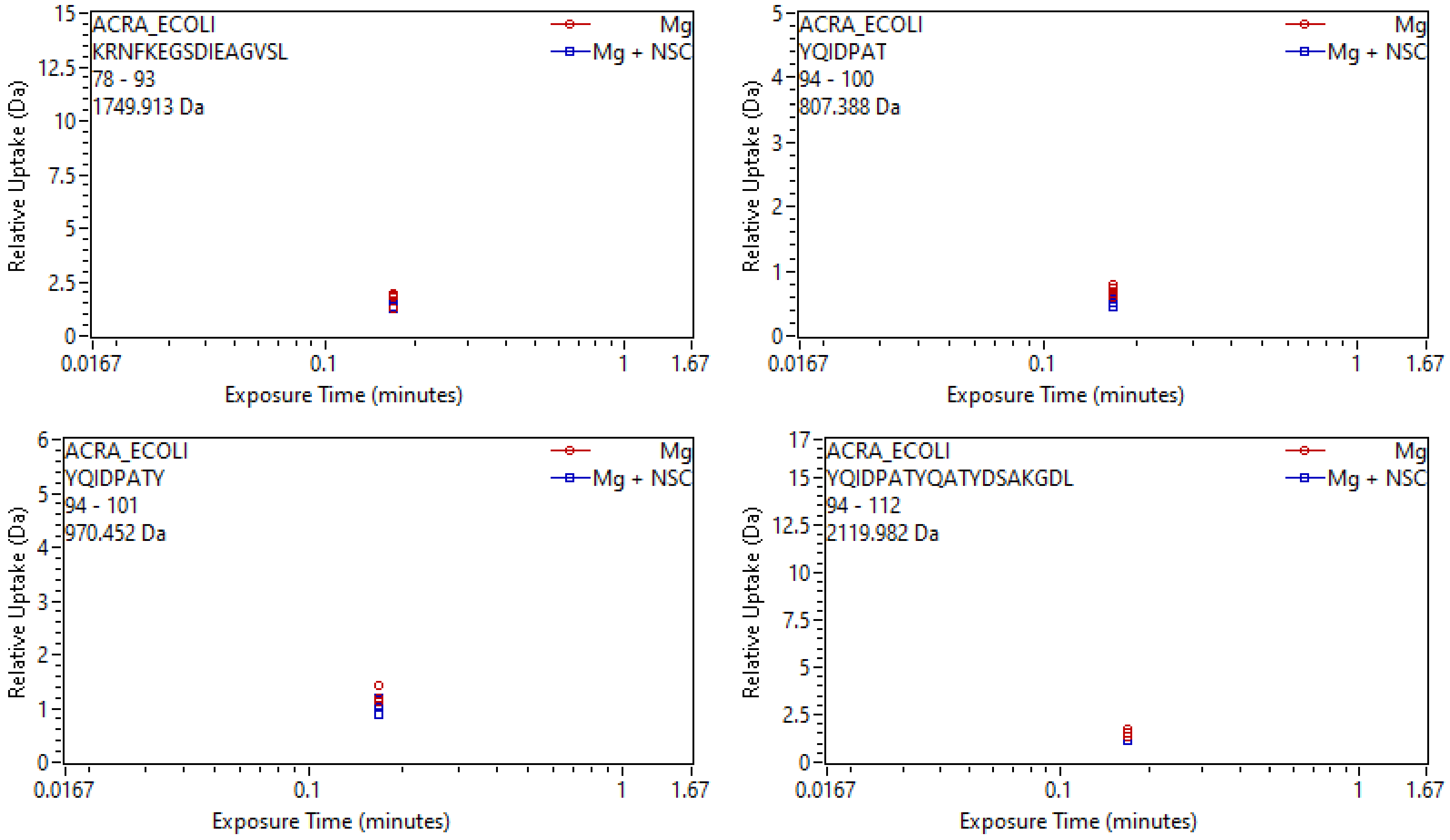
ACRA_ECOLI
KRNFKEGSDIEAGVSL
78 - 93
1749.913 Da
Mg
Mg + NSC
Relative Uptake (Da)
Exposure Time (minutes)
0.0167
0.1
1
1.67
ACRA_ECOLI
YQIDPAT
94 - 100
807.388 Da
ACRA_ECOLI
YQIDPATY
94 - 101
970.452 Da
ACRA_ECOLI
YQIDPATYQATYDSAKGDL
94 - 112
2119.982 Da

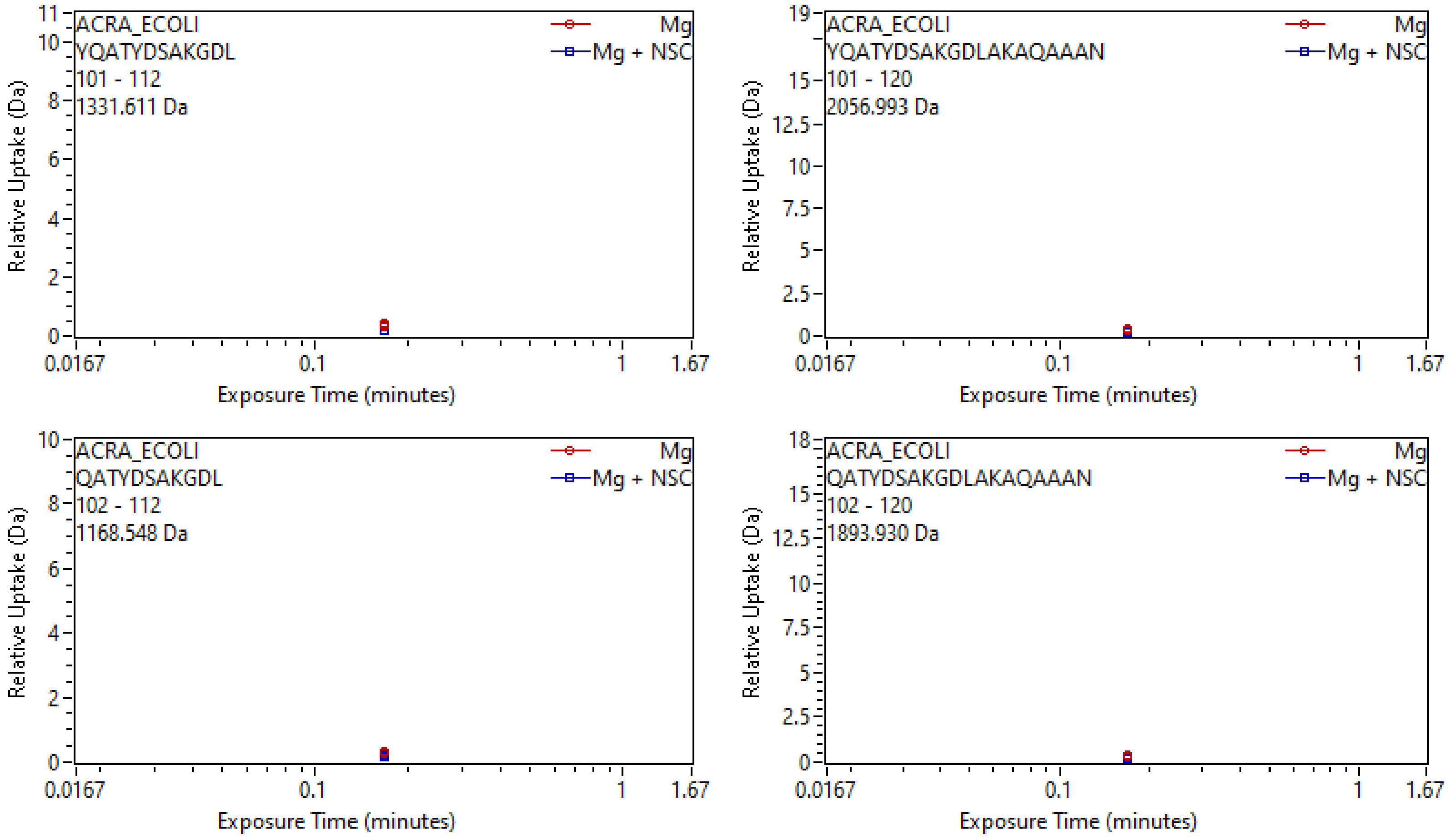
ACRA_ECOLI
YQATYDSAKGDL
101 - 112
1331.611 Da
Mg
Mg + NSC
Relative Uptake (Da)
Exposure Time (minutes)
0.0167
0.1
1
1.67
ACRA_ECOLI
YQATYDSAKGDLAKAQAAAN
101 - 120
2056.993 Da
Mg
Mg + NSC
Relative Uptake (Da)
Exposure Time (minutes)
ACRA_ECOLI
QATYDSAKGDL
102 - 112
1168.548 Da
Mg
Mg + NSC
Relative Uptake (Da)
Exposure Time (minutes)
ACRA_ECOLI
QATYDSAKGDLAKAQAAAN
102 - 120
1893.930 Da
Mg
Mg + NSC
Relative Uptake (Da)
Exposure Time (minutes)

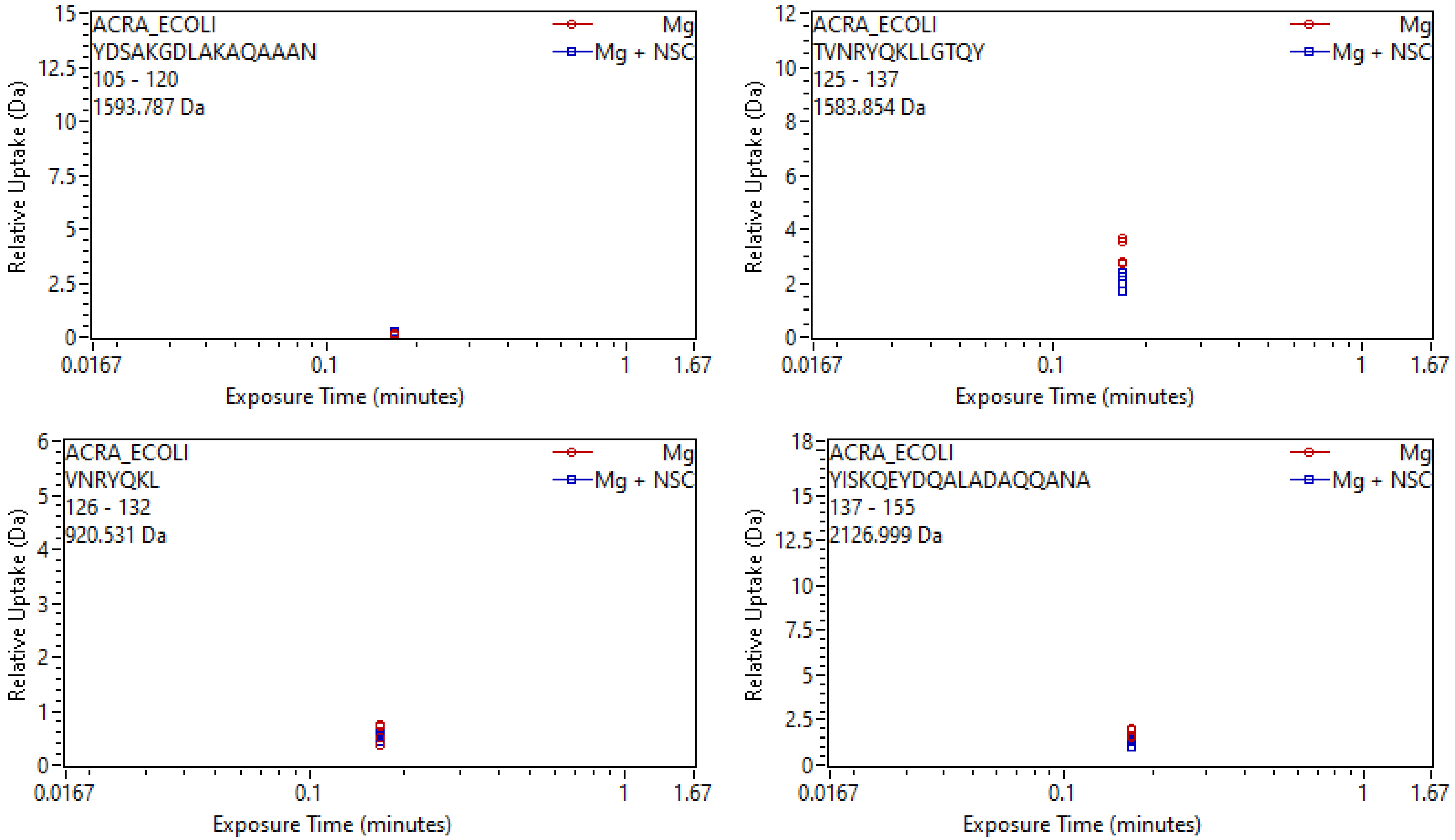
ACRA_ECOLI
YDSAKGDLAKAQAAAN
105 - 120
1593.787 Da
Mg
Mg + NSC
Relative Uptake (Da)
Exposure Time (minutes)
ACRA_ECOLI
TVNRYQKLLGTQY
125 - 137
1583.854 Da
Mg
Mg + NSC
Relative Uptake (Da)
Exposure Time (minutes)
ACRA_ECOLI
VNRYQKL
126 - 132
920.531 Da
Mg
Mg + NSC
Relative Uptake (Da)
Exposure Time (minutes)
ACRA_ECOLI
YISKQEYDQALADAQQANA
137 - 155
2126.999 Da
Mg
Mg + NSC
Relative Uptake (Da)
Exposure Time (minutes)

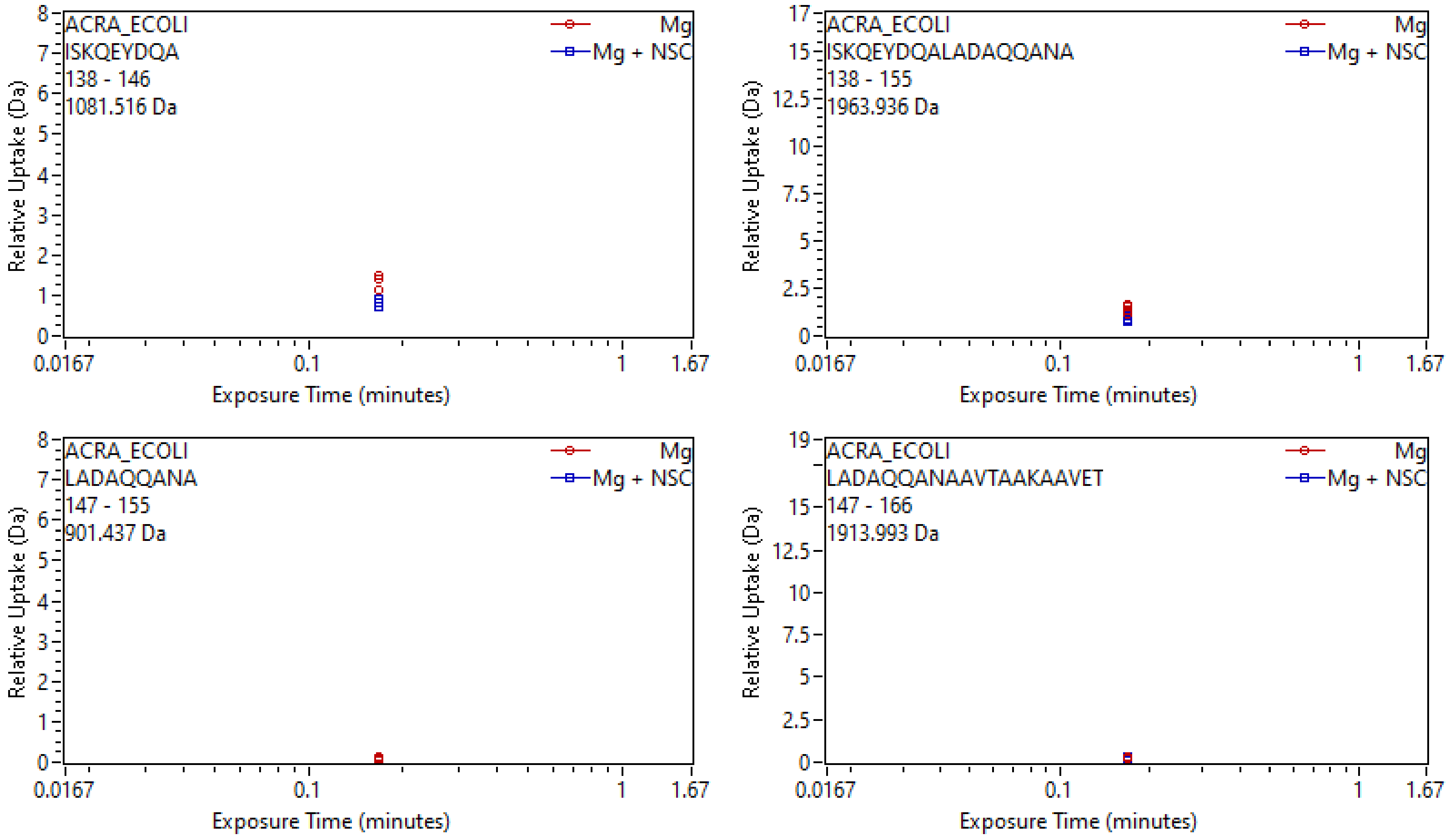
ACRA_ECOLI
ISKQEYDQA
138 - 146
1081.516 Da
Mg
Mg + NSC
Relative Uptake (Da)
Exposure Time (minutes)
ACRA_ECOLI
ISKQEYDQALADAQQANA
138 - 155
1963.936 Da
ACRA_ECOLI
LADAQQANA
147 - 155
901.437 Da
ACRA_ECOLI
LADAQQANAAVTAAKAAVET
147 - 166
1913.993 Da

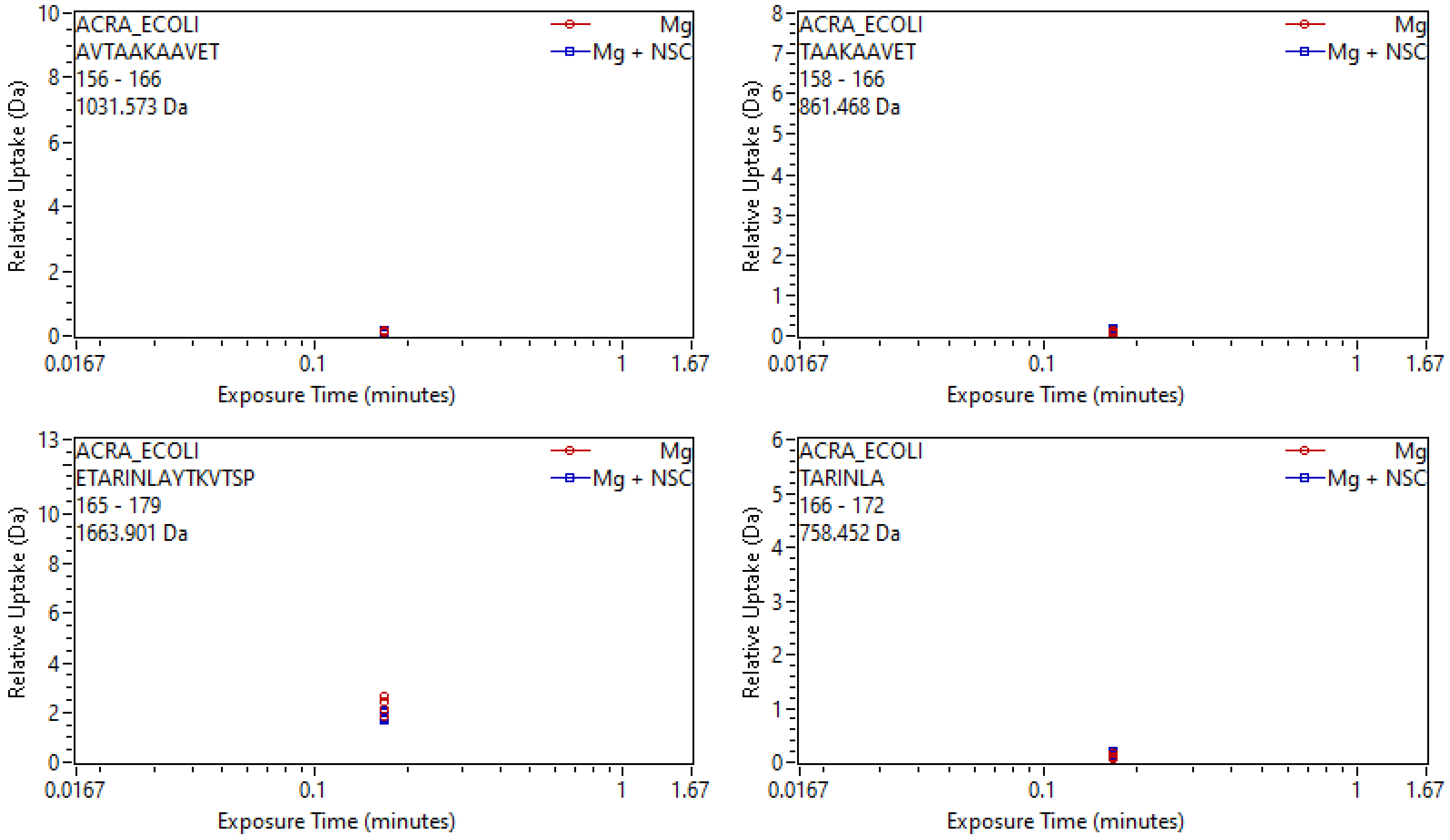
ACRA_ECOLI
AVTAAKAAVET
156 - 166
1031.573 Da
Mg
Mg + NSC
Relative Uptake (Da)
Exposure Time (minutes)
ACRA_ECOLI
TAAKAAVET
158 - 166
861.468 Da
Mg
Mg + NSC
Relative Uptake (Da)
Exposure Time (minutes)
ACRA_ECOLI
ETARINLAYTKVTSP
165 - 179
1663.901 Da
Mg
Mg + NSC
Relative Uptake (Da)
Exposure Time (minutes)
ACRA_ECOLI
TARINLA
166 - 172
758.452 Da
Mg
Mg + NSC
Relative Uptake (Da)
Exposure Time (minutes)

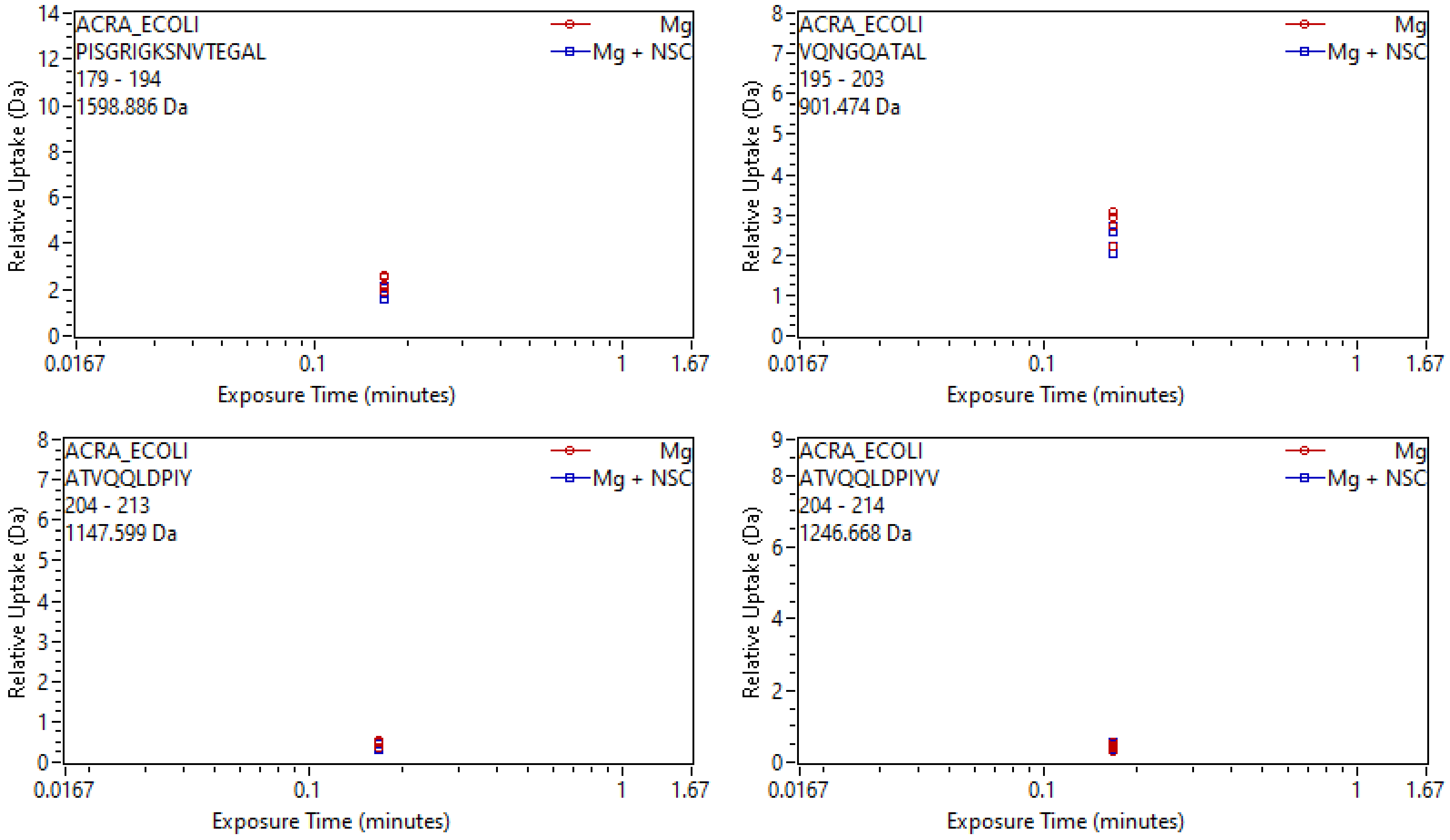
ACRA_ECOLI
PISGRIGKSNVTEGAL
179 - 194
1598.886 Da
Mg
Mg + NSC
Relative Uptake (Da)
Exposure Time (minutes)
ACRA_ECOLI
VQNGQATAL
195 - 203
901.474 Da
ACRA_ECOLI
ATVQQLDPIY
204 - 213
1147.599 Da
ACRA_ECOLI
ATVQQLDPIYV
204 - 214
1246.668 Da
0.0167
0.1
1
1.67

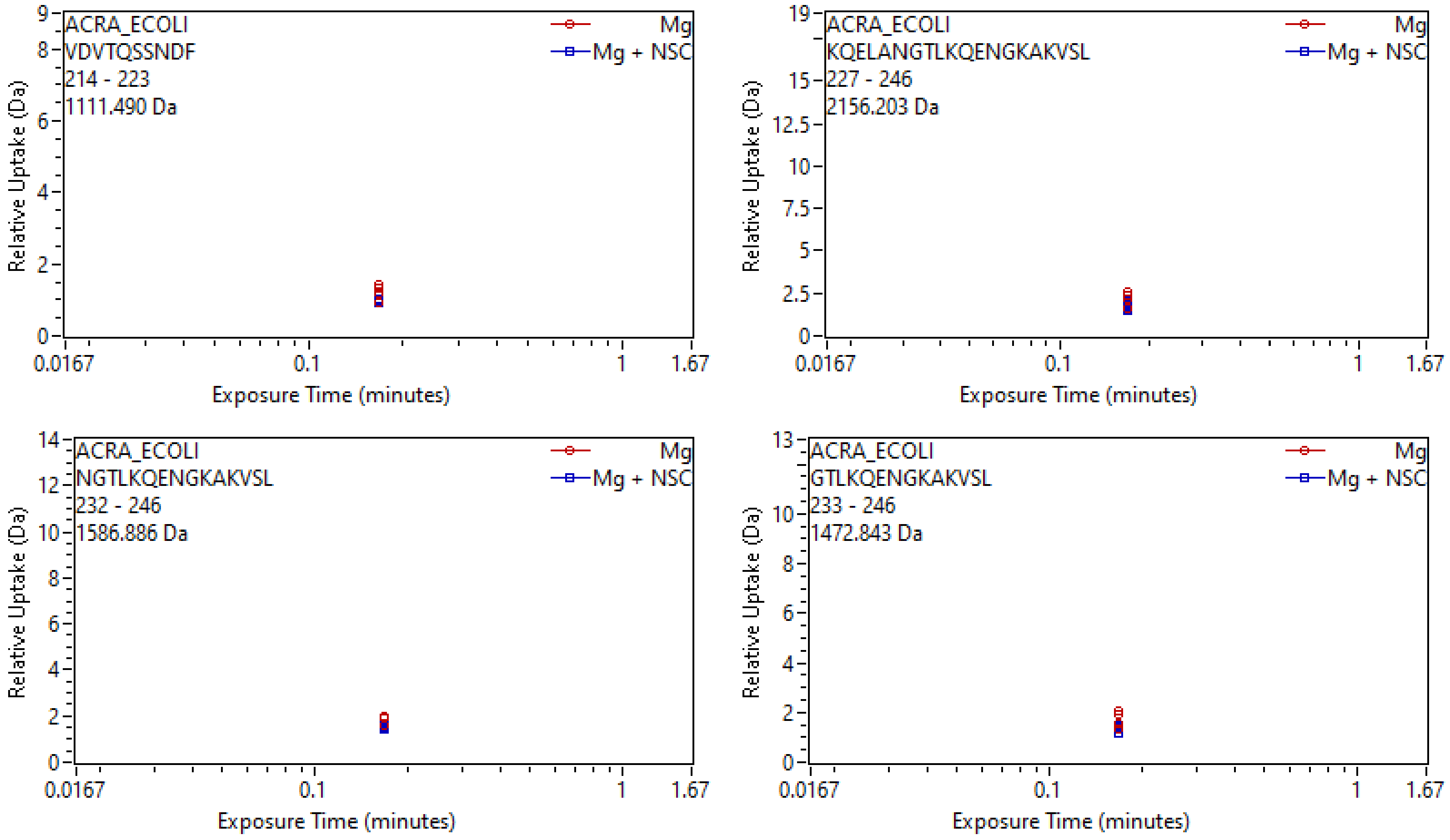
ACRA_ECOLI
VDVTQSSNDF
214 - 223
1111.490 Da
Mg
Mg + NSC
Relative Uptake (Da)
Exposure Time (minutes)
ACRA_ECOLI
KQELANGTLKQENGKAKVSL
227 - 246
2156.203 Da
ACRA_ECOLI
NGTLKQENGKAKVSL
232 - 246
1586.886 Da
ACRA_ECOLI
GTLKQENGKAKVSL
233 - 246
1472.843 Da

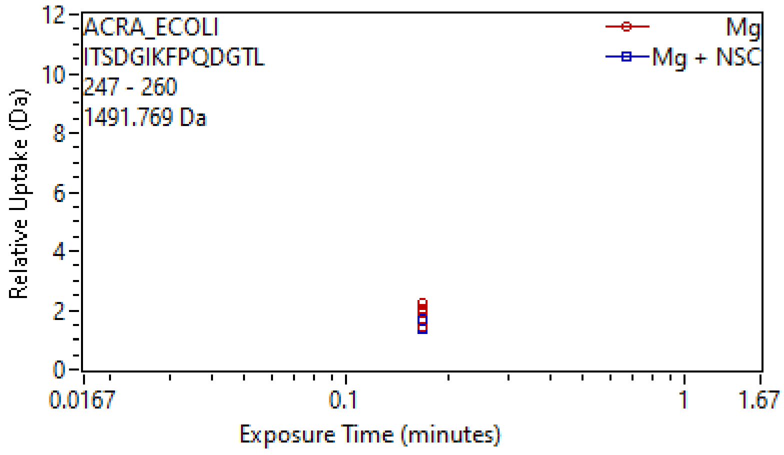
ACRA_ECOLI
ITSDGIKFPQDGTL
247 - 260
1491.769 Da
Mg
Mg + NSC
Relative Uptake (Da)
Exposure Time (minutes)
0.0167
0.1
1
1.67

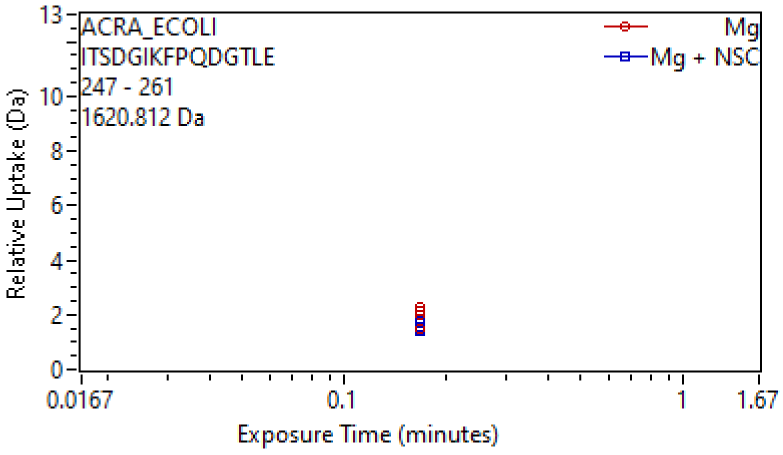
ACRA_ECOLI
ITSDGIKFPQDGTLE
247 - 261
1620.812 Da
Mg
Mg + NSC
Relative Uptake (Da)
Exposure Time (minutes)
0.0167
0.1
1
1.67

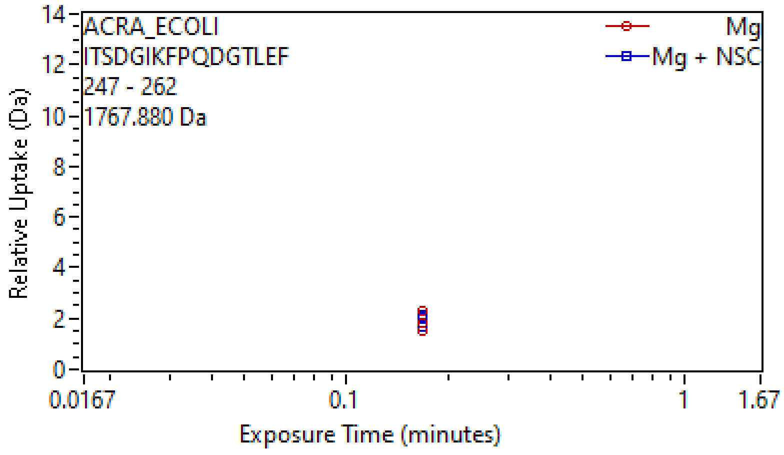
ACRA_ECOLI
ITSDGIKFPQDGTLEF
247 - 262
1767.880 Da
Mg
Mg + NSC
Relative Uptake (Da)
Exposure Time (minutes)
0.0167
0.1
1
1.67

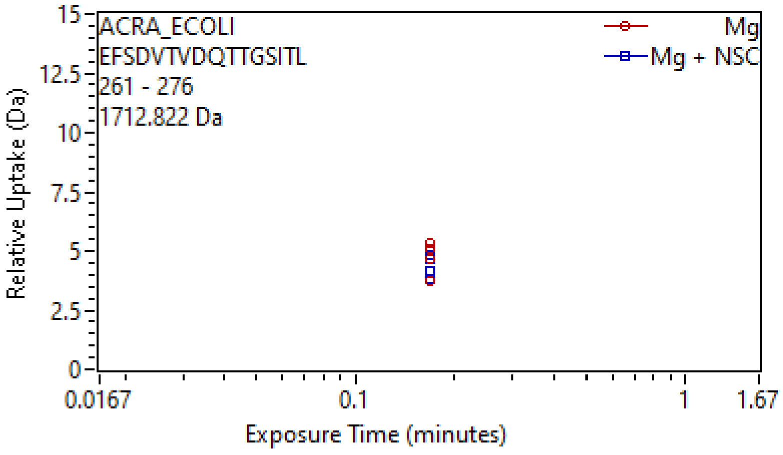
ACRA_ECOLI
EFSDVTVDQTTGSITL
261 - 276
1712.822 Da
Mg
Mg + NSC
Relative Uptake (Da)
Exposure Time (minutes)
0.0167
0.1
1
1.67

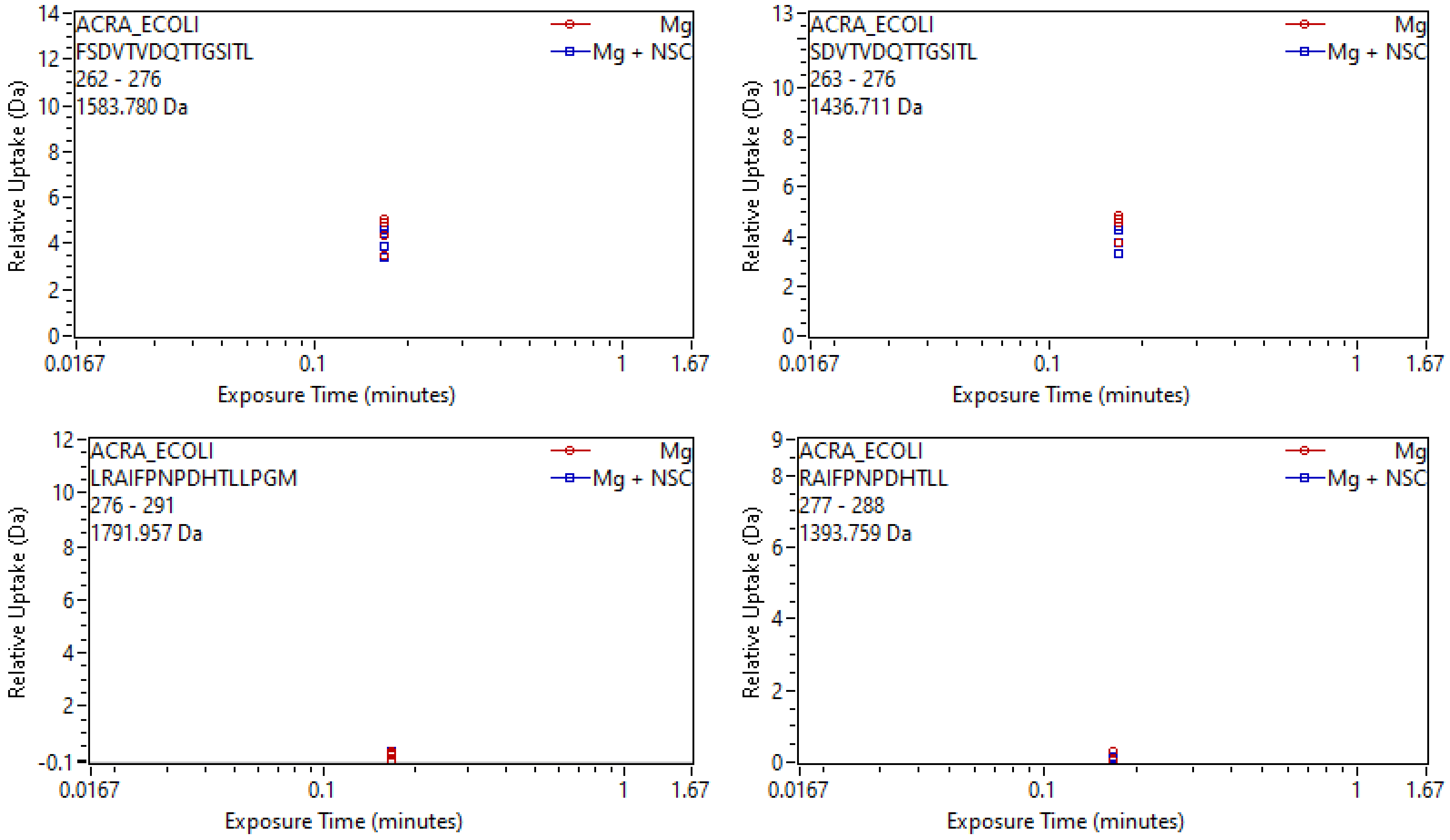
ACRA_ECOLI
FSDVTVDQTTGSITL
262 - 276
1583.780 Da
Mg
Mg + NSC
Relative Uptake (Da)
Exposure Time (minutes)
ACRA_ECOLI
SDVTVDQTTGSITL
263 - 276
1436.711 Da
ACRA_ECOLI
LRAIFPNPDHTLLPGM
276 - 291
1791.957 Da
ACRA_ECOLI
RAIFPNPDHTLL
277 - 288
1393.759 Da

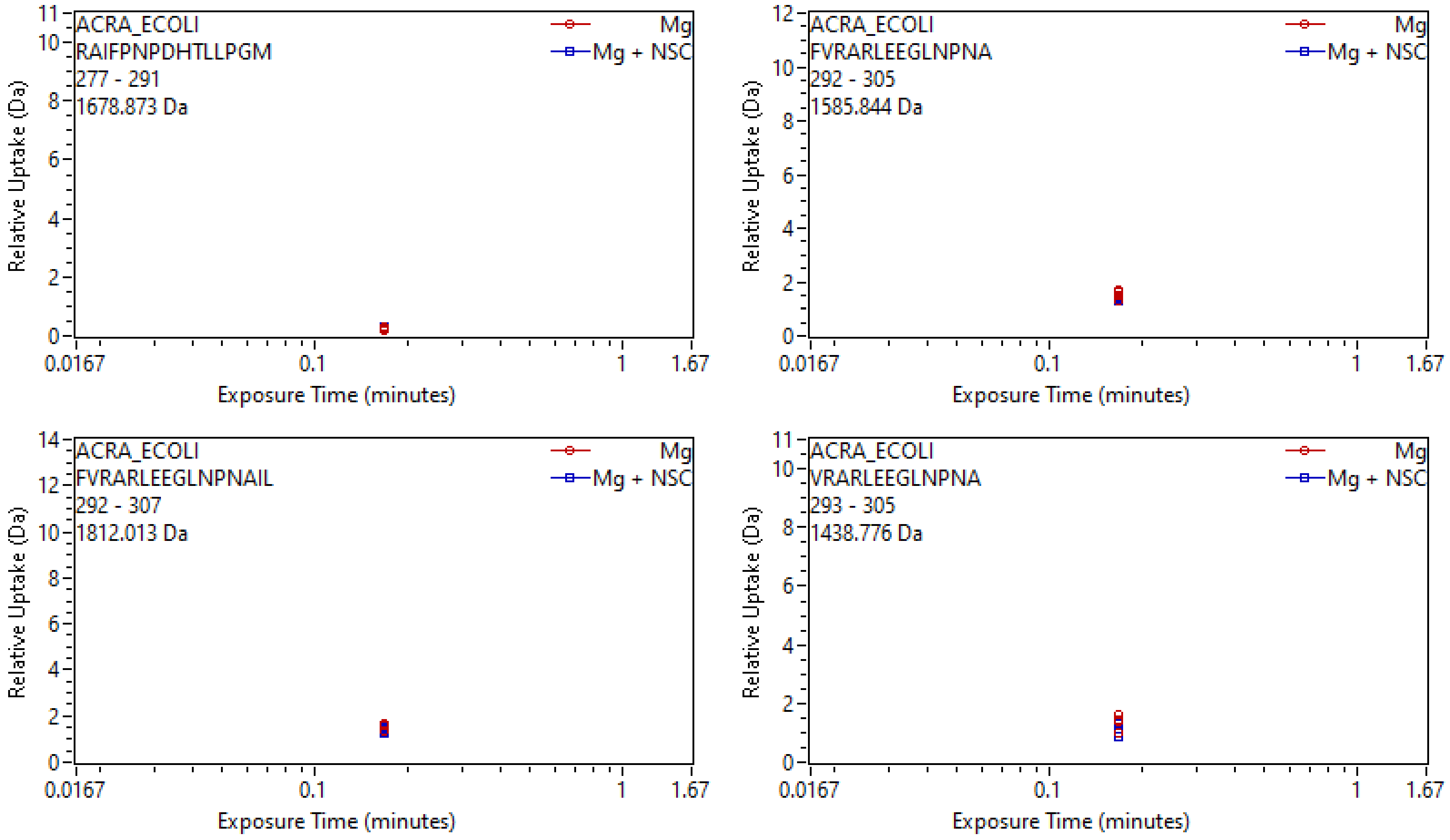
ACRA_ECOLI
RAIFPNPDHTLLPGM
277 - 291
1678.873 Da
Mg
Mg + NSC
Relative Uptake (Da)
Exposure Time (minutes)
ACRA_ECOLI
FVRARLEEGLNPNA
292 - 305
1585.844 Da
Mg
Mg + NSC
Relative Uptake (Da)
Exposure Time (minutes)
ACRA_ECOLI
FVRARLEEGLNPNAIL
292 - 307
1812.013 Da
Mg
Mg + NSC
Relative Uptake (Da)
Exposure Time (minutes)
ACRA_ECOLI
VRARLEEGLNPNA
293 - 305
1438.776 Da
Mg
Mg + NSC
Relative Uptake (Da)
Exposure Time (minutes)

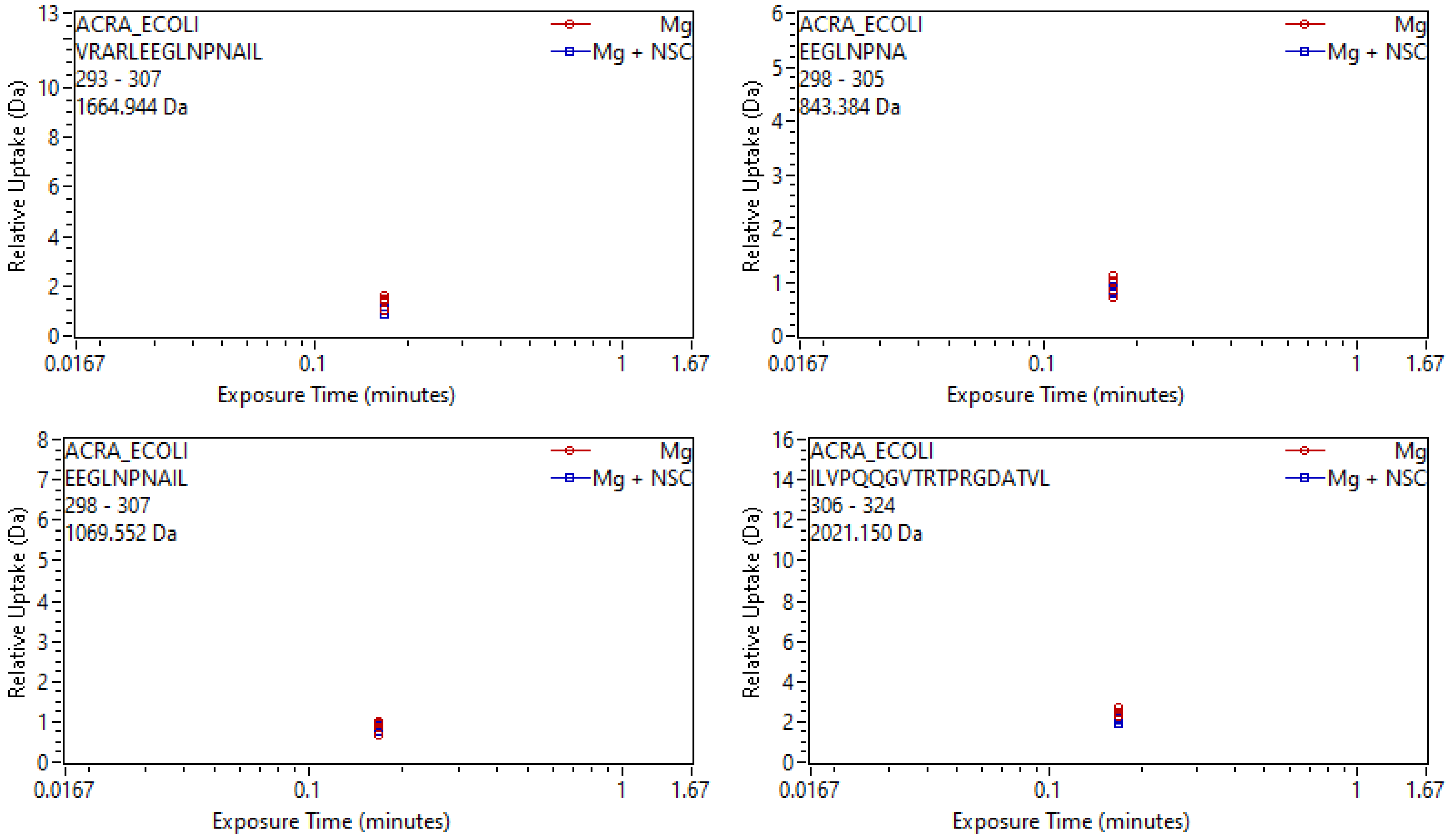
ACRA_ECOLI
VRARLEEGLNPNAIL
293 - 307
1664.944 Da
Mg
Mg + NSC
Relative Uptake (Da)
Exposure Time (minutes)
ACRA_ECOLI
EEGLNPNA
298 - 305
843.384 Da
Mg
Mg + NSC
Relative Uptake (Da)
Exposure Time (minutes)
ACRA_ECOLI
EEGLNPNAIL
298 - 307
1069.552 Da
Mg
Mg + NSC
Relative Uptake (Da)
Exposure Time (minutes)
ACRA_ECOLI
ILVPQQGVTRTPRGDATVL
306 - 324
2021.150 Da
Mg
Mg + NSC
Relative Uptake (Da)
Exposure Time (minutes)

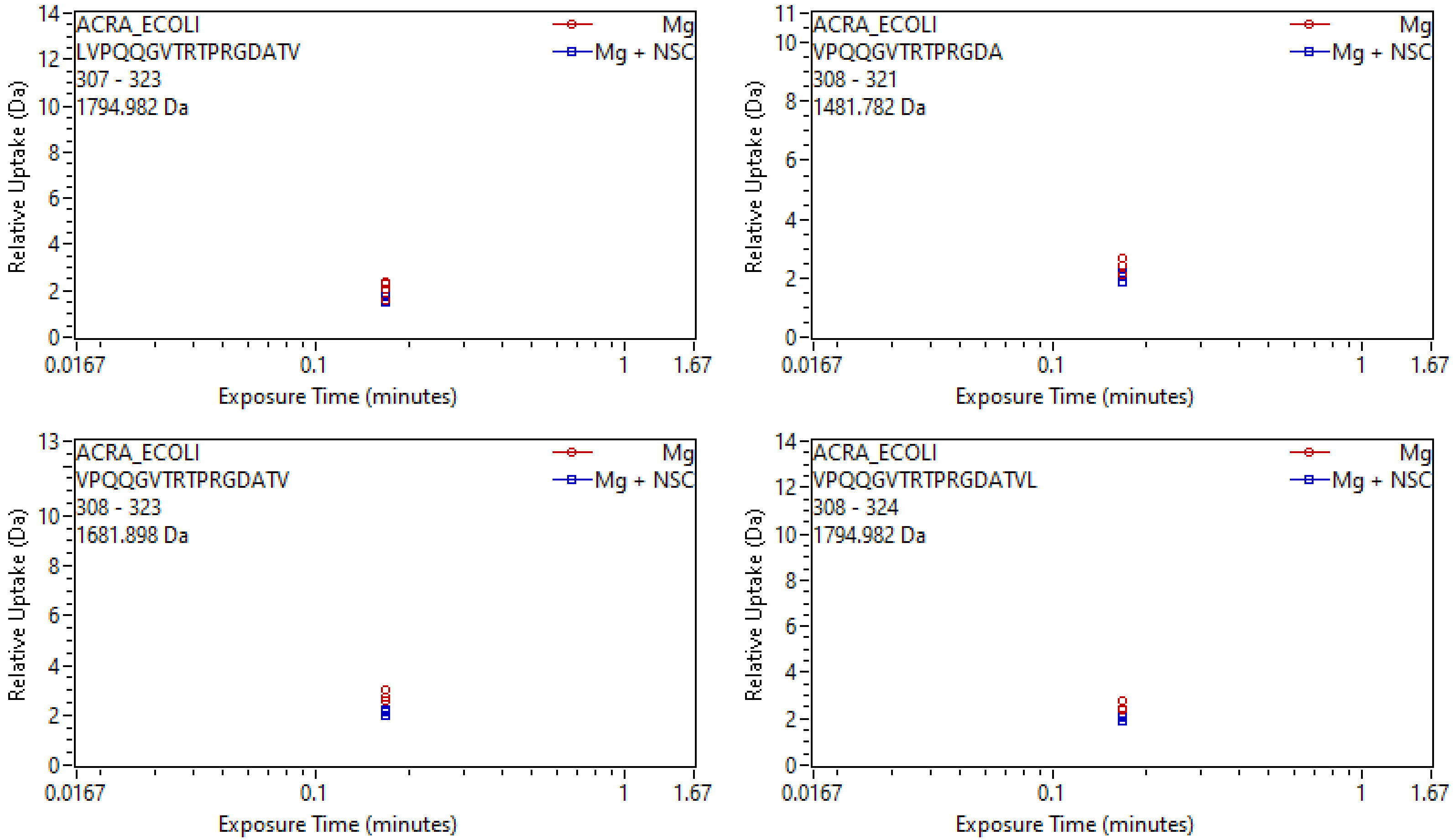
ACRA_ECOLI
LVPQQGVTRTPRGDATV
307 - 323
1794.982 Da
Mg
Mg + NSC
Relative Uptake (Da)
Exposure Time (minutes)
0.0167
0.1
1
1.67
ACRA_ECOLI
VPQQGVTRTPRGDA
308 - 321
1481.782 Da
Mg
Mg + NSC
Relative Uptake (Da)
Exposure Time (minutes)
ACRA_ECOLI
VPQQGVTRTPRGDATV
308 - 323
1681.898 Da
Mg
Mg + NSC
Relative Uptake (Da)
Exposure Time (minutes)
ACRA_ECOLI
VPQQGVTRTPRGDATVL
308 - 324
1794.982 Da
Mg
Mg + NSC
Relative Uptake (Da)
Exposure Time (minutes)

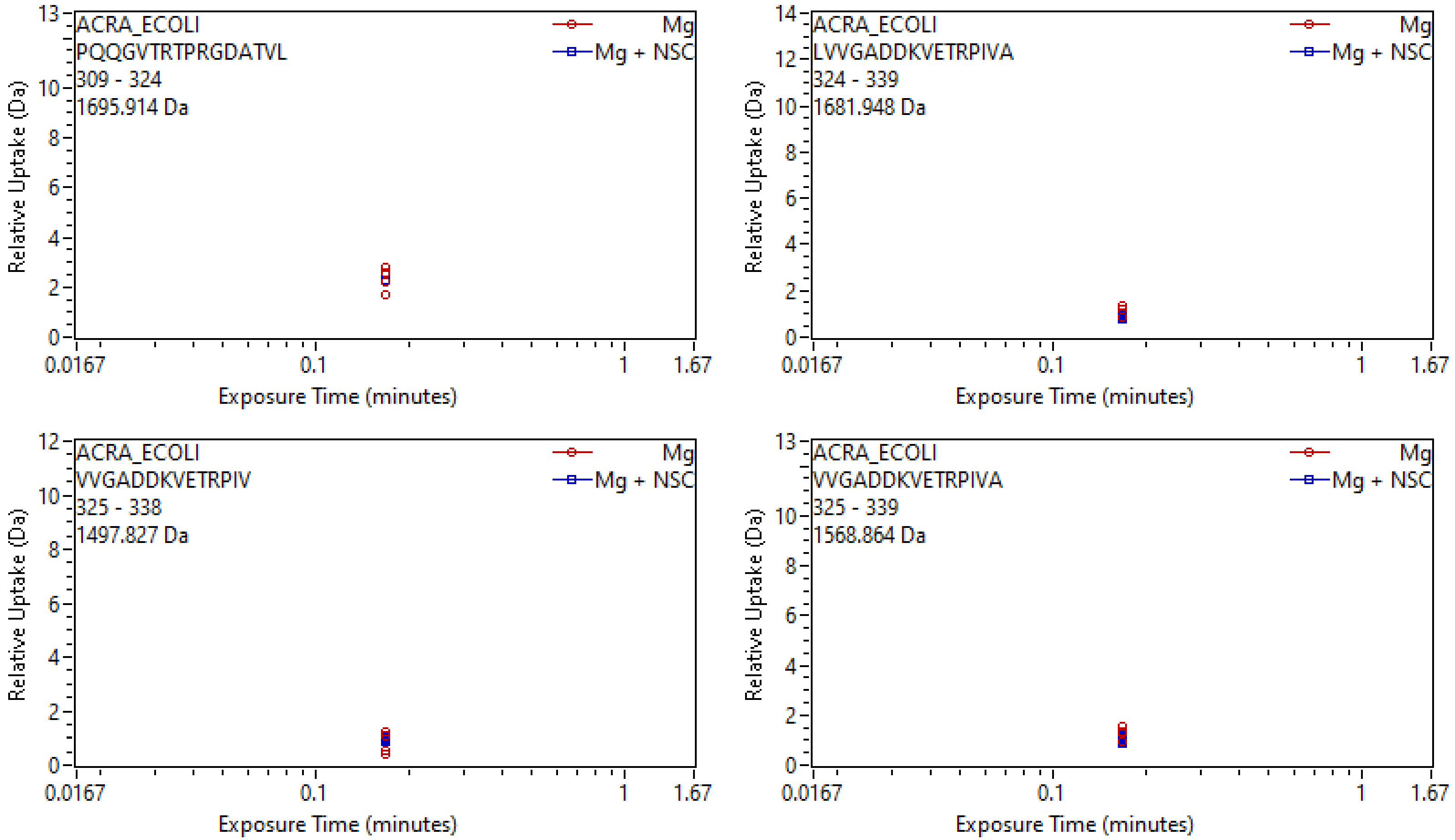
ACRA_ECOLI
PQQGVTRTPRGDATVL
309 - 324
1695.914 Da
Mg
Mg + NSC
Relative Uptake (Da)
Exposure Time (minutes)
ACRA_ECOLI
LVVGADDKVETRPIVA
324 - 339
1681.948 Da
Mg
Mg + NSC
Relative Uptake (Da)
Exposure Time (minutes)
ACRA_ECOLI
VVGADDKVETRPIV
325 - 338
1497.827 Da
Mg
Mg + NSC
Relative Uptake (Da)
Exposure Time (minutes)
ACRA_ECOLI
VVGADDKVETRPIVA
325 - 339
1568.864 Da
Mg
Mg + NSC
Relative Uptake (Da)
Exposure Time (minutes)

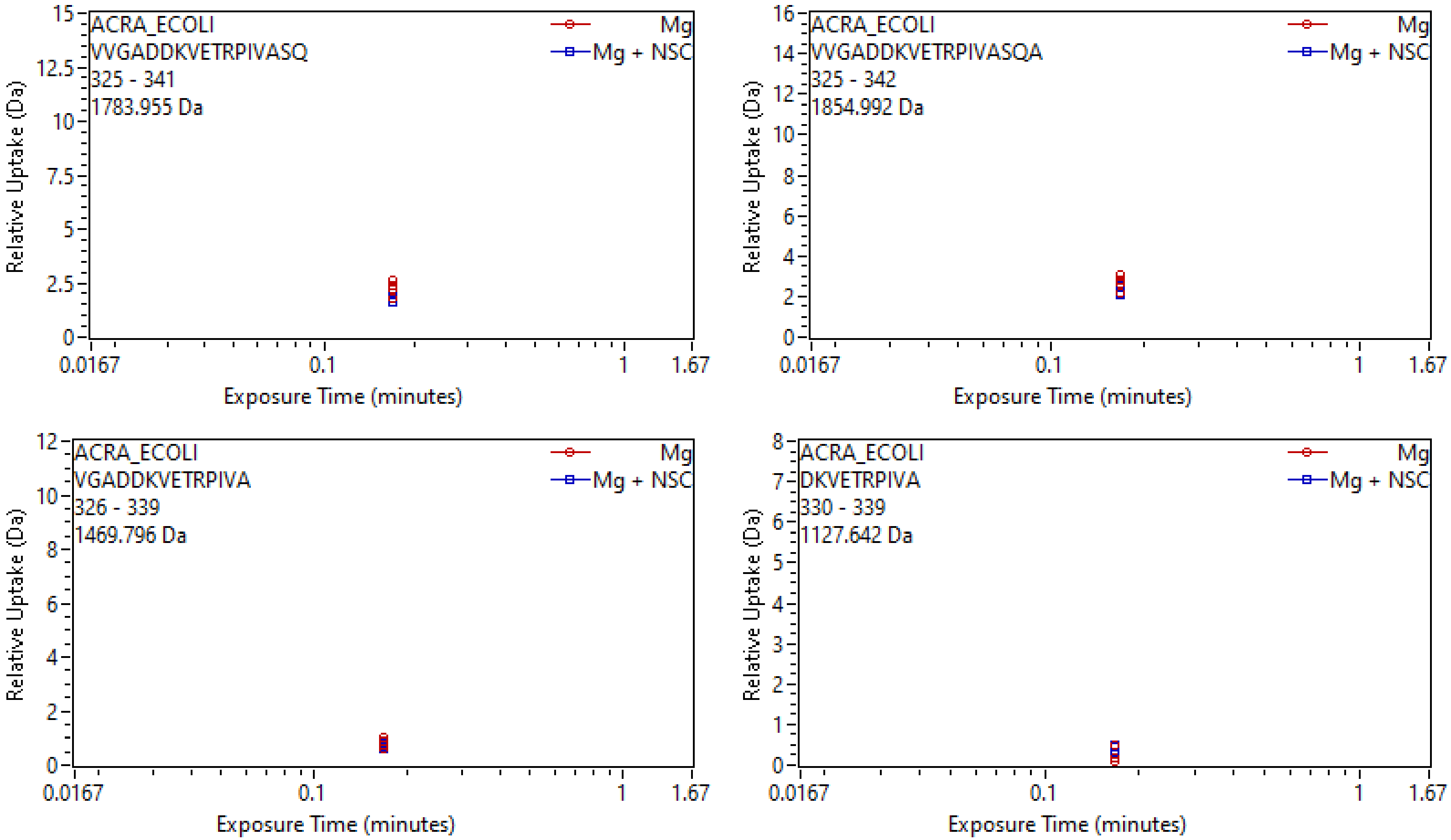
ACRA_ECOLI
VVGADDKVETRPIVASQ
325 - 341
1783.955 Da
Mg
Mg + NSC
Relative Uptake (Da)
15
12.5
10
7.5
5
2.5
0
0.0167
0.1
1
1.67
Exposure Time (minutes)
ACRA_ECOLI
VVGADDKVETRPIVASQA
325 - 342
1854.992 Da
Mg
Mg + NSC
Relative Uptake (Da)
16
14
12
10
8
6
4
2
0
0.0167
0.1
1
1.67
Exposure Time (minutes)
ACRA_ECOLI
VGADDKVETRPIVA
326 - 339
1469.796 Da
Mg
Mg + NSC
Relative Uptake (Da)
12
10
8
6
4
2
0
0.0167
0.1
1
1.67
Exposure Time (minutes)
ACRA_ECOLI
DKVETRPIVA
330 - 339
1127.642 Da
Mg
Mg + NSC
Relative Uptake (Da)
8
7
6
5
4
3
2
1
0
0.0167
0.1
1
1.67
Exposure Time (minutes)

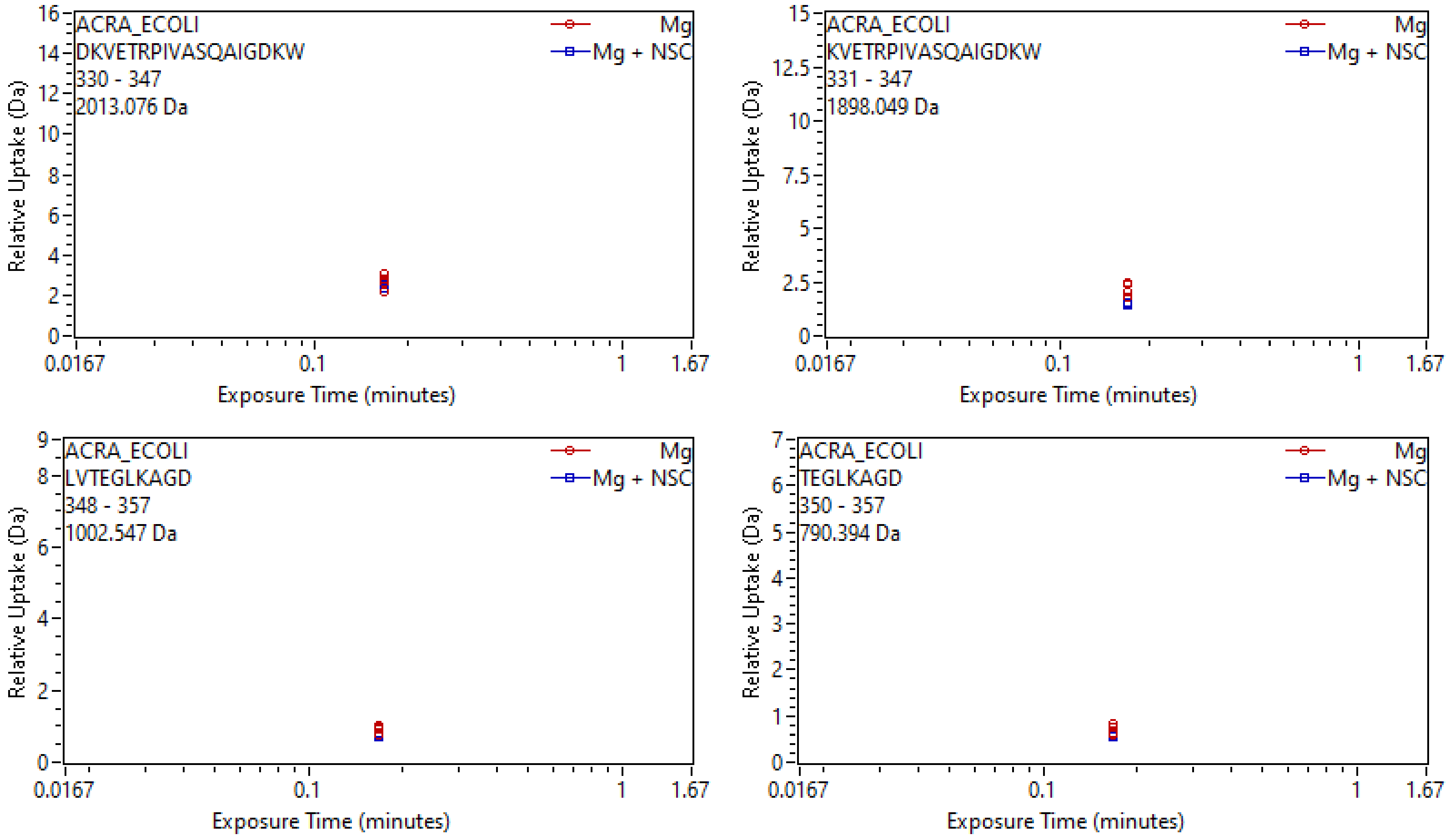
ACRA_ECOLI
DKVETRPIVASQAIGDKW
330 - 347
2013.076 Da
Mg
Mg + NSC
Relative Uptake (Da)
Exposure Time (minutes)
0.0167
0.1
1
1.67
0
2
4
6
8
10
12
14
16
ACRA_ECOLI
KVETRPIVASQAIGDKW
331 - 347
1898.049 Da
Mg
Mg + NSC
Relative Uptake (Da)
Exposure Time (minutes)
0
2.5
5
7.5
10
12.5
15
ACRA_ECOLI
LVTEGLKAGD
348 - 357
1002.547 Da
Mg
Mg + NSC
Relative Uptake (Da)
Exposure Time (minutes)
0
2
4
6
8
9
ACRA_ECOLI
TEGLKAGD
350 - 357
790.394 Da
Mg
Mg + NSC
Relative Uptake (Da)
Exposure Time (minutes)
0
1
2
3
4
5
6
7

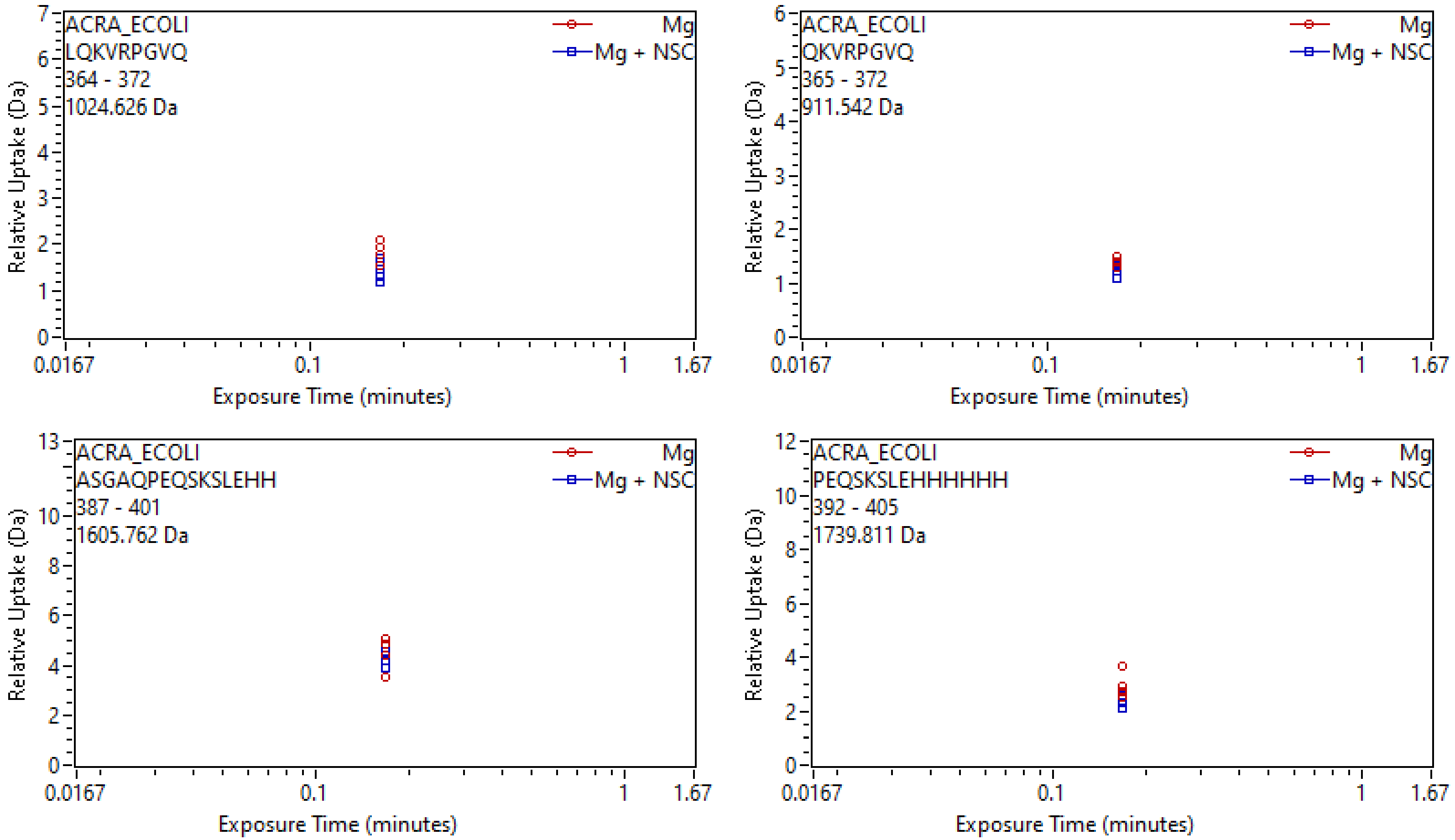
ACRA_ECOLI
LQKVRPGVQ
364 - 372
1024.626 Da
Mg
Mg + NSC
Relative Uptake (Da)
Exposure Time (minutes)
0.0167
0.1
1
1.67
ACRA_ECOLI
QKVRPGVQ
365 - 372
911.542 Da
Mg
Mg + NSC
Relative Uptake (Da)
Exposure Time (minutes)
ACRA_ECOLI
ASGAQPEQSKSLEHH
387 - 401
1605.762 Da
Mg
Mg + NSC
Relative Uptake (Da)
Exposure Time (minutes)
ACRA_ECOLI
PEQSKSLEHHHHHH
392 - 405
1739.811 Da
Mg
Mg + NSC
Relative Uptake (Da)
Exposure Time (minutes)
